# Analysis of stress-induced small proteins in *Escherichia coli* reveals that YoaI mediates cross-talk between distinct signaling systems

**DOI:** 10.1101/2024.09.13.612970

**Authors:** Sangeevan Vellappan, Junhong Sun, John Favate, Pranavi Jagadeesan, Debbie Cerda, Premal Shah, Srujana S. Yadavalli

## Abstract

Bacterial small proteins (≤ 50 amino acids) are an emerging class of regulators that modulate the activity of signaling networks that enable bacterial adaptation to stress. The *Escherichia coli* genome encodes at least 150 small proteins, most of which are functionally uncharacterized. We identified and characterized 17 small proteins induced in *E*. *coli* during magnesium (Mg^2+^) starvation using ribosome profiling, RNA sequencing, and transcriptional reporter assays. Several of these were transcriptionally activated by the PhoQ-PhoP two-component signaling system, which is crucial for Mg^2+^ homeostasis. Deletion or overexpression of some of these small proteins led to growth defects and changes in cell size under low-Mg^2+^ conditions, indicating physiological roles in stress adaptation. The small transmembrane protein YoaI, which was transcriptionally induced by the phosphate-responsive PhoR-PhoB signaling pathway, increased in abundance under Mg^2+^ limitation independently of *yoaI* transcription or PhoQ-PhoP signaling. YoaI activated a third signaling system, EnvZ-OmpR, which mediates responses to osmotic stress. Overall, this study establishes an initial framework for understanding how small proteins contribute to bacterial stress adaptation by facilitating cross-talk between different signaling systems. Our results suggest these proteins play broader roles in coordinating stress responses, reflecting the interconnected nature of cellular stress networks rather than strictly compartmentalized pathways responding to specific stressors.

## Introduction

Living organisms sense and respond to environmental stressors through a wide variety of gene regulatory mechanisms (*1*). In bacteria, signal transduction is primarily carried out by two-component signaling systems (*2–4*). Gene expression regulation can occur at various stages, including transcription, post-transcription, translation, and post-translation (*5–8*). Although some of these mechanisms involving regulators such as transcription factors and small RNAs have been investigated in depth (*9–11*), other regulatory pathways essential for stress adaptation remain less well understood. Small proteins (≤50 amino acids in prokaryotes and ≤100 amino acids in eukaryotes), encoded by authentic small open reading frames, are emerging as key regulators of stress responses (*12–14*), highlighting a need for further investigation into their roles and mechanisms of action.

Advances in bioinformatics, gene expression studies, and ribosome profiling have identified thousands of previously unannotated small open reading frames (sORFs) in all kingdoms of life, including humans (*13*, *15–20*). In *Escherichia coli,* more than 150 small proteins have been documented (*13*, *15*). There has been substantial progress in identifying small proteins, but the functions of most of these proteins remain to be determined (*12*, *15*). Small proteins are typically nonessential, at least under standard laboratory growth conditions, and their requirement is likely limited to contexts in which the activity of a larger target needs fine-tuning. In support of this idea, some small proteins accumulate in higher amounts in a specific growth phase, medium, or stress condition (*21*, *22*). Magnesium (Mg^2+^) is an important cofactor affecting the catalysis and stability of numerous proteins and RNAs involved in vital cellular processes, including translation (*23*). Given the central role of Mg^2+^, its deprivation poses a major stress to cells. The signal transduction and physiological response to low Mg^2+^ is fairly well understood, which involves the activation of the master regulator PhoQP, a two-component signaling system in gammaproteobacteria (*23–25*). In addition to Mg^2+^ limitation, this signaling system responds to other stimuli such as cationic antimicrobial peptides, mildly acidic pH, and high osmolarity (*24*, *26–28*).

So far, three small proteins are found to mediate stress responses to Mg^2+^ limitation in bacteria. MgrB, a 47-aa transmembrane protein, is transcriptionally induced by the response regulator PhoP upon Mg^2+^ starvation and negatively feeds back on the PhoQP signaling system by inhibiting the sensor kinase PhoQ (*29*, *30*). The absence of MgrB during Mg^2+^ starvation leads to hyperactivation of the PhoQP pathway, resulting in cell division inhibition and filamentation in *E. coli* (*31*). In addition to its role in Mg^2+^ stress, loss of MgrB in *E. coli* increases tolerance to trimethoprim, an antibiotic commonly used to treat bacterial infections (*32*). In clinical isolates of *Klebsiella pneumoniae*, disruption of MgrB-mediated inhibition of PhoQ also leads to acquired colistin resistance (*33*). A second small transmembrane protein, MgtS, is also induced under low-Mg^2+^ stress and is important for Mg^2+^ homeostasis (*34*, *35*). MgtS protects the Mg^2+^ transporter MgtA from degradation when Mg^2+^ is limited (*34*). PmrR is a third putative small membrane-associated protein expressed under low-Mg^2+^ stress in *Salmonella enterica* (*36*, *37*). PmrR is activated by the PmrAB two-component system (*36*, *38*, *39*) in a PhoP-dependent manner and inhibits the lipopolysaccharide modification enzyme LpxT.

Here we asked what subset of the ∼150 small proteins in *E. coli* were induced under Mg^2+^ limitation. We hypothesized that, like MgrB, MgtS, and PmrR, other small proteins may play regulatory roles as part of this stress response. We leveraged the translation initiation site profiling method, Ribo-RET (*40*, *41*) to identify small proteins induced by low-Mg^2+^ stress, revealing a set of 17 proteins representing a substantial proportion (∼11%) of the documented small proteins in *E. coli*. Using RNA-Seq and transcriptional reporter assays, we investigated the transcriptional regulation of these stress-induced small proteins, shedding light on the genomic regions responsible for their expression and discerning any regulatory influence from the PhoQP system. Additionally, using epitope-tagging, microscopy, and biochemical analyses, we examined the localization and the overexpression and loss-of-function phenotypes of these small proteins. Through these experiments, we uncovered how the small transmembrane protein YoaI, transcriptionally activated by the PhoRB signaling pathway (*42*), displays increased protein abundance under Mg^2+^ stress and activates another well-studied two-component system EnvZ-OmpR (*43*) in *E. coli*.

## Results

### Identification of low-Mg^2+^ stress–induced small proteins in *E. coli* using translation initiation profiling

To identify small proteins that accumulate under low-Mg^2+^ stress, we adapted the translation initiation profiling method called Ribo-RET (Fig. 1A) – a modified ribosome profiling approach utilizing translation initiation inhibitors for the identification of small proteins in bacteria (*44*). Retapamulin (RET) is an antimicrobial compound that can stall ribosomes at translation start sites to help identify previously unannotated open reading frames (*40*, *41*). In this study, we prepared Ribo-RET libraries of wild-type (WT) *E. coli* cells grown in media with or without Mg^2+^ to examine small proteins induced in response to Mg^2+^ starvation.

**Fig. 1.**
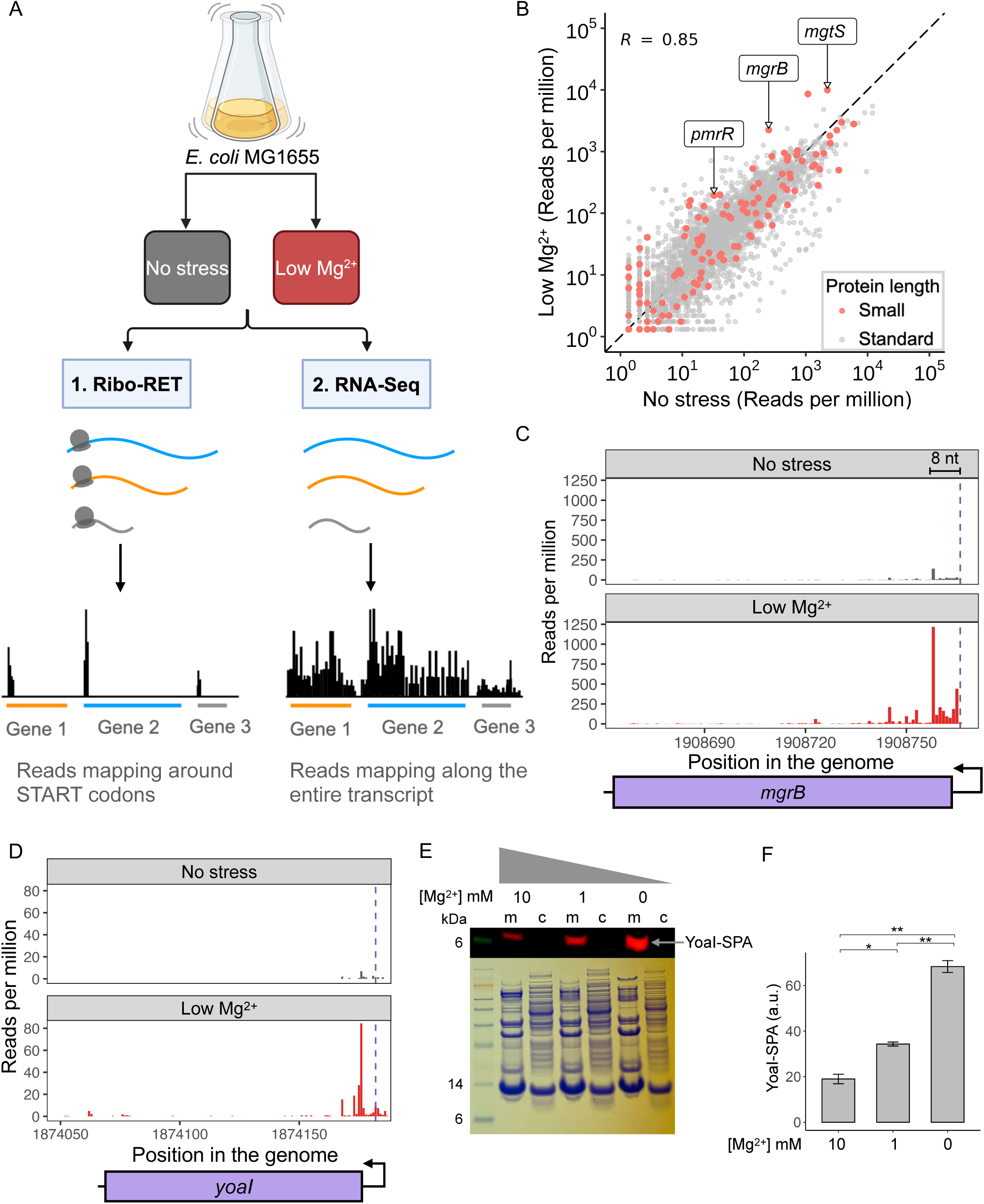
Identification of small proteins induced under low-Mg^2+^ stress in *E. coli* using translation-initiation profiling (Ribo-RET). (**A**) Schematic diagram showing the Ribo-RET and RNA-Seq experimental setup used to identify small proteins induced by low-Mg^2+^ stress. (**B**) Scatterplot showing the correlation between the Ribo-RET reads mapping to the annotated translation start sites of transcripts expressed under low-Mg^2+^ stress and no stress. The red dots represent transcripts encoding small proteins of ≤ 50 amino acids in length, and the gray dots represent those encoding proteins > 50 amino acids long. Transcripts for the small proteins MgrB, MgtS, and PmrR, previously identified as induced under Mg^2+^ starvation, are highlighted. Pearson’s coefficient, r= 0.85, n = 1. (**C and D**) Ribo-RET data for the small proteins MgrB (C) and YoaI (D) induced under low-Mg^2+^ stress. The dashed line indicates the translation start site, and the 8 nucleotide (nt) annotation indicates the distance between the highest ribosome density peak and the first nucleotide of the start codon. (**E**) Validation of YoaI production in media containing MgSO_4_ at the indicated concentration using a strain containing *yoaI-SPA* genomic fusion (GSO317). The SPA tag consists of 3x FLAG epitope sequence. Membrane (m) and cytoplasmic (c) fractions were analyzed by western blotting using an antibody specific for FLAG and by Coomassie Brilliant Blue staining. n = 3 biological replicates per group. **(F)** Quantification of the YoaI-SPA protein amounts from the western blot in (E). Bars represent the mean ± standard error of the mean from n = 3 biological replicates per group. P-values were calculated using a t-test as indicated; *P < 0.05, **P ≤ 0.01.

In our Ribo-RET data, we observed an increase in reads mapping to a specific distance of the ribosomal P-site codon from the 3’ end of the reads in our libraries, ∼6-10 nucleotides (nt) downstream of the expected start codon (fig. S1), similar to the pattern observed in a previous *E*. *coli* translation initiation profiling study using the translation initiation inhibitor Onc112 (*41*). However, this distance differs from the ∼15 nt peak reported in a previous Ribo-RET analysis in *E*. *coli* (*40*).

We think that this discrepancy in the ribosome footprint distances is likely due to the difference in the library preparation methods and independent of the translation inhibitor used. Our Ribo-RET method and the Ribo-Seq with Onc112 (*41*) both used sucrose cushion treatment to sediment ribosomes before nuclease digestion and fractionation, a sedimentation step thought to remove tRNAs from the ribosomal A-site (*41*). The previous Ribo-RET protocol did not include this sedimentation step using a sucrose cushion (*40*). We calculated ribosome density at annotated translation initiation sites using reads mapped from 4 to 20 nt downstream of the first nucleotide in the start codon, using a broad window of 16 nt to capture all relevant footprint sizes. In our analysis, the ribosome footprint sizes ranged from 16-24 nt (fig. S1), as anticipated for the bacterial systems (*45*).

The PhoQP signaling system is stimulated under Mg^2+^ limitation (*46*) (fig. S2, A and B). As expected, the read counts at the translation start site for PhoP and PhoQ were more than 4-fold and 10-fold higher, respectively, under stress conditions compared to no stress. Three small proteins known to be induced under low-Mg^2+^ stress – MgrB, MgtS, and PmrR, showed a 9-fold, 5-fold, and 5-fold increase in the reads mapping to the translation start sites, respectively, under stress vs. no stress (Fig. 1, B and C, fig. S2C). Overall, in our Ribo-RET data, we saw a moderate correlation between the reads mapping to the start sites of all proteins expressed under these two conditions (Fig. 1B), which is indicative of the anticipated changes in the translation initiation of proteins during stress response.

We detected 379 ORFs with ≥3-fold increase in read counts (Reads per Million, RPM) under stress, several of which were sORFs encoding proteins <50 aa in length (table S4). To obtain a short list of sORFs for further analyses, we used a strict expression cut-off (RPM values ≥ 5). After applying this filter, we obtained ∼19 sORFs, of which 17 were selected for detailed characterization. Two candidates, *rhoL* and *yodE*, were not included for further analysis. *rhoL* encodes the *rho* operon leader peptide, which controls the expression of the gene encoding Rho transcription termination factor, which in turn affects at least 20-30% of the global gene expression (*47*). As a result, any perturbation of this gene is likely to lead to large systemic changes, making it harder to characterize its specific role in the Mg^2+^ stress response. *yodE* is a sORF encoded within the previously identified ribosome-associated small RNA RyeG (*48*) and encodes a toxic peptide, which has been a challenge to clone to date by our group as well as others (*48*). We proceeded with a set of 17 small proteins for further analysis: MgtS, MgrB, PmrR, MgtT, YmiC, YmiA, YoaI, YobF, YddY, YriB, YadX, YkgS, YriA, YdgU, YqhI, YadW, and DinQ (Table 1).

**Table 1.**
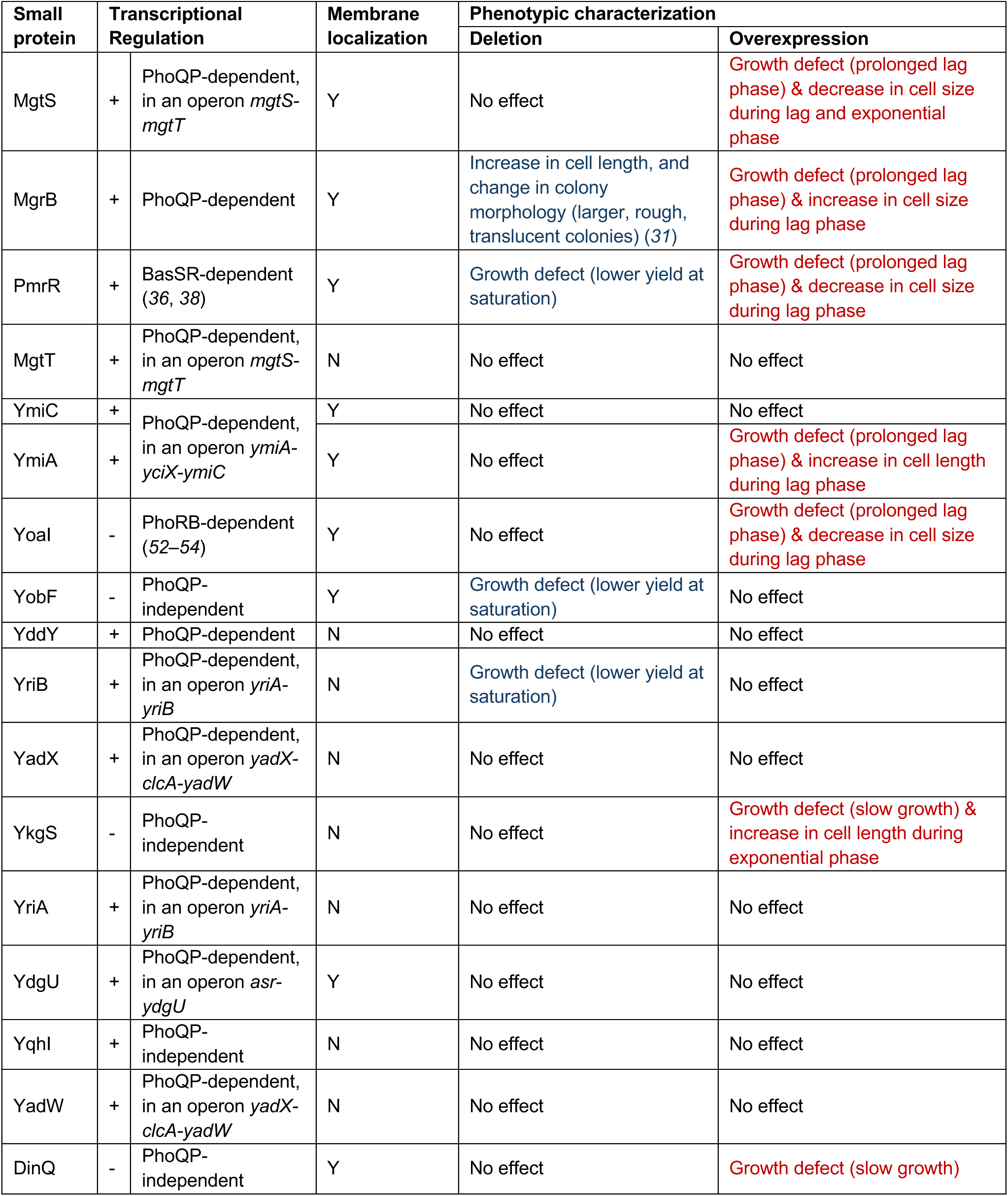
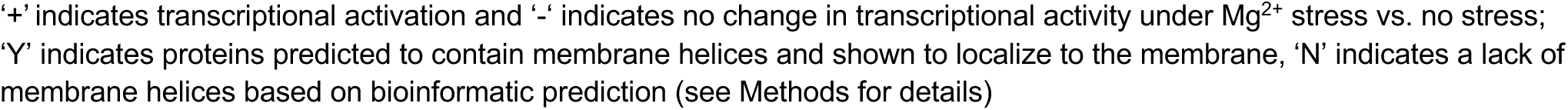
Summary of the 17 small proteins induced under low-magnesium stress in *E. coli*. Small proteins with at least a 3-fold increase in reads mapping to the translation initiation site under low Mg^2+^ relative to no stress are indicated. Results regarding the transcriptional regulation, localization, and characterization of deletion and overexpression phenotypes are summarized for each low-Mg^2+^–induced small protein.

| Small protein | Transcriptional Regulation |  | Membrane localization | Phenotypic characterization |  |
| --- | --- | --- | --- | --- | --- |
|  |  |  |  | Deletion | Overexpression |
| MgtS | + | PhoQP-dependent, in an operon *mgtS-mgtT* | Y | No effect | Growth defect (prolonged lag phase) & decrease in cell size during lag and exponential phase |
| MgrB | + | PhoQP-dependent | Y | Increase in cell length, and change in colony morphology (larger, rough, translucent colonies) (31) | Growth defect (prolonged lag phase) & increase in cell size during lag phase |
| PmrR | + | BasSR-dependent (36, 38) | Y | Growth defect (lower yield at saturation) | Growth defect (prolonged lag phase) & decrease in cell size during lag phase |
| MgtT | + | PhoQP-dependent, in an operon *mgtS-mgtT* | N | No effect | No effect |
| YmiC | + | PhoQP-dependent, in an operon *ymiA-yciX-ymiC* | Y | No effect | No effect |
| YmiA | + |  | Y | No effect | Growth defect (prolonged lag phase) & increase in cell length during lag phase |
| Yoal | - | PhoRB-dependent (52–54) | Y | No effect | Growth defect (prolonged lag phase) & decrease in cell size during lag phase |
| YobF | - | PhoQP-independent | Y | Growth defect (lower yield at saturation) | No effect |
| YddY | + | PhoQP-dependent | N | No effect | No effect |
| YriB | + | PhoQP-dependent, in an operon *yriA-yriB* | N | Growth defect (lower yield at saturation) | No effect |
| YadX | + | PhoQP-dependent, in an operon *yadX-clcA-yadW* | N | No effect | No effect |
| YkgS | - | PhoQP-independent | N | No effect | Growth defect (slow growth) & increase in cell length during exponential phase |
| YriA | + | PhoQP-dependent, in an operon *yriA-yriB* | N | No effect | No effect |
| YdgU | + | PhoQP-dependent, in an operon *asr-ydgU* | Y | No effect | No effect |
| Yqhl | + | PhoQP-independent | N | No effect | No effect |
| YadW | + | PhoQP-dependent, in an operon *yadX-clcA-yadW* | N | No effect | No effect |
| DinQ | - | PhoQP-independent | Y | No effect | Growth defect (slow growth) |

'+' indicates transcriptional activation and '-' indicates no change in transcriptional activity under $Mg^{2+}$ stress vs. no stress; 'Y' indicates proteins predicted to contain membrane helices and shown to localize to the membrane, 'N' indicates a lack of membrane helices based on bioinformatic prediction (see Methods for details)

Ribosome footprints at the start site of an open reading frame are a good proxy for the production of the corresponding protein. However, it is critical to establish whether a given sORF results in a protein product of that specific size. All but one protein encoded by these sORFs, YoaI, were previously validated for expression by genomic epitope-tagging (table S5). In the case of YoaI, the protein could not be detected in rich media or specific growth conditions, including envelope stress, acid stress, or heat shock (*21*, *22*). Given the strong signal for expression of *yoaI* in our Ribo-RET data (∼12-fold increase in RPM for stress vs. no stress) (Fig. 1D), we asked if YoaI is conditionally induced during Mg^2+^ limitation. To examine the production of YoaI, we utilized a strain carrying the chromosomally-encoded fusion protein YoaI-SPA, wherein SPA refers to an 8-kDa sequential peptide affinity tag consisting of 3x FLAG epitope sequence and a calmodulin-binding peptide (*21*, *22*). We grew cells in media containing different concentrations of Mg^2+^, prepared membrane and cytoplasmic fractions, and performed western blot analysis. Consistent with our hypothesis, we detected YoaI-SPA in a Mg^2+^-dependent manner, where the protein abundance was highest at the lowest concentration of Mg^2+^ (Fig. 1, E and F). In addition, this result also confirmed that YoaI was predominantly associated with membranes, consistent with the bioinformatic prediction that it is a transmembrane protein (table S7).

Among the 17 shortlisted small proteins, 14 were not previously associated with induction under low-Mg^2+^ stress. Small proteins YobF (*22*, *49*) and DinQ (*50*) have been linked to other stress responses in *E. coli*, and here we found them to be induced during low-Mg^2+^ stress as well. Many of these small proteins have not been characterized biochemically and have no documented function. In the following sections, we describe our systematic analysis of these low-Mg^2+^–induced small proteins by investigating their transcriptional regulation and cellular localization and characterizing their overexpression and deletion phenotypes.

### Transcriptional regulation of low-Mg^2+^ stress–induced small proteins

To determine changes in the amounts of transcripts encoding stress-induced small proteins, we performed RNA-Seq using cells grown to early exponential phase in high-or low-Mg^2+^ conditions. In cells depleted for Mg^2+^, the PhoQP signaling system is stimulated, leading to PhoP-mediated transcriptional activation of hundreds of genes necessary for the stress response (*24*). As expected, we saw an increase in the mRNA abundances for *phoQ* and *phoP* (fig. S3). Differential expression analysis of our RNA-seq data revealed significant transcriptional activation (fold change threshold of >2 and a p-value threshold of <0.05) under low-Mg^2+^ stress for at least 10 of the 17 small proteins shortlisted, including the *mgtS*, *mgtT*, and *mgrB* transcripts (Fig. 2A). It is worth noting that *yadX* mRNA also showed a small increase (1.7-fold), which was below the >2-fold threshold we applied in our data analysis. However, *yriA*, whose open reading frame overlaps with *yriB* by ∼60%, did not show an increase in the transcript abundance. In addition, we did not see a significant change in the amounts of *yoaI*, *yobF*, *ykgS*, or *dinQ* mRNAs. In these cases, it is possible that the change in transcript abundances was generally low and therefore fell below the significance threshold.

**Fig. 2.**
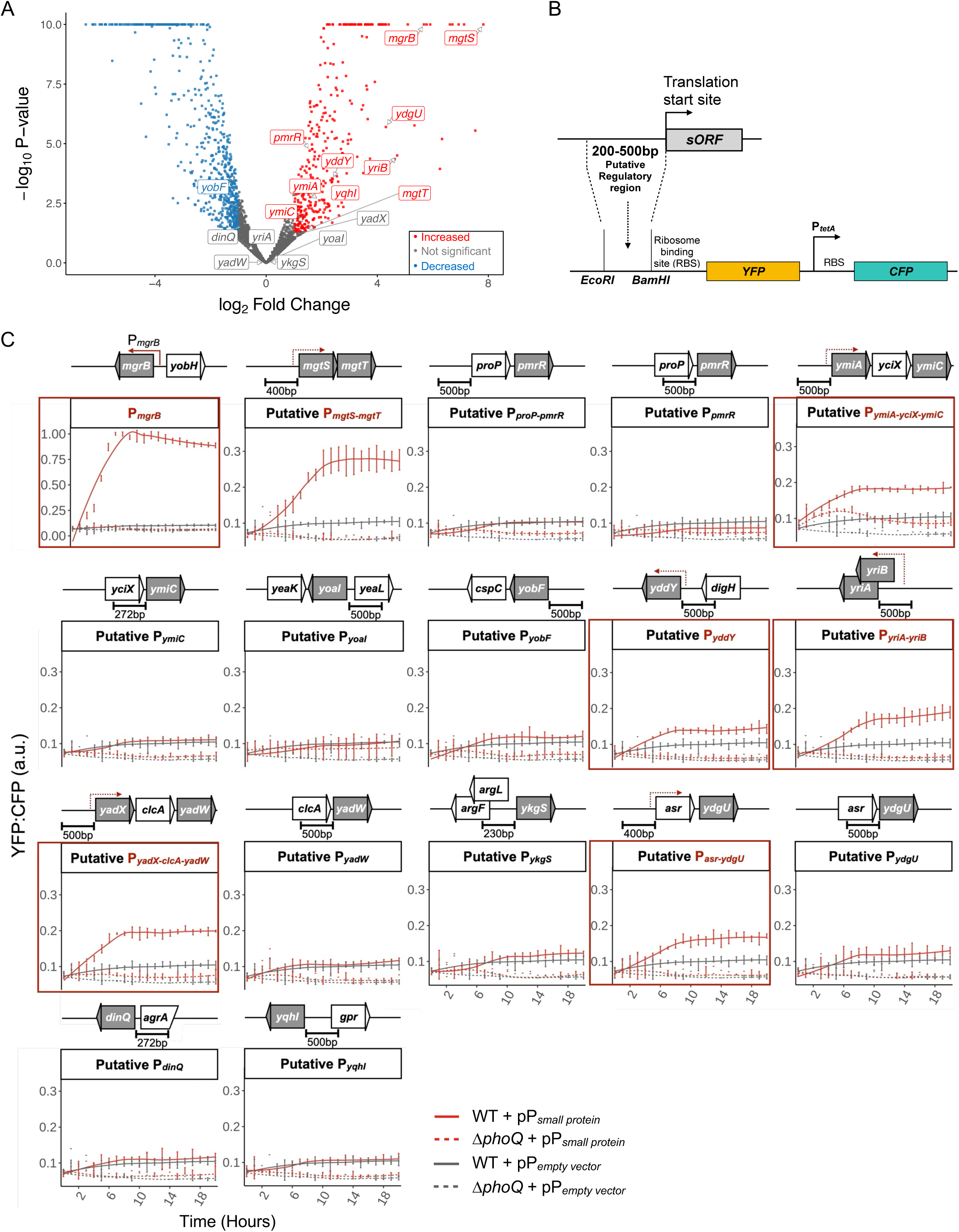
Transcriptional regulation of small proteins induced under low-Mg^2+^ stress. (**A**) Volcano plot illustrating differential expression of transcripts representing small proteins induced under low-Mg^2+^ stress from RNA-Seq data. The -log_10_ P-value of *mgrB* and *mgtS* were 75 and 140, respectively. To prevent distortion from their high values, the transformed p-values were capped at 10 to enhance visualization. Data represent differential expression analysis from n = 2 biological replicates. (**B**) Schematic representation of the transcriptional reporter fusion construct carrying putative regulatory regions (200-500 bp) upstream of the sORFs of interest cloned into a YFP reporter plasmid (pMR120), resulting in plasmids pPJ14, pPJ16-pPJ20, pPJ23, pSV16-pSV21, pSV24, pSV28-pSV29. The reporter plasmid also includes CFP under the control of a constitutively active promoter. Genomic coordinates of the regulatory regions (table S6) and a list of the plasmids (table S2) are provided in the Supplementary Materials. (**C**) Measurements of YFP transcriptional reporter activity relative to CFP (YFP:CFP) for the indicated small proteins during low-Mg^2+^ stress. The solid and dashed lines represent WT *E. coli* MG1655 and *ΔphoQ* cells, respectively, harboring plasmids with either transcriptional fusion for a putative regulatory region of a small protein with *yfp* or *yfp* only. The schematic above each plot depicts the arrangement of the small protein–encoding genes (shown in gray), including the putative regulatory regions and operons where applicable. Transcriptional reporters showing increased activity compared to the empty vector control (pMR120) are outlined in red. The data represent averages and standard errors of the means for n = 4 biological replicates per strain.

Alternatively, the abundances of these transcripts or the proteins they encode might be activated at a post-transcriptional or translational level under low-Mg^2+^ stress. Indeed, *yobF* expression is induced post-transcriptionally during the heat shock response (*22*). The expression of *yobF* under Mg^2+^ limitation may be controlled through a similar post-transcriptional regulatory mechanism.

To delve deeper into the transcriptional control mechanism of the small proteins we sought to identify the regulatory regions upstream of each sORF and determine if the expression was PhoQ-dependent. To identify the regions containing the putative promoters of low-Mg^2+^–induced small proteins, we focused on ∼200 to 500 base pairs upstream of each sORF (table S6). In general, we did not include any annotated full-length ORFs or noncoding RNAs occurring within the regulatory region being tested to avoid interference from the ectopic expression of the corresponding product. In cases where a sORF appeared to be in the middle of an operon, we selected two regions to test: one upstream of the entire operon and another immediately upstream of the sORF of interest. We designed transcriptional reporter constructs to measure the activity of each putative regulatory region corresponding to the following genes or operons as a driver of yellow fluorescent protein (YFP) expression: *mgtST*, *proP-pmrR*, *pmrR*, *ymiA-yclX-ymiC*, *ymiC*, *yoaI*, *yobF*, *yddY*, *yriAB*, *yadX-clcA-yadW*, *ykgS*, *asr-ydgU*, *ydgU, yqhI*, and *dinQ* (Fig. 2, B and C, table S6). Briefly, the putative regulatory region corresponding to each sORF was cloned upstream of a YFP reporter in a single-copy vector (pMR120) that also carries a constitutively expressed cyan fluorescent protein (CFP) reporter under control of the *tetA* promoter as an internal control (Fig. 2B). An *mgrB* transcriptional reporter strain (TIM92) was included as positive control for a gene activated by PhoQ-PhoP (*30*). To analyze which of these sORFs were positively regulated by the PhoQP two-component system, we measured the transcriptional reporter activities in a *phoQ* deletion strain and compared them to that of WT (Fig. 2C).

Consistent with our RNA-seq data, we saw an increase in the PhoQ-dependent transcriptional activity for regulatory regions corresponding to 11 sORFs (*mgrB*, *mgtS*, *mgtT*, *ymiA*, *ymiC*, *yddY*, *yriA*, *yriB, yadX*, *yadW*, and *ydgU*) under low Mg²⁺ (Fig. 2C). Two sets of overlapping (out-of-frame) sORFs including *mgtS-mgtT* and *yriA-yriB*, strongly suggest their association within the same operon. Three other genes, *yadW, ymiC,* and *ydgU* are likely coregulated as part of the operons *yadX-clcA-yadW*, *ymiA-yciX-ymiC*, and *asr-ydgU*, respectively. For all the sORFs occurring within an established or putative operon, the region upstream of the operon showed a higher transcriptional reporter activity in response to stress, but a second region immediately upstream of the given sORF had negligible activity (table S6, Fig. 2C). Whereas the RNA-Seq data represent transcriptomic abundances at a given time point of growth (early exponential phase), the transcriptional reporter assay captures temporal changes in reporter activity and serves as a complementary approach to RNA-Seq, where changes in low-abundance transcripts would be missed. For instance, we did not observe a significant activation of *yadX* and *yadW* in the RNA-Seq data, but we saw a PhoQ-dependent increase in the reporter activity for the putative regulatory region P*_yadX-clcA-yadW_* under low-Mg^2+^ stress. Conversely, for *pmrR* and *yqhI*, although the RNA-Seq data suggested an increase in transcription, we did not detect a significant change in reporter activity under stress, suggesting that there may be distal or additional regulatory factors, outside of the regions included in our reporter constructs, that are involved in controlling their transcription. Indeed, during Mg^2+^ limitation in *Salmonella enterica*, a PhoQP-stimulated small protein PmrD activates PmrB sensor kinase of the PmrA-PmrB signaling system, which positively regulates *pmrR* transcription (*36*). This PmrD-mediated cross-talk between the PhoQ-PhoP and PmrA-PmrB signaling pathways is also active in *E. coli* under specific conditions, including high stimulation of the PhoQP system (*38*, *51*). We did not observe reporter activity for *dinQ, ykgS, yobF*, or *yoaI* in either WT or *ΔphoQ* cells, consistent with the results from the RNA-Seq. In summary, our RNA-Seq and transcriptional reporter assays reveal transcriptional activation of 13 of the 17 small proteins, of which PhoQ promotes at least 11 under Mg^2+^ starvation.

### Mg*^2+^* stress induces *yoaI* expression independently of PhoQ

The expression of *yoaI* is positively regulated by the PhoRB two-component system (*52–54*), which responds to phosphate starvation (*42*). PhoRB signaling is also activated under low-Mg^2+^ stress due to the slowdown of protein synthesis and reduction of cytoplasmic phosphate concentration (*55*).

Activation of the PhoRB signaling includes the positive autoregulation of the *phoRB* genes themselves, increasing their expression as well as the other PhoB-dependent genes (*56*). Consistently, we noticed a 10-fold increase in reads mapping to the PhoB translation start site under Mg^2+^ stress (fig. S2D). We then tested our transcriptional reporter for *yoaI* (P*_yoaI_*-YFP) in a low-phosphate medium and saw increased transcriptional activity for P*_yoaI_* when compared to no-stress condition, in a PhoB-dependent manner (Fig. 3A). This transcriptional activation was PhoQ-independent, because we saw an increase in the P*_yoaI_* reporter in *ΔphoQ* cells as well (Fig. 3A).

**Fig. 3.**
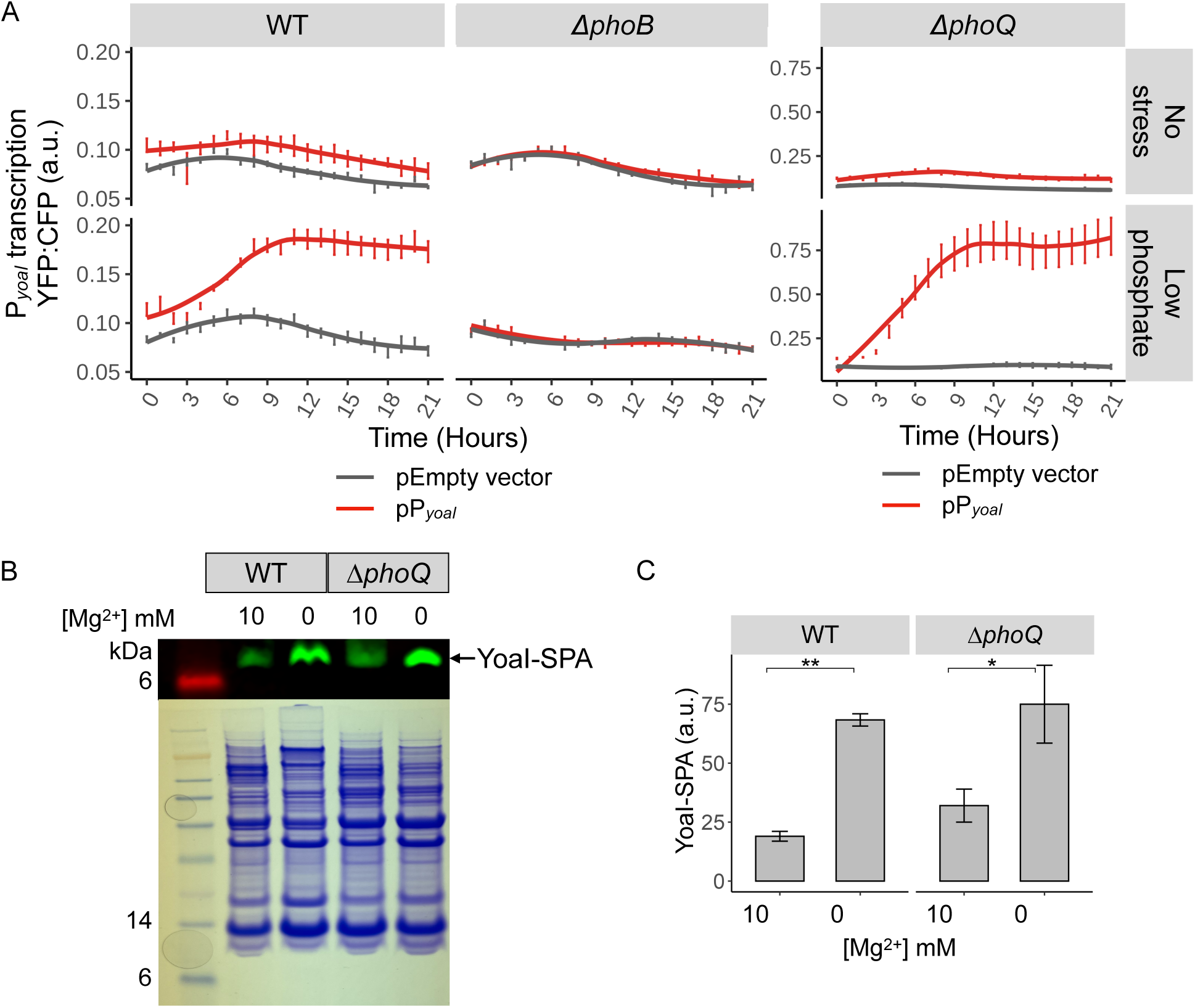
Regulation of YoaI during magnesium limitation. (**A**) Transcriptional activity of *yoaI* was measured as a function of YFP reporter fluorescence relative to CFP in *E. coli* WT (MG1655), *ΔphoB* (SV48), and *ΔphoQ* (TIM100) strains under low-phosphate stress as indicated. The red and gray lines represent cells carrying plasmid-encoded P*_yoaI_-yfp* transcriptional reporter (pPJ18) and *yfp*-only control vector (pMR120), respectively. The data show average YFP:CFP fluorescence with standard error of the mean, derived from n = 4 biological replicates per strain. (**B**) Detection of YoaI protein in *E. coli* WT (GSO317) and *ΔphoQ* (SV60) strains containing the *yoaI-SPA* genomic translational fusion in media containing MgSO_4_ at the specified concentrations. Membrane fractions were analyzed by western blotting using an antibody specific for FLAG and by Coomassie Brilliant Blue staining. The data represent results from n = 3 biological replicates per group. (**C**) Quantification of YoaI-SPA protein amounts from the western blot in (B). Bars represent the mean ± standard error of the mean from n = 3 biological replicates per group. P-values were calculated using a t-test as indicated; *P < 0.05, **P ≤ 0.01.

Together, these results show that *yoaI* transcription is induced by PhoB, and there is no significant effect of low-Mg^2+^ stress on *yoaI* transcription based on RNA-Seq and transcriptional reporter data (Fig. 2, A and C). However, YoaI protein abundance increased under low-Mg^2+^ conditions, and we wondered if PhoQ enhances Yoal abundance. To test this possibility, we examined YoaI protein in WT and *ΔphoQ* cells in high- and low-Mg^2+^ conditions by western blotting. Our results demonstrate that YoaI abundance remained Mg^2+^-dependent in a *ΔphoQ* strain, mirroring the pattern seen in WT (Fig. 3, B and C), suggesting that PhoQ does not affect YoaI abundance. Therefore, YoaI expression is induced in a PhoQ-independent manner under Mg^2+^ limitation, potentially mediated by a post-transcriptional regulatory mechanism that enhances translation initiation.

### About half of the low-Mg^2+^ stress–induced small proteins localize to the membrane

As a first step to classifying the 17 stress-induced small proteins, we checked if a given small protein is likely associated with the membrane or cytoplasm. Most of these proteins have limited or no documented information, so their cellular localization can provide insights into their potential mechanisms of action and interactions within the cell. Using bioinformatic tools TMHMM (*57*), TMPred (*58*), and Phobius (*59*) to predict membrane helices, we found that at least 9 of our candidates are putative transmembrane proteins. Three of them, MgrB, MgtS, and DinQ, localize to the membrane in *E. coli* (*21*, *50*), and PmrR from *S. enterica* is membrane-associated (*36*). Our bioinformatic analyses predicted that small proteins YdgU, YmiA, YmiC, YoaI, and YobF also localize to the membrane (table S7).

To confirm the predicted membrane association and visualize the small protein localization, we used epitope tagging with green fluorescent protein (GFP). Taking into account the predicted orientation of the putative transmembrane protein, we created either an N- or C-terminal fusion of GFP by tagging the end that was expected to face the cytoplasm (Fig. 4A, table S7) to ensure proper folding of GFP (*60*). Accordingly, YdgU, YmiA, and YmiC were tagged at the N-terminus, and YoaI was tagged at the C-terminus. Upon observation by fluorescence microscopy, cells expressing GFP-tagged YdgU, YmiA, YmiC, and YoaI revealed bright fluorescence at the cell periphery, indicative of membrane localization, similar to the control small transmembrane protein, MgrB (Fig. 4B). Cells expressing GFP alone showed uniform fluorescence indicative of cytoplasmic localization.

**Fig. 4.**
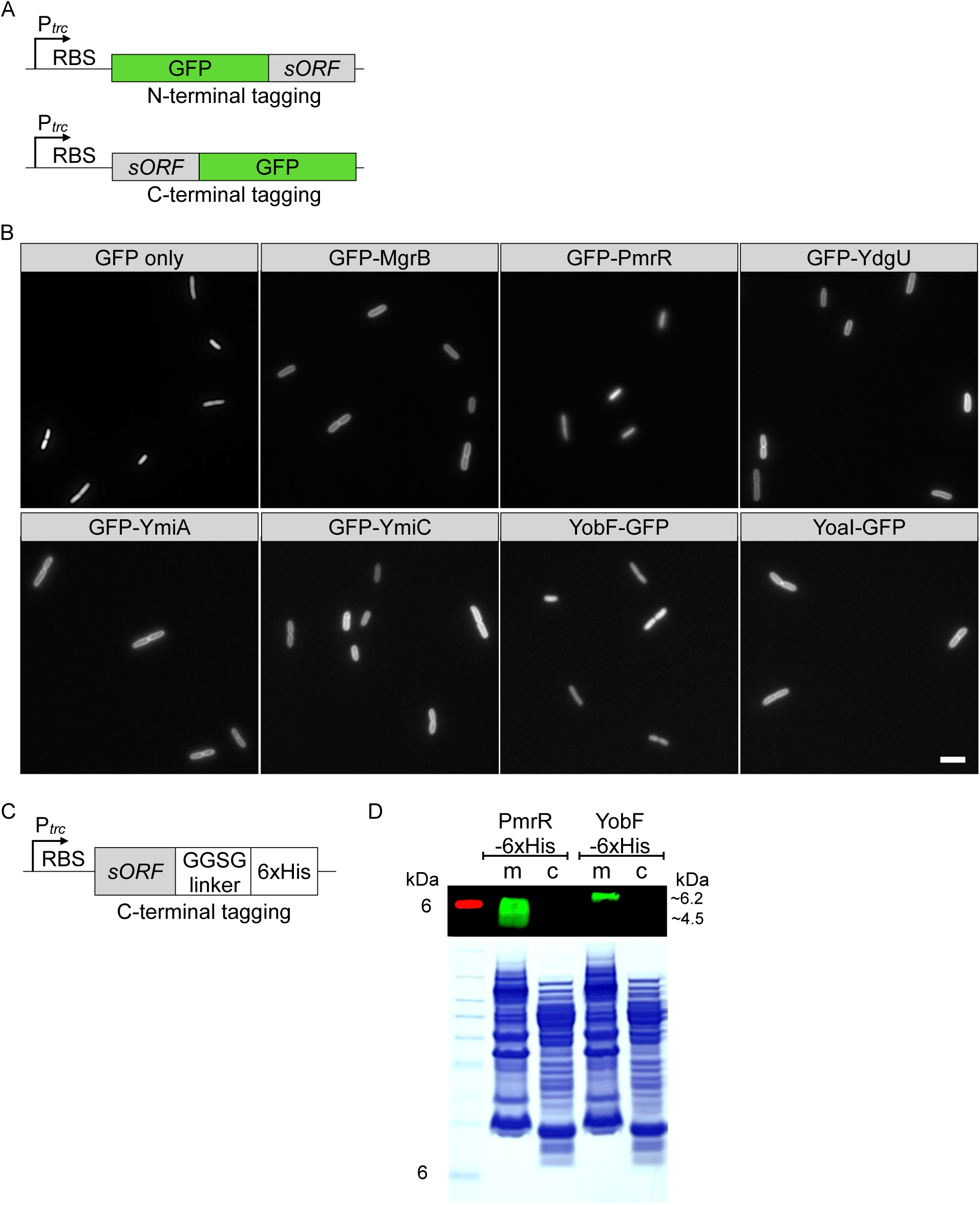
Subcellular localization of epitope-tagged small proteins. (**A**) Schematic representation of GFP-tagged small protein constructs. GFP was inserted N- or C-terminal to the sORF downstream of a ribosome-binding site (RBS), and expression was driven by an inducible *trc* promoter (P*_trc_*). (**B**) Fluorescence micrographs of *E. coli* K-12 MG1655 cells expressing GFP-MgrB (pAL38), GFP-YdgU (pPJ3), GFP-YmiA (pPJ12), GFP-YmiC (pPJ13), YoaI-GFP (pPJ4), GFP-PmrR (pPJ2), YobF-GFP (pPJ6), or GFP only (pAL39). n = 2 biological replicates per strain. Scale bar, 5 µm. (**C**) Schematic representation of 6xHis-tagged small protein constructs. His_6_ was inserted N- or C-terminal to the sORF downstream of an RBS, and expression was driven by an inducible *trc* promoter (P*_trc_*). (**D**) Membrane (m) and cytoplasmic (c) fractions of *E. coli* K-12 MG1655 cells expressing PmrR-6xHis (pSV14) or YobF-6xHis (pSV26) were analyzed by western blotting using an antibody specific for the His tag and by Coomassie Brilliant Blue staining. The data represent results from n = 2 biological replicates per group.

For PmrR, we constructed an N-terminal fusion with GFP, where the tag was expected to localize to the cytoplasm. However, we did not observe membrane localization (Fig. 4B). Considering the predicted membrane helices, this suggests that the tag interferes with PmrR localization. Alternately, we tagged the C-terminus of *E. coli* PmrR with a hexahistidine (6xHis) tag (Fig. 4C) and performed western blotting using membrane and cytoplasmic fractions. PmrR-6xHis strongly associated with the membrane fraction (Fig. 4D), consistent with the bioinformatic prediction and data from *S. enterica* PmrR (*36*). As in the case of PmrR, YobF-GFP localized to the cytoplasm, which was inconsistent with its predicted membrane localization (table S7). Therefore, we used a 6xHis-tagged version of YobF, which showed a strong association with the membranes (Fig. 4D). Overall, among the 17 small proteins induced under low Mg^2+^, we found 9 of them to be integral membrane proteins.

### Loss of PmrR, YobF, or YriB cause Mg^2+^ stress–specific growth defects

To identify potential defects in growth or cellular morphology caused by the absence of each small protein, we analyzed single gene deletions of *mgrB*, *mgtS*, *mgtT*, *pmrR*, *yadW*, *yadX*, *yddY*, *ykgS*, *ymiC*, *yqhI*, *yobF, dinQ*, *ydgU*, *ymiA*, and *yoaI* and a double deletion removing *yriA* and *yriB* in the *E. coli* genome. *mgtT* lies in an operon with *mgtS* (*41*), with an out-of-frame overlap between the stop codon of MgtS and the start codon of MgtT. Therefore, we carefully generated an *mgtS* deletion without affecting the start codon of MgtT, and similarly, an *mgtT* deletion while keeping the stop codon for MgtS intact (for details see Materials and Methods and table S1). Because *yriA* and *yriB* have overlapping open reading frames in which ∼60% of *yriA*’s ORF overlaps with *yriB* and ∼80% of the *yriB* gene overlaps with that of *yriA* (*41*), we first deleted the combined *yriAB* ORF to see if there is a phenotype. Of the 17 small proteins, 12 displayed no discernible growth defects upon deletion relative to the WT strain when grown under Mg^2+^ limitation. However, four mutants – namely *ΔpmrR, ΔyobF, ΔyqhI,* and *ΔyriAB* – exhibited reduced growth yields when grown over 24 hours in low-Mg^2+^ medium (Fig. 5A). The cells carrying these deletions entered the stationary phase earlier than did WT cells. PmrR and YobF are membrane-associated proteins, whereas YqhI, YriA, and YriB are putative soluble proteins (Fig. 4D, table S7).

**Fig. 5.**
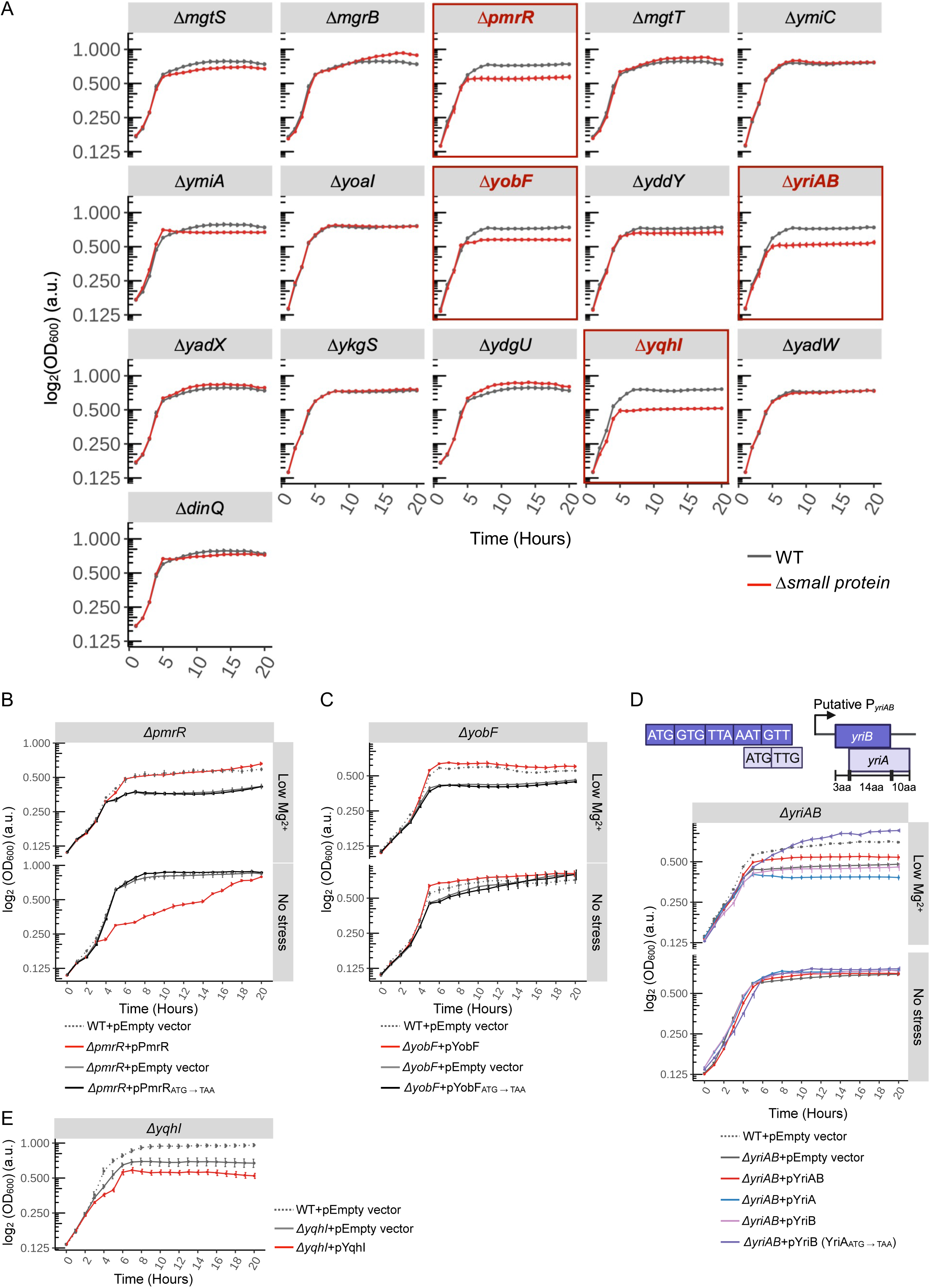
Effect of deleting genes encoding low-Mg^2+^ stress–induced small proteins on bacterial growth. (**A**) Growth curves of WT *E. coli* MG1655 and mutants corresponding to the deletions of small protein–encoding genes in low-Mg^2+^ medium (strains SV64, AML67, SV35, SV31, SV41, SV57, SV59, SV33, SV29, SV25, SV39, SV43, SV58, SV37, SV27, and SV56). Deletion mutants showing reduced growth yield compared to WT cells are outlined in red. (**B and C**) Complementation of growth phenotypes associated with *pmrR* or *yobF* deletion. Growth curves are shown for Δ*pmrR* (SV35) (B), and Δ*yobF* (SV33) (C) deletion strains carrying an empty vector (pEB52) or complemented with plasmids encoding WT small proteins, pPmrR (pSV35) or pYobF (pJS5) or variants in which the start codon (ATG) was mutated to a stop codon (TAA), pPmrR_ATG→TAA_ (pSV38), and pYobF_ATG→TAA_ (pSV45). (**D**) Schematic of the overlapping genes *yriAB* and growth curves of deletion strain *ΔyriAB* (SV25) complemented with pYriAB (pSV58), pYriA (pJS1), pYriB (pSV37), pYriB (YriA_ATG→TAA_) (pSV59), or an empty vector (pEB52). (**E**) Growth curves of deletion strain *ΔyqhI* (SV37) carrying an empty vector (pEB52) or complemented with a plasmid encoding YqhI. For panels (B) to (D), WT cells containing an empty vector (pEB52, dashed gray line) are included as a control. All data represent averages and standard errors of means for n = 4 biological replicates per strain.

To determine whether these growth defects were specifically related to the loss of these small proteins, we complemented the deletion strains with plasmids encoding the corresponding small proteins and induced with IPTG. Successful rescue of growth was observed for *ΔpmrR* and *ΔyobF* (Fig. 5, B and C; top). To confirm that this complementation was specific to the small protein and not due to a cryptic ORF or regulatory element present within the small protein coding region, we prepared plasmids carrying *pmrR* and *yobF* in which their start codons were mutated to stop codons. When tested for complementation in *ΔpmrR* and *ΔyobF*, respectively, the start codon mutants retained the growth defect similar to that observed for the empty vector (Fig. 5, B and C; top), reinforcing that the growth phenotype is specific to the small proteins themselves. In addition, we investigated whether the growth defect observed with *pmrR* and *yobF* deletions was specific to low-Mg^2+^ stress. Our results indicate that deletion of *pmrR* or *yobF* did not cause growth defects in the absence of stress (Fig. 5, B and C; bottom), and the phenotype was specific to Mg^2+^-limited conditions. On the other hand, expression of plasmid-encoded PmrR in the absence of stress resulted in a growth defect (Fig. 5B, bottom), suggesting that increased expression of PmrR under no-stress conditions may be detrimental.

Given the growth phenotype observed for *ΔyriAB*, we performed complementation with either plasmid-encoded YriA, YriB, or both to determine whether the growth defect was due to the loss of one or both proteins. Complementation with either YriA or YriB alone failed to restore growth, but a plasmid expressing the region from *yriB* to *yriA,* mimicking the native genomic arrangement (*yriAB*), partially rescued the growth defect (Fig. 5D). In our plasmid construct encoding YriB and YriAB, there is a possibility of a spurious small protein produced due to the intact, out-of-frame YriA start codon within the *yriB* ORF (Fig. 5D, schematic) that potentially interferes with the complementation. To address this issue, we created a modified *yriB* construct in which the YriA start codon ATG was mutated to TAA, a substitution that also changes codons 4 (Asp to Ile) and 5 (Val to Ile) in YriB. This variant of YriB conferred full complementation (Fig. 5D), indicating that loss of YriB contributes to the growth phenotype, at least in part. In addition, we observed that the growth defect associated with *yriAB* deletion was specific to the low-Mg^2+^ stress condition.

Finally, the growth defect observed in *ΔyqhI* was not complemented with a plasmid encoding YqhI (Fig. 5E), which suggests that this phenotype is not directly related to the YqhI protein itself. We did not see a growth defect in the *ΔmgtS* strain when grown in low-Mg^2+^ medium (Fig. 5A), contrasting with the previous report (*34*). This discrepancy might be due to differences in strain construction and/or growth media. Together, our results underscore the complexities in studying overlapping small ORFs and their phenotypes.

### Seven small proteins affect growth and cell morphology upon overexpression

As a parallel approach to analyzing loss-of-function phenotypes, we explored the phenotypes associated with the overexpression of small proteins. We cloned the genes corresponding to 16 of the 17 untagged small proteins into an IPTG-inducible vector. Because we were unable to generate a DinQ clone in the IPTG-inducible pEB52 background, we cloned DinQ into an arabinose-inducible (DinQ) vector. Upon induction of the transgenes in low-Mg^2+^ medium, slow growth or a prolonged lag phase was observed for 7 out of the 17 small proteins tested: MgtS, MgrB, PmrR, YmiA, YoaI, YkgS, and DinQ (Fig. 6), of which six proteins are membrane-associated (MgtS, MgrB, PmrR, YmiA, YoaI, and DinQ). In the case of MgtS, MgrB, PmrR, YmiA, and YoaI, induction of expression with IPTG led to an extended lag phase lasting for ∼8-10 hours, after which the cells grew exponentially at growth rates similar to that of control cells carrying an empty vector, eventually reaching saturation to a similar extent. In contrast, cells expressing YkgS or DinQ did not display a long lag phase but grew at a slower rate, gradually catching up to the same growth yield as the control cells. The growth defect due to DinQ overexpression aligns with previous studies indicating that its overexpression disrupts membrane potential and depletes intracellular ATP (*50*). Overexpression of the other 10 small proteins, including 3 membrane proteins YdgU, YmiC, and YobF, had a negligible impact on cell growth under the same low-Mg^2+^ condition, suggesting that cells may tolerate high amounts of these proteins (Fig. 6).

**Fig. 6.**
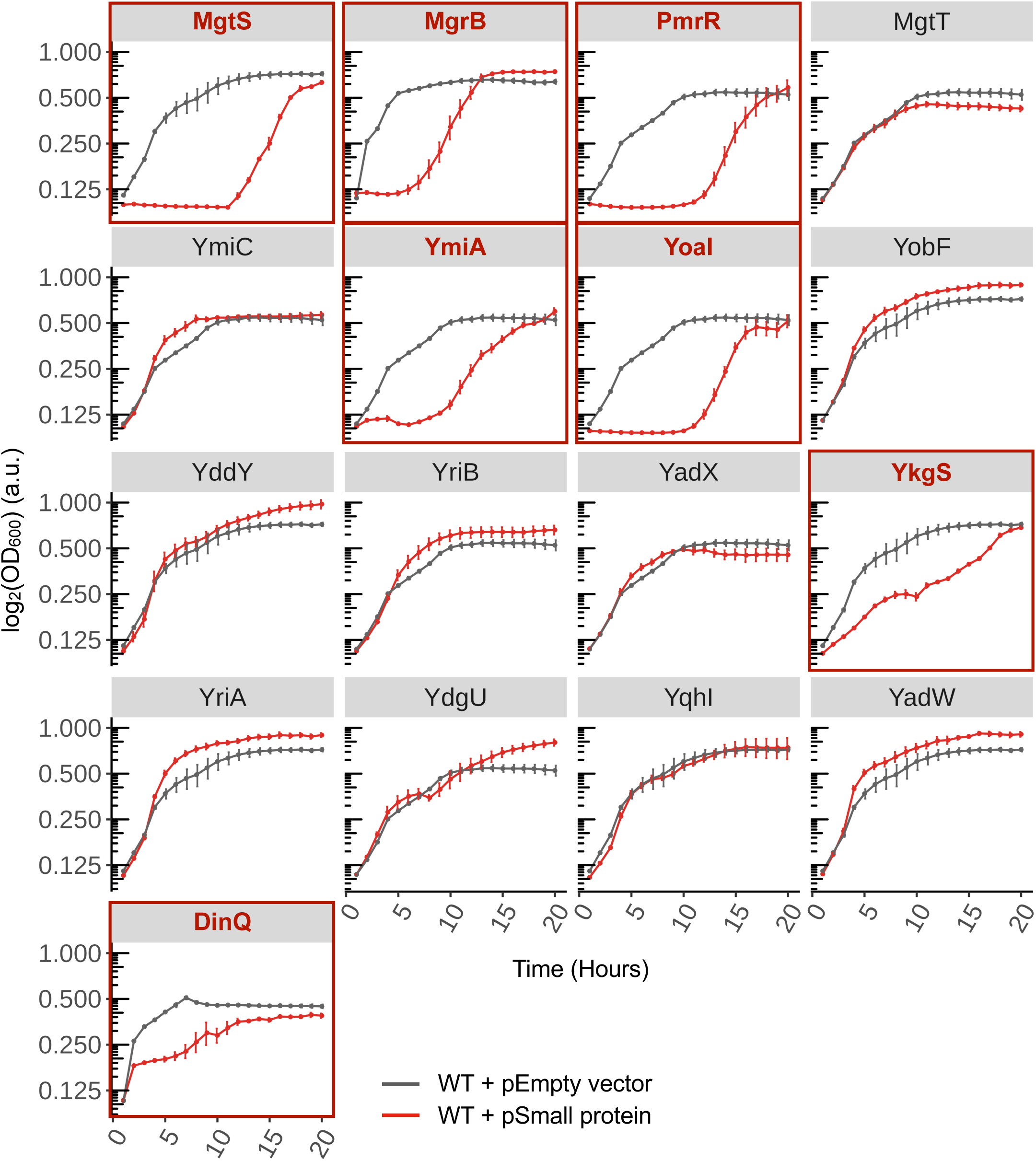
Effect of overexpression of low-Mg^2+^ stress–induced small proteins on growth. Growth curves of WT *E. coli* MG1655 cells grown under low-Mg^2+^ stress carrying an empty vector or overexpressing each of the 17 stress-induced small proteins (encoded by plasmids pSV30–37, pSV54, pSV60, or pJS1–7). Small proteins that caused a growth defect or delay when overexpressed are outlined in red. All data represent averages and standard error of means for n = 4 biological replicates per strain.

To be cautious in interpreting the overexpression data, we considered the possibility that the observed growth defects might be attributed to a cryptic small open reading frame or regulatory element within the small ORF being overexpressed. To check if the growth defects were indeed small protein–dependent, we prepared plasmid constructs containing variants of the small protein in which the start codon (ATG) was substituted with a stop codon (TAA). For all 7 small proteins displaying overexpression phenotypes, we repeated the growth assay with the start codon mutants of each small protein. Expression of these mutant small proteins did not cause a growth defect, and the cells exhibited similar growth patterns to those carrying the empty vector (Fig. 7A), reinforcing that the observed growth defects were specific to the overexpression of these small proteins.

**Fig. 7.**
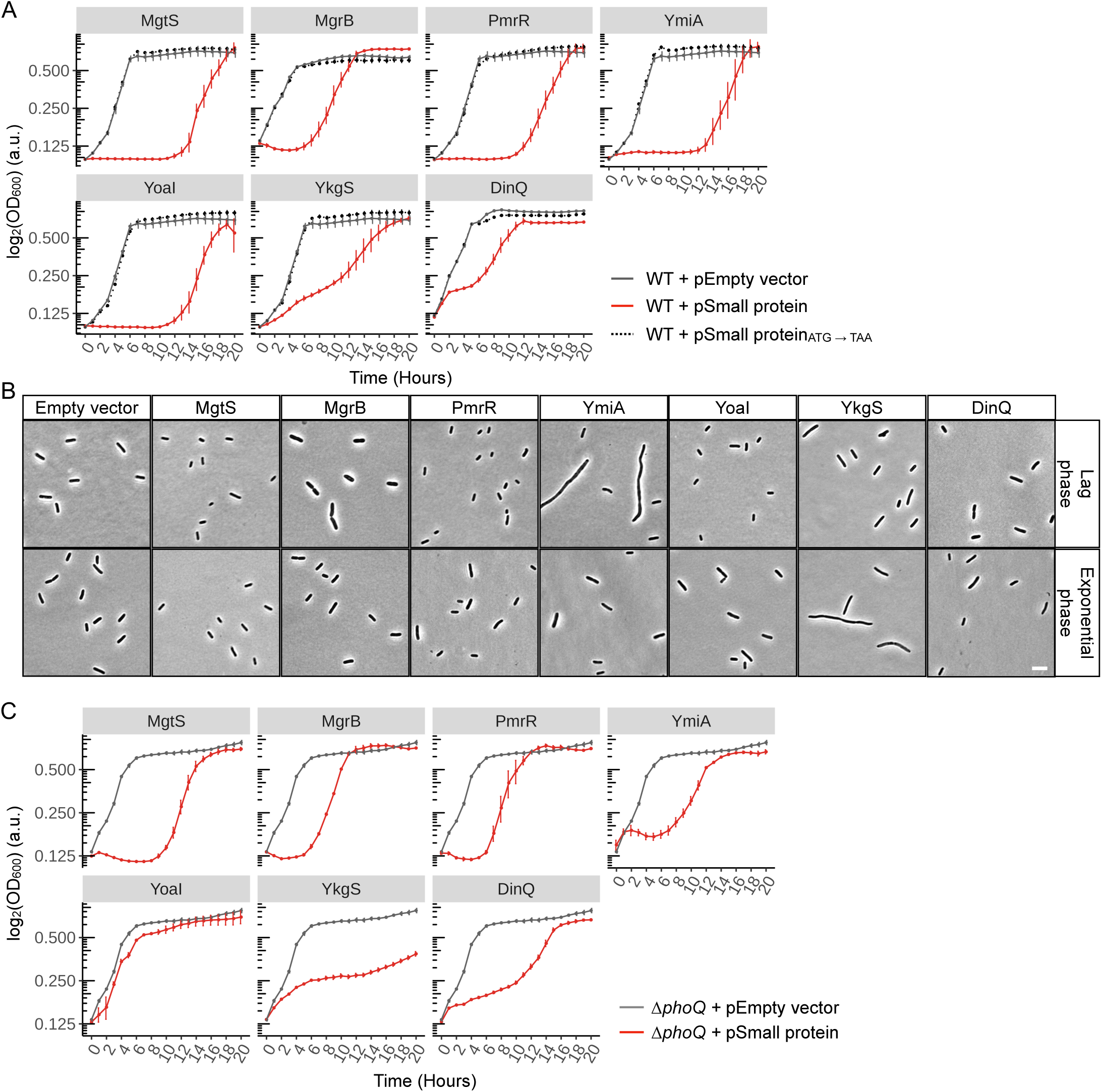
Analysis of growth defect and cell size changes associated with overexpression of MgtS, MgrB, PmrR, YkgS, YoaI, YmiA, or DinQ. (**A**) Growth curves of WT *E. coli* MG1655 cells expressing an empty vector, the indicated small proteins, or variants of the small protein in which the start codon (ATG) of each small protein was mutated to a stop codon (TAA). Plasmids pSV34-36, pSV38-42, pSV54-55, pSV60-61, or pJS4, pJS7 (table S2) were used for these experiments. Data represent averages and standard error of means for n = 4 biological replicates per strain. (**B**) Representative phase contrast micrographs of cells during the lag and exponential phases of growth after the nduction of plasmids encoding the indicated small proteins. Scale bar, 5 µm. Data are representative of n = 2 biological replicates per strain. (**C**) Growth curves of *ΔphoQ* (TIM202) cells expressing the indicated small proteins or empty vector (pEB52 or pBAD24). Plasmids pSV34-36, pSV54, pSV60, pJS4, and pJS7 (table S2) were used for these experiments. Data represent averages and standard error of means for n = 4 biological replicates per strain.

In addition to the impact on growth, we explored whether overexpression of the 7 small proteins that slowed cell growth (MgtS, MgrB, PmrR, YmiA, YoaI, YkgS, and DinQ) also influenced cell size or morphology (Fig. 6, 7A). We hypothesized that cells overexpressing these small proteins may undergo morphological changes as part of adaptation during the extended lag and log phases of growth. Using phase contrast microscopy, we imaged cells overexpressing the small proteins during the lag and exponential growth phases in low-Mg^2+^ medium. As controls, we used WT cells carrying either an empty vector or a plasmid containing a variant of the small ORF in which its start codon (ATG) was mutated to a stop (TAA). The control cells exhibited a roughly uniform cell length with volumes around 1.1-1.3 fl and 1.2-1.4 fl in the lag and log phases, respectively. The overexpression of MgtS, MgrB, PmrR, YkgS, YoaI, or YmiA caused distinct changes in cell sizes and/or lengths (Fig. 7B, fig. S4). Specifically, overexpression of MgtS resulted in smaller cell size (Fig. 7B), with a reduction in cellular volume to ∼30-36% of the control cells carrying either empty vector or start codon mutants in both lag and log phases (fig. S4). Similarly, overexpression of PmrR or YoaI led to a ∼29% decrease in cell size during the lag phase, however, the cell size reverted to that of control cells during the exponential phase (Fig. 7B, fig. S4). In contrast, overexpression of MgrB led to an increase in cell size, with an average volume of 2.2 fl, ∼1.6-fold larger than the controls in the lag phase (Fig. 7B, fig. S4), akin to *ΔphoQ* cells grown in low-Mg^2+^ conditions (fig. S5, A and B). That the overexpression of MgrB caused a phenotype similar to the loss of PhoQ is consistent with MgrB being a known inhibitor of PhoQ (*29*, *30*). These cells eventually returned to normal size (cell volume of 1.2 fl) in the exponential phase. Overexpression of YkgS or YmiA caused varying extents of cell elongation and increased cell volume (Fig. 7B, fig. S4). Cells overexpressing DinQ did not show a significant change in cell size in the exponential phase (Fig. 7B, fig. S4). Overall, the slow or delayed growth observed here, along with the diverse morphological phenotypes at distinct growth stages, reflects how cells adapt in response to the overexpression of small proteins mentioned above.

In *Salmonella enterica,* PmrR inhibits the inner membrane protein LpxT, which plays a role in lipopolysaccharide modification (*36*, *38*, *39*). We explored whether the growth defects associated with PmrR deletion and overexpression in *E. coli* were mediated through its interaction with LpxT. Expression of a plasmid encoding PmrR in a *ΔlpxT* background did not alleviate the growth defects regardless of stress (fig. S6A). It is possible that higher LpxT amounts, resulting from the absence of PmrR, could lead to growth defects. However, both single deletions (*ΔlpxT* and *ΔpmrR*) and the double deletion (*ΔpmrR ΔlpxT*) exhibited growth defects under Mg^2+^ stress (fig. S6B). This result suggests that the growth defect upon *pmrR* deletion is independent the interaction of PmrR with LpxT.

### YoaI mediates cross-talk between the PhoR-PhoB and EnvZ-OmpR signaling systems

We hypothesized that the growth delay observed upon overexpression of the 7 small proteins described above may be PhoQ-dependent, given the pivotal role of PhoQ in low-Mg^2+^ stress. To explore this, we evaluated the effects of overexpressing these proteins in a *ΔphoQ* strain under Mg^2+^ limitation. The growth phenotype associated with small protein overexpression was retained in the *ΔphoQ* cells, similar to those observed in the WT strain for 6 out of 7 candidates: MgrB, MgtS, PmrR, YmiA, YkgS, and DinQ (Fig. 7C), indicating that these effects are PhoQ-independent. However, in the case of YoaI overexpression, the extended lag phase observed in WT cells was completely abolished in the *ΔphoQ* cells, suggesting a direct or indirect link between PhoQ and YoaI, with PhoQ functioning upstream of YoaI in this regulatory pathway.

To test if YoaI and PhoQ physically interacted with each other, we employed a bacterial two-hybrid (BACTH) assay based on reconstituting split adenylyl cyclase (CyaA) (*61*, *62*). In this system, CyaA activity is restored when the two fragments T18 and T25 come close to one another, leading to an increase in the amount of cyclic AMP (cAMP) and subsequent expression of β-galactosidase (β-gal) from the *lac* promoter. We fused the T18 fragment to YoaI (T18-YoaI) and tested it against T25 fused to PhoQ (T25-PhoQ). We included the T18-MgrB and T25-PhoQ pair as a positive control and two other histidine kinases, PhoR and EnvZ, each fused to the T25 fragment as additional controls. In this experiment, YoaI did not show an interaction with either PhoQ or PhoR, however, there was a strong signal between YoaI and EnvZ that was comparable to the positive control (Fig. 8A). EnvZ is an osmosensor, integral to the EnvZ-OmpR two-component system, which regulates porins in response to changes in osmolarity, pH, temperature, and growth phase (*43*). Several small membrane proteins modulate the activities of sensor kinases (*14*). MzrA, a 127-amino acid protein, stimulates EnvZ, resulting in higher amounts of phosphorylated OmpR (*63*). Based on the BACTH interaction between YoaI and EnvZ, we suspected YoaI might influence the activity of the EnvZ-OmpR signaling system. To address this possibility, we utilized an EnvZ-OmpR–activated transcriptional reporter strain with a chromosomal fusion of mScarlet to the promoter of *omrB* (P*_omrB_*) (*64*). We measured *omrB* promoter activity in cells grown in low-Mg^2+^ vs. no-stress media and observed a 2.5-fold increase in EnvZ-activated transcription under stress in WT cells (Fig. 8B). The significantly high activity of the *omrB* reporter in low-Mg^2+^ conditions suggests that increased abundance of YoaI protein may mediate this effect (Fig. 1, D to F, Fig. 3, B and C). Upon deletion of *yoaI*, expression of the *omrB* promoter reporter was reduced by ∼1.4-1.5-fold in both low- and high-Mg^2+^ media, suggesting that YoaI affects EnvZ-OmpR activity regardless of stress. Together with our observation that YoaI-SPA abundance increased under low-Mg^2+^ stress (Fig. 3, B and C, Fig. 1, E and F), the higher activity of the *omrB* promoter could be attributed to the positive interaction of YoaI with EnvZ. Because *yoaI* is transcriptionally activated by the phosphate-responsive PhoB-PhoR system, we asked whether phosphate-limiting growth conditions had a greater effect on EnvZ-activated *omrB* expression than did Mg^2+^ limitation. In cells grown in phosphate-excess and -limiting conditions, there was a ∼1.8-2-fold decrease in *omrB* reporter activity in *ΔyoaI* cells relative to WT cells, irrespective of the stress (Fig. 8B). Overall, the observed reduction in *omrB* transcription in *ΔyoaI* cells relative to WT was similar in both Mg^2+^- and phosphate-limiting conditions, suggesting that YoaI activates EnvZ to comparable extents under the two stress conditions. Consistent with the idea that YoaI affects *omrB* transcription through EnvZ, an *ΔenvZ ΔyoaI* double deletion strain exhibited negligible *omrB* reporter activity across all conditions tested (Fig. 8B).

**Fig. 8.**
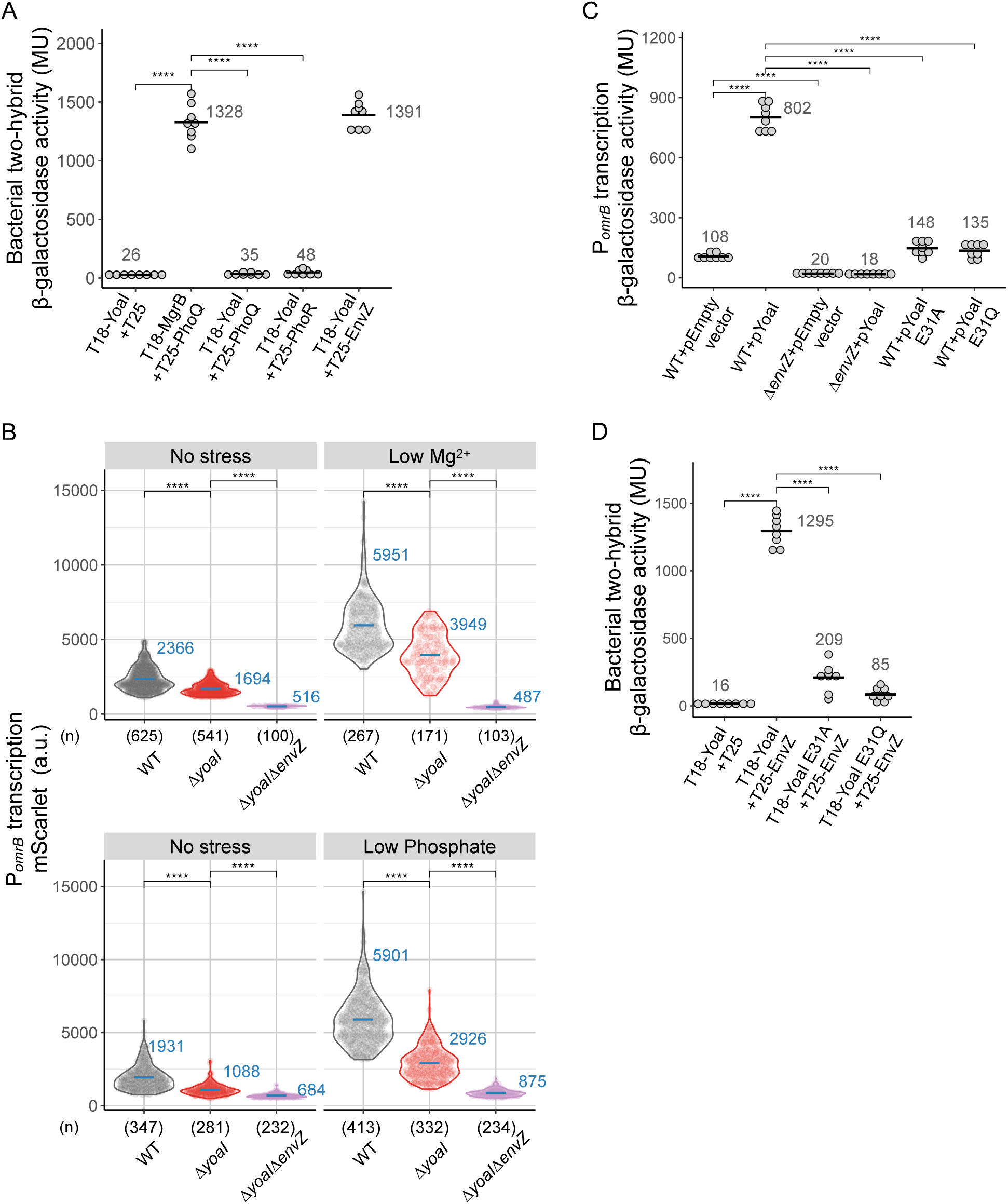
Effect of YoaI expression on the EnvZ-OmpR system. (**A**) BACTH assay reporting β-gal activity to measure interactions between T18-YoaI (pSV47) and T25 fused to PhoQ (pAL27), PhoR (pSY68), or EnvZ (pKK14). T25 alone was used as a negative control, and T18-MgrB (pAL33) combined with T25-PhoQ (pAL27) was used as a positive control. Mean values are indicated with black bars and numbers beside each set of data points. n = 8 biological replicates per strain. (**B**) Transcriptional activity of *omrB* was measured as a function of mScarlet reporter fluorescence in *E. coli* MG1655 (SV99), *ΔyoaI* (SV95), and *ΔyoaIΔenvZ* (SV101) strains at the stationary phase. Each circle corresponds to a single cell. Cells were grown in either minimal A or MOPS minimal medium for Mg^2+^ or phosphate stress experiments, respectively. n ≥ 100 cells per strain from 3 biological replicates per condition. The numbers below the x-axis indicate the number of cells analyzed per group. Mean fluorescence is shown in blue bars and numbers. (**C**) BACTH assay to assess interactions between T18-YoaI (pSV47) and its derivatives T18-YoaI E31A (pJS15) and T18-YoaI E31Q (pJS16) with T25-EnvZ (pKK14). Mean values are indicated by black bars and numbers. n = 8 biological replicates per group. (**D**) Transcriptional activity of *omrB* was measured as a function of *lacZ*-driven β-galactosidase activity in WT (JM2110) and *ΔenvZ* (SV98) strains upon overexpression of WT YoaI (pSV34), YoaI E31A (pSV70), YoaI E31Q (pJS13), or an empty vector (pEB52). Mean values indicated by black bars and numbers, n = 8 biological replicates per grpup. For all panels, P-values represent the results of a t-test as indicated, ****P ≤ 0.0001.

Furthermore, we examined *omrB* transcription upon plasmid overexpression of YoaI and found a greater than 7-fold increase in the β-galactosidase activity as a function of the *lacZ* reporter driven by the *omrB* promoter (P*_omrB_*-*lacZ*) in WT cells expressing YoaI relative to the control cells carrying an empty vector (Fig. 8C). *ΔenvZ* cells expressing plasmid-encoded YoaI showed minimal reporter activity, indicating that YoaI acts through EnvZ. We also tested two other promoter reporters of EnvZ-OmpR–regulated genes, *ompF* (P*_ompF_*-YFP) and *ompC* (P*_ompC_*-CFP) (*65*), which showed a reciprocal change of ∼2-fold in their transcriptional activities when YoaI was expressed from a plasmid (fig. S7, A and B), consistent with the activation of EnvZ (*43*). In summary, these findings strongly indicate that YoaI interacts with and stimulates EnvZ activity, potentially enhancing OmpR phosphorylation. To test whether the overexpression phenotype associated with YoaI overexpression depended on EnvZ, we monitored the growth of *ΔenvZ* cells expressing YoaI from a plasmid. These cells experienced a long lag phase similar to that observed in WT cells (fig. S7C), suggesting that this growth phenotype was independent of YoaI’s interaction with the EnvZ-OmpR signaling pathway.

### A conserved glutamate in small protein YoaI mediates its interaction with EnvZ sensor kinase

To gain a mechanistic understanding of how the small protein YoaI may interact with and promote activation of its target EnvZ, we first looked for conserved amino acid residues within YoaI. We generated a multiple sequence alignment of YoaI orthologs from 57 representative Gammaproteobacteria (fig. S8A) and identified several conserved residues. Amino acids at three positions were invariant: Ser^26^, Val^27^, and Glu^31^. Among these, Ser^26^ and Val^27^ are predicted to be part of the transmembrane region, and Glu^31^ is positioned in the cytoplasmic tail (residues 31–34) of YoaI (table S7). The cytoplasmic domain of EnvZ is essential for its function, whereas the periplasmic and transmembrane domains are dispensable (*66*). Based on this information, we propose a model wherein the conserved Glu^31^ in YoaI’s cytoplasmic tail may mediate binding to EnvZ’s cytoplasmic domain. To assess the importance of Glu^31^ in the YoaI-EnvZ interaction, we created a single alanine substitution (E31A). This mutant YoaI E31A induced weak β-gal activity in a BACTH assay with EnvZ, suggesting that Glu^31^ is critical for this physical interaction (Fig. 8D). To test if the charge on this residue is important for YoaI-EnvZ binding, we substituted Glu^31^ with glutamine (E31Q), which replaces the negatively charged side chain with a neutral amide group. Like E31A, the E31Q mutation disrupted the YoaI-EnvZ interaction (Fig. 8D), implying that Glu^31^ participates in electrostatic interactions with positively charged residues in EnvZ.

We also tested the effect of these mutations on the EnvZ-OmpR–activated *omrB* transcriptional reporter. In contrast to WT YoaI, plasmid expression of the mutants E31A and E31Q failed to stimulate *omrB* transcription (Fig. 8C), indicating that Glu^31^ is critical for YoaI-mediated activation of EnvZ-OmpR signaling. To test whether these mutations (E31A/Q) affected YoaI protein stability or localization to the membrane, we analyzed the membrane fractions of cells expressing the His-tagged forms of WT or mutant YoaI. Both YoaI E31A and E31Q mutants were detected at amounts comparable to that of WT YoaI in the membrane fractions (fig. S8, B and C), indicating that the inability of these mutants to activate *omrB* transcription was not due to protein degradation or mislocalization upon overexpression. Despite their inability to interact with and activate EnvZ, YoaI E31A and E31Q mutants caused a prolonged lag phase similarly to WT when overexpressed (fig. S8D). Together with the finding that *ΔenvZ* cells expressing WT YoaI also have a prolonged lag phase (fig. S7C), we conclude that the effect of YoaI overexpression on cell growth is independent of its role in the EnvZ-OmpR signaling pathway and may involve a separate PhoQ-dependent (Fig. 7C) regulatory mechanism.

## Discussion

Here, we identified 17 small proteins (≤50 amino acids) induced by low-Mg^2+^ stress from approximately 150 documented small proteins in *E. coli*, using Ribo-RET (*40*, *41*). Only 3 of these proteins, MgrB, MgtS, and PmrR were previously studied in the context of Mg^2+^ starvation (*30*, *34*, *36*). For 16 of these stress-induced small proteins, production of the small proteins from the genomic locus was verified in previous studies (table S5), and production of YoaI was verfiied here (Fig. 1, E and F). In a previous report, expression of YoaI tagged with an SPA epitope from a genomic translation fusion was not detected when cells were grown in a rich medium (*21*). Our observation that cells grown in low-Mg^2+^ conditions showed robust production of YoaI-SPA highlights the condition-specific activation of *yoaI* expression (Fig. 1, D to F, Fig. 3, B and C). Using RNA-Seq, we found that many of these stress-induced proteins were transcriptionally activated in low-Mg^2+^ media (Fig. 2A). Further, we identified the regions upstream of these small ORFs responsible for transcriptional regulation using operon fusions and reporter assays. Few hits, including *yoaI* and *yobF*, did not show a change in transcription abundance under low Mg^2+^, suggesting that they are activated post-transcriptionally (Fig. 2C).

Lacking any functional or phenotypic information for most of these stress-induced proteins, we began their characterization by predicting their subcellular localization using bioinformatic tools (Table 1, table S7). By epitope tagging and imaging or western blotting, we empirically determined that 9 of the 17 shortlisted stress-induced small proteins localized to the membrane (Fig. 4, B and D). These membrane-bound small proteins may interact with, stabilize, or fine-tune the activities of large transmembrane proteins, such as sensor kinases, channel proteins, and drug efflux pumps, thereby mediating cellular adaptation and stress responses (*14*). Among these 9 membrane-localized candidates, three proteins—MgrB, MgtS, and PmrR—were previously shown to have specific membrane protein targets (*30*, *34–36*). To further investigate their biological roles, we performed targeted deletion and protein overexpression analysis.

Loss-of-function analysis of small proteins can be a useful approach to understanding their physiological roles. For example, deleting *mgrB* causes hyperactivation of the PhoQP two-component system, leading to cell division inhibition and filamentation of cells grown under low-Mg^2+^ stress (*31*). In clinical isolates of *Klebsiella pneumoniae,* the absence of MgrB-mediated PhoQ inhibition has been linked to acquired colistin resistance (*33*, *67*). In the current work, we found deletion phenotypes associated with the following small proteins—PmrR, YobF, YriA/B, and YqhI— leading to an early entry into the stationary phase and reduced yield at saturation under Mg^2+^ limitation (Fig. 5A). Based on plasmid complementation, growth defects caused by PmrR and YobF were specific to the respective small proteins (Fig. 5, B and C). The overlapping ORFs of YriA and YriB present a unique challenge in interpreting the *ΔyriAB* growth phenotype observed under low-Mg^2+^ stress. Although expressing the native *yriAB* region partially rescued the growth defect, individual expression of YriA or YriB failed to do so; however, modifying YriB by mutating the out-of-frame YriA start codon fully restored growth, emphasizing that the loss of YriB was a critical factor in the observed phenotype (Fig. 5D). In the case of YqhI, the deletion phenotype could not be fully complemented by the plasmid-encoded small protein (Fig. 5E), pointing to putative cryptic ORF or regulatory factors within the small ORF that warrant further investigation.

We also analyzed overexpression phenotypes associated with each of the stress-induced small proteins to gain insights into cellular pathways or components functionally linked to these proteins. For instance, overexpression of MgtS activates the PhoRB two-component system by regulating the phosphate transporter, PitA (*35*). Similarly, DinQ overexpression disrupts membrane potential and depletes intracellular ATP (*50*). We observed overexpression phenotypes for 7 of the 17 small proteins induced under Mg^2+^ stress—MgtS, MgrB, PmrR, YmiA, YoaI, YkgS, and DinQ—resulting in either a long lag phase or slow growth (Fig. 6). These overexpression phenotypes were specific to each small protein, because control plasmids carrying a mutated start codon did not cause growth defects (Fig. 7A). Analysis of cell morphology during the lag and exponential phases of growth revealed variations in cell length and volume that reflect how cells adapt to the overexpression of these small proteins (Fig. 7B, fig. S4). For one of the small proteins, PmrR, either the deletion or overexpression caused growth defects under low-Mg^2+^ stress (Fig. 5A, 6). It is unclear how cells balance small protein abundance when needed during specific conditions versus when not needed, as in the absence of stress. In this context, complementation of the *ΔpmrR* strain with plasmid-encoded PmrR restored normal growth under Mg^2+^ limitation but caused a growth defect under no-stress conditions (Fig. 5B). This suggests that PmrR expression may be precisely controlled in cells undergoing stress, because either its absence or its overexpression disrupts the balance necessary for cellular homeostasis. PmrR may be involved in pathways or regulatory mechanisms in which it interacts with partners other than LpxT. It is not uncommon for small proteins to have multiple binding partners, as seen with MgtS, which interacts with both MgtA and PitA (*34*, *35*). Whether PmrR associates with other targets contributing to phenotypes observed here warrants further investigation.

Finally, we would like to discuss the particular case of the small transmembrane protein YoaI, the transcription of which is activated by the PhoR-PhoB signaling system (*52–54*). YoaI expression is quite low or undetectable when cells are grown in a rich medium or under specific stress conditions (*21*, *22*). However, we found that YoaI protein was detectable under Mg^2+^ limitation and increased with decreasing Mg^2+^ concentrations (Fig. 1, E and F, Fig. 3, B and C). Overexpression of YoaI in WT cells led to a long delay in growth (Fig. 6, 7A). This phenotype was not observed in a *phoQ* mutant (Fig. 7C), hinting at an unknown PhoQ-dependent factor that may contribute to the growth defect observed upon YoaI overexpression. Through a bacterial two-hybrid assay, we found a direct interaction of YoaI with the EnvZ sensor kinase (Fig. 8A), which is involved in osmoregulation (*43*), suggesting that this small protein acts as a connector between the PhoR-PhoB and EnvZ-OmpR signaling networks. Activation of EnvZ by YoaI is supported by the data from our transcriptional reporter analysis of promoters regulated by the EnvZ-OmpR system (Fig. 8B, fig. S7, A and B). We further showed that the physical interaction with EnvZ and its activation were both mediated by a conserved Glu^31^ residue in YoaI (Fig. 8, C and D). The modulation of sensor kinases by small proteins that are induced or activated by a different two-component system appears to be a general theme that may apply to many small membrane proteins (*14*). For instance, under acidic conditions, the 65-amino acid protein SafA is activated by the EvgS-EvgA two-component system and subsequently interacts with and activates the PhoQ-PhoP system in *E. coli* (*68*). In a second example, the 88-amino acid protein PmrD, whose transcription is PhoQP-dependent, stimulates the PmrB-PmrA signaling system in *Salmonella* and *E. coli* (*38*, *69*, *70*). The 127-amino acid protein MzrA, transcriptionally activated by the CpxA-CpxR signaling system, binds to and activates EnvZ, thereby linking the CpxA-CpxR and EnvZ-OmpR pathways in *E. coli* (*63*). Together, our findings outlining the transcriptional regulation of YoaI by PhoB (Fig. 3A), the PhoQ-dependent growth phenotype upon YoaI overexpression (Fig. 7C), and YoaI’s physical interaction with EnvZ and activation of EnvZ-regulated promoter activity (Fig. 8, A to D, fig. S7, A and B) reveal that this 34-amino acid small protein mediates cross-talk between distinct two-component signaling systems – PhoRB, PhoQP, and EnvZ-OmpR to integrate diverse stress responses (Fig. 9, A and B).

**Fig. 9.**
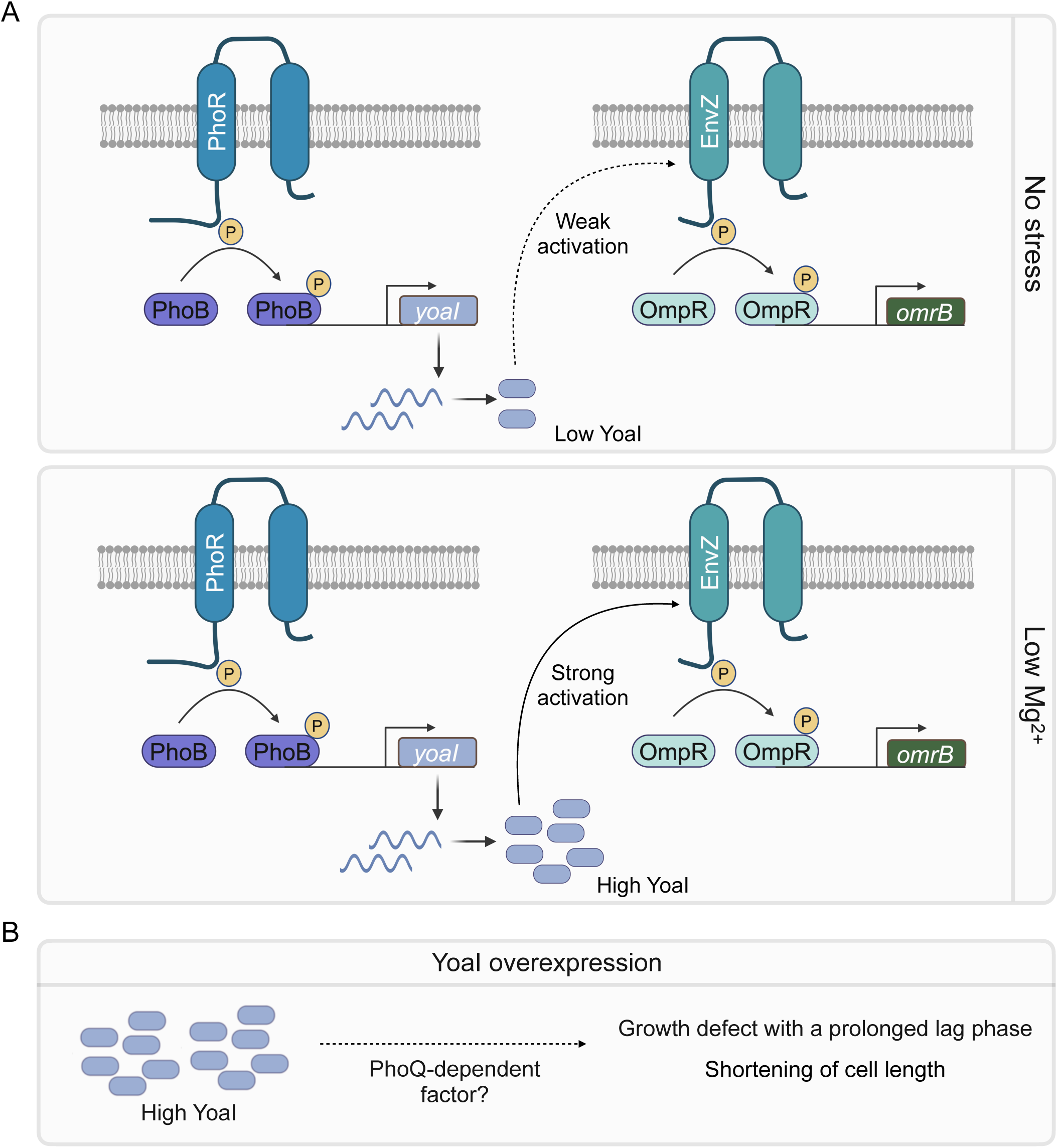
YoaI acts as a connector between two-component signaling systems, PhoR-PhoB and EnvZ-OmpR. (**A**) Small protein YoaI is transcriptionally activated by the PhoR-PhoB system, which is directly stimulated by phosphate starvation and indirectly stimulated by Mg^2+^ starvation. Under low-Mg^2+^ conditions, there is no significant change in *yoaI* transcription, and its expression is activated post-transcriptionally in a PhoQ-independent manner, leading to an increase in YoaI abundance. YoaI binds to the sensor kinase EnvZ and promotes signaling through the EnvZ-OmpR pathway. (**B**) Overexpression of YoaI causes a PhoQ-dependent growth defect that is independent of its action on EnvZ.

Most sORFs (≤150 bp in prokaryotes) are of unknown function, and little is known about when—or even if—they are expressed in the cell and their physiological relevance. Small proteins are thought to fine-tune the activity and/or stability of larger target molecules in the cell, adding a layer of regulation to signaling systems. This type of conditional regulation by small proteins may be especially important when cells are grown in non-ideal or stressful environments. Overall, our work describing the identification of small proteins that are produced condition-specifically and targeted phenotypic characterization provides a deeper understanding of small protein regulation and the physiological consequences of altering their expression, laying a foundation for deciphering their functions and regulatory mechanisms under relevant stress conditions.

## Materials and Methods

### Media and growth conditions

Bacterial cultures were grown in either Luria-Bertani (LB) Miller medium (IBI Scientific) or minimal A medium (*71*) supplemented with 0.2% glucose, 0.1% casamino acids, and the indicated concentration of MgSO_4_, with aeration at 37°C, unless otherwise specified. In general, overnight cultures are grown in supplemented minimal A medium containing 1 mM MgSO_4_ and antibiotic(s) as applicable. Saturated cultures are diluted 1:500 into fresh medium. For low-Mg^2+^ stress, no Mg^2+^ was added when overnight cultures were diluted into fresh medium. Media containing 10 mM Mg^2+^ was used as the no-stress condition. For phosphate stress experiments, MOPS minimal medium (Teknova) supplemented with 0.2% glucose, 0.1% casamino acids, and either 1 mM K_2_HPO_4_ (no-stress) or no K_2_HPO_4_ (phosphate stress). For routine growth on solid medium, LB containing 1.5% bacteriological-grade agar (VWR Lifesciences) was used. For antibiotic selection, carbenicillin, kanamycin, and chloramphenicol were added at a final concentration of 50-100 µg ml^-1^, 12.5-50 µg ml^-1^, and 6-25 µg ml^-1^. For measurement of transcriptional reporter activity, cultures were grown in either supplemented minimal medium or MOPS minimal medium with 6 μg ml^-1^ chloramphenicol in the presence or absence of stress, as indicated. b-isopropyl-D-thiogalactoside (IPTG) was used at a final concentration of 500 µM, unless otherwise specified. Arabinose was used at final concentrations of 0.5% and 30 mM for pBAD24-based vectors and pMR120-based vectors, respectively, to induce protein expression when needed.

### Strains, plasmids, and cloning

See supplemental information for details of the strains (table S1), plasmids (table S2), and primers (table S3) used in this study. All the strains used were derived from *Escherichia coli K-12* MG1655. Strain JW2162, JW3460 and JW3367 were gifts from Dr. Bryce Nickels; plasmids pEB52, pAL38, pAL39, pSMART, pMR120 and strains AML67, MDG147, TIM92, TIM100, and TIM202 were gifts from Dr. Mark Goulian. Strains GSO195, GSO219, GSO225, GSO232, and GSO317 were gifts from Dr. Gisela Storz. Strains JM2110 and JM2113 were gifts from Dr. Maude Guillier.

A modified version of the method described by (*72*) was used to generate in-frame gene deletions corresponding to each small protein. Briefly, the entire small open reading frame (sORF) (including the stop codon) was deleted from the MG1655 genome and substituted with a kanamycin resistance cassette. While constructing the gene deletions, we avoided deleting any overlapping annotated ORFs, non-coding RNAs, or putative regulatory regions in the vicinity of the target gene. In the case of *yriA-yriB (yriAB),* the two small protein ORFs overlap; therefore, we deleted the combined *yriAB* ORF. Detailed information on the genomic coordinates of deleted regions can be found in table S1. Genomic deletions and reporter constructs carrying the kanamycin resistance cassettes were transferred between strains using P1_vir_ transduction (*71*) and validated by PCR. When necessary, the kanamycin cassette was excised from the chromosome by FLP-mediated recombination using pCP20, as previously described (*73*). PCR and loss of antibiotic resistance confirmed the excision of the kanamycin cassette.

Plasmids expressing translational fusions of GFP-A206K to the small open reading frames (sORFs), specifically pPJ1-13 and pSV25, were generated through inverse PCR (*74*) using pAL39 as a template. Plasmids encoding C-terminally 6xHis-tagged-YobF, YoaI and PmrR (sORF-GGSG linker-6xHis) were created using inverse PCR with pEB52 as a template, resulting in pSV14, pSV44 and pSV26, respectively. Plasmids encoding untagged small proteins (pSV30-pSV37, pSV58, pSV60, and pJS1-pJS7) were also generated by inverse PCR with pEB52 as a template. pSV54 was generated by inverse PCR using pBAD24 as a template. Plasmids pSV38-pSV45, pSV55, pSV59, and pSV61, encoding variants of sORFs with start codon substitutions to stop codons (TAA), were generated by site-directed mutagenesis with inverse PCR using templates pSV35, pSV34, pJS7, pJS4, pSV36, pJS5, pSV54, pSV37, and pSV60 as applicable. For pSV39-pSV42, another in-frame start codon within five amino acids of the annotated start was also substituted to TAA. Plasmids pSV70, pJS13, pJS15-16, pJS18 and pJS24 were generated by site-directed mutagenesis with inverse PCR using templates pSV34, pSV47, and pSV44 as applicable (see table S2 for details). All constructs were verified by Sanger sequencing. To generate pSV47, *yoaI* was PCR amplified from the MG1655 genome, digested with *XbaI/EcoRI* restriction enzymes, and then cloned into pUT18C at *XbaI/EcoRI* sites.

To examine the transcriptional regulation of the sORFs and identify the putative regulatory regions, ∼200-500 bp upstream region of the start codon of each sORF was selected for cloning (see details in table S6). If a given sORF was suspected to be part of an operon, two putative regions were chosen for analysis, the first directly upstream of the gene and a second region upstream of the entire operon. Each DNA segment was amplified from MG1655 genomic DNA by PCR and inserted upstream of the *yfp* reporter gene in pMR120 into the restriction sites (*XhoI/BamHI* to construct pSV16, while *EcoRI-BamHI* sites were used for the remaining constructs). The pMR120 vector also includes a *cfp* reporter activated by a constitutive *tetA* promoter, serving as an internal plasmid control. The resulting plasmids (pPJ14, pPJ16-pPJ20, pPJ23, pSV16-21, pSV24, pSV28, pSV29) were confirmed by Sanger sequencing and transformed into MG1655 and TIM202 (*ΔphoQ)*. Note that pSMART (Lucigen), pMR120, and the derivatives are single-copy plasmids when transformed into *E. coli* (*75*). *E. cloni* replicator cells (Lucigen) are transformed with these plasmids and cultured with arabinose induction to increase the copy number for plasmid isolation.

### Translation initiation profiling (Ribo-RET) and RNA-Seq

Overnight cultures of WT *E. coli* grown in supplemented minimal A medium and 1 mM MgSO_4_ were diluted 1:500 in 100 ml of fresh medium and the specified concentrations of MgSO_4_. Once the cultures reached an OD600 of ∼0.4, they were split into two 50 ml conical tubes. Cells from one tube were pelleted and submitted to Genewiz for RNA-Seq, while the cells from the second tube were used for Ribo-RET. For Ribo-RET, cells were treated with retapamulin at a final concentration of 100 µg ml^-1^ for 5 minutes at 37°C with continuous shaking. After retapamulin treatment, cells were filtered through a nitrocellulose membrane with a pore size of 0.45 µm, quickly scraped off the surface, and flash-frozen in liquid nitrogen. The resulting samples were processed, and libraries were prepared as described previously (*44*) with the following modifications. Specifically, RNA fragments of 15-30 nucleotides (nt) in length were isolated by performing a tight gel size selection, and rRNA depletion was performed using the RiboCop for Gram-negative bacteria kit (Lexogen). The resulting RNA fragments were then pooled, and libraries were prepared, with a final library structure consisting of a 5’ adapter – 4 random bases – insert – 5 random bases – sample barcode – 3’ adapter. The randomized bases serve as Unique Molecular Identifiers (UMIs) for deduplication.

The multiplexed library was sequenced on an Illumina HiSeq 4000 with a paired end reading of 150 bases (PE150 runs) at a sequencing depth of 40-60 million raw reads per sample.

### Sequencing data analysis

The raw sequencing data for this project has been deposited in the GEO database with the accession number GSE276379. An R package with codes for data processing and analysis has been made available on GitHub (https://github.com/yadavalli-lab/Small-proteins-induced-under-low-magnesium-stress-in-E.coli). An archived version of the code for the current paper is deposited at Zenodo (DOI <u>10.5281/zenodo.15597036</u>). The raw sequencing data was first demultiplexed, and the adaptors were removed using cutadapt (*76*). The reads were then depleted of rRNA and tRNA and deduplicated using umi_tools dedup (*77*). The Ribo-RET reads were aligned to the *E. coli* MG1655 genome NC_000913.3 (RefSeq assembly accession: GCF_000005845.2) using hisat2 (*78*), and the ribosome density was assigned to the 3’ end of the reads. Initiation peak density was calculated for the annotated translation start sites by summing up the normalized reads (reads per million, RPM) within 4 to 20 nt downstream of the first nucleotide in the start codon. For transcript quantification of RNA-Seq reads, Kallisto (*79*) was used. DESeq2 (*80*) was used for the differential expression analysis of gene expression in the RNA-Seq data. Normalization of the data for differential expression was carried out using the “apeglm” method (*81*). Fold-changes were estimated by comparing two replicates of the low-Mg^2+^ dataset to two replicates of the no-stress dataset. The criteria for significant changes include a fold change threshold of >2 and a p-value (FDR) of <0.05.

### Measurement of growth (optical density) and fluorescence using a microplate reader

The optical density (OD_600_) measurements for small protein overexpression and deletion constructs were performed in a clear flat bottom 96-well microplate (Corning) containing 150 µL culture per well using a microplate reader (Agilent BioTek Synergy Neo2S). Fluorescence measurements for transcriptional reporters were carried out similarly, using black-walled, clear-flat bottom 96-well microplates (Corning). Saturated cultures grown in supplemented minimal A or MOPS minimal medium were normalized to the same initial OD_600_ values across all samples and diluted 1:50 in fresh medium containing specified concentrations of MgSO_4_ or K_2_HPO_4_, along with appropriate antibiotics and inducers, as indicated. For growth curves of small protein-encoding gene deletions, cultures were grown in supplemented minimal A medium with no added Mg^2+^ (low Mg^2+^). For plasmid complementation, culture media included 50 μg ml^-1^ carbenicillin, and either no added Mg^2+^ or 10 mM MgSO_4_ as indicated. 500 µM IPTG was used for induction, except for PmrR derivatives (pSV35 and pSV38), where 12.5 µM IPTG was used. For plasmid overexpression of small proteins, cultures were grown in supplemented minimal A medium with no added Mg^2+^ and 50 μg ml^-1^ carbenicillin. 500 µM IPTG was used as an inducer for all P*_trc_*-based vectors, in the case of pBAD24 and its derivative pSV54, 0.5% arabinose was used. To generate growth curves of *ΔlpxT* and *ΔpmrR* strains, cells were grown in supplemented minimal A medium with no added Mg^2+^. For plasmid complementation of the *ΔlpxT* strain, cultures were grown in supplemented minimal A medium containing 50 μg ml^-1^ carbenicillin, 500 µM IPTG, and either 10 mM MgSO_4_ or no added Mg^2+^. For growth curves of MG1655 WT or *ΔenvZ* strains complemented with plasmid-encoded YoaI (or its variants), cultures were grown in supplemented minimal A medium containing no added Mg^2+^, 50 μg ml^-1^ carbenicillin, and 500 µM IPTG. The microplates were incubated at 37 °C with double orbital shaking before the real-time measurements of optical density (at 600 nm), YFP (excitation: 500 nm; emission: 540 nm), or CFP (excitation: 420 nm; emission: 485 nm) were acquired every hour for 20-24 hours.

### Microscopy and image analysis

For localization of GFP-tagged small proteins, cells were cultured in supplemented minimal A medium containing 1 mM Mg^2+^ and 50 μg ml^-1^ carbenicillin to an OD_600_ of ∼0.2 (∼3 hours) and then rapidly cooled in an ice slurry. Streptomycin was added to a final concentration of 250 µg ml^-1^. GFP fluorescence was captured with a 475 nm excitation wavelength and a 540 nm emission filter, with an exposure time of 30 ms and 20% intensity. While the exposure times were typically set to these specified values, slight variations were occasionally made to optimize imaging based on the fluorescence intensities observed.

For overexpression analysis, cultures were grown in supplemented minimal A medium with no added Mg^2+^. For pSV34-36, pSV38-42, pSV60-61, pJS4, and pJS7, the medium contained 50 μg ml^-1^ carbenicillin and 500 µM IPTG, while for pSV54-55, 0.5% arabinose was utilized. Cells were sampled in the lag phase (OD_600_ ∼0.005), then rapidly cooled in an ice slurry before phase contrast microscopy. For empty vector control and start codon variants, we captured lag phase micrograph images approximately 30-40 minutes post-inoculation. For MgtS, MgrB, PmrR, YoaI, and YmiA overexpression, lag phase measurements were performed at 2-3 hours, and for YkgS and DinQ, at 1-hour post-inoculation. To concentrate the cells, 1 mL of the culture was pelleted and resuspended in 30-50 µL minimal A salt solution. For exponential phase analysis (OD_600_ between 0.2-0.4) in cultures carrying empty vector control and start codon variants images were taken at ∼3-4 hours. In the case of cells expressing MgtS, MgrB, PmrR, YoaI, and YmiA, images were taken at 16-18 hours, and YkgS and DinQ at 6-8 hours. For the analysis of cell size changes in the *ΔphoQ* strain, cells were grown in supplemented minimal A medium with no added Mg^2+^ and 50 μg ml^-1^ carbenicillin.

Microscope slides were prepared using 1% agarose pads as described previously (*31*), with approximately 5-7 µL of the resuspended cells. A 40 ms exposure time and 20% intensity were applied to capture the phase-contrast images.

For single-cell fluorescence measurements of WT and *ΔyoaI* cells carrying P*_omrB_*-*mScarlet* reporter, overnight cultures were diluted 1:500 into fresh supplemented minimal A medium containing either 10 mM or no added MgSO_4_ for Mg^2+^ stress experiments. Similarly, for phosphate stress, cultures were back-diluted in MOPS minimal medium containing either 1 mM K_2_HPO_4_ or no K_2_HPO_4._ Cells were grown to stationary phase (OD_600_ ∼1.0 for stress condition and ∼3.5 for no stress condition), and then rapidly cooled in an ice slurry. Streptomycin was added to a final concentration of 250 µg ml^-1^. Fluorescence was measured using the mCherry channel with an excitation wavelength of 570 nm and an emission filter of 645 nm. The exposure time was set to 100 ms, with an intensity of 20%. Background fluorescence was determined by imaging MG1655 cells grown under the same conditions.

For single-cell fluorescence measurements upon YoaI overexpression (to measure reporter activities of P*_ompF_*-*yfp* and P*_ompC_*-*cfp*), the overnight cultures were diluted 1:500 into fresh supplemented minimal A medium containing 10 mM MgSO_4_. Cells were grown to exponential phase (OD_600_ between 0.2-0.4), then rapidly cooled in an ice slurry, and streptomycin was added at 250 µg ml^-1^.

YFP fluorescence was measured using a 500 nm excitation wavelength and a 535 nm emission filter. CFP fluorescence was measured using a 435 nm excitation wavelength and a 480 nm emission filter. The exposure time for YFP and CFP measurements was set to 50 ms, respectively, with an intensity of 20%. The background fluorescence was determined by imaging MG1655/pEB52 cells grown under the same conditions.

Image visualization and acquisition were performed using a Nikon Ti-E epifluorescence microscope equipped with a TI2-S-HU attachable mechanical stage. The images were captured with a Prime 95B sCMOS camera from Teledyne Photometrics with 1×1 binning. All image acquisition was managed using Metamorph software (Molecular Devices), version 7.10.3.279. A minimum of 50 cells per replicate were analyzed for cell volume and single-cell fluorescence quantification. The distributions of cellular volume were assumed to follow the sphero-cylindrical shape of *E. coli.* The volume was calculated using the formula 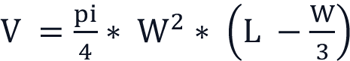. Images were analyzed, and the cellular length, width, and fluorescence intensity were quantified using ImageJ (*82*) and MicrobeJ plugin (*83*).

### Preparation of membrane and cytoplasmic fractions

Membrane and cytoplasmic protein fractions were prepared as described previously (*29*). Briefly, saturated cultures of MG1655/pSV14, MG1655/pSV26, MG1655/pSV44, MG1655/pJS18, MG1655/pJS24 and MG1655/pEB52 (empty vector) were diluted 1:500 in 4 ml of LB media containing 100 µg ml^-1^ carbenicillin and grown at 37 °C. After 4 hours of growth, 0.5 mM IPTG was added to the cultures, and the cells were harvested after 2 hours of induction. Cell pellets were resuspended in 50 µl of cold resuspension buffer containing 20% sucrose, 30 mM Tris pH 8.0, and 1X protease inhibitor cocktail (Sigma, cOmplete, EDTA-free). Then, 50 µl of 10 mg ml^-1^ lysozyme, freshly prepared in 0.1 M EDTA pH 7.0, was added to the cell suspension, and the mixture was incubated on ice for 30 minutes. Next, 1 ml of 3 mM EDTA pH 7.5 was added to each sample, followed by sonication (10s pulse, 10s gap, 6 times) using VCX-130 Vibra-Cell Ultrasonic sonicator (Thermo Fisher Scientific). Samples were briefly centrifuged at 6,000 rpm for 10 minutes at 4 °C to remove cellular debris, and the supernatant was collected and spun at 21,000 ×g for 30 minutes at 4°C. The supernatant from this step is separated and stored as the cytoplasmic fraction. The pellet, representing the membrane fraction, was resuspended in a storage buffer containing 20 mM Tris pH 8.0/20% glycerol. For strains carrying genomic YoaI-SPA translational fusion, cultures were grown to exponential phase (OD_600_ of ∼0.2-0.4) before harvesting, and membrane and cytoplasmic fractions were prepared as mentioned above.

### Western blot analysis

The membrane and cytoplasmic fractions were resuspended in 4X Laemmli sample buffer containing 5% 2-mercaptoethanol and no dyes. The samples were heated at 70 °C for 10 minutes and run on a 12% Bis-Tris gel (NuPage, Invitrogen) using MES running buffer (Invitrogen) at 160V for 80 minutes. After electrophoresis, proteins were transferred to a PVDF membrane (Amersham^TM^ Hybond^TM^) with a 0.2 µm pore size using a semi-dry transfer cell (Bio-Rad Trans-Blot SD). The membranes were blocked with 5% w/v milk in Tris-buffered saline, pH 7.4, with 1% Tween 20 (TBS-T). Primary antibodies – mouse M2 anti-FLAG (Sigma-Aldrich, Cat# F1804) and rabbit anti-6xHis (Rockland, Cat# 600-401-382) were used at 1:1000 dilution to detect SPA- and 6xHis-tagged constructs, respectively. IRDye 800CW goat anti-mouse, part number: 926-32210 and IRDye 680RD donkey anti-rabbit, part number: 926-68073 antibodies (LiCOR) were used for secondary detection. SeaBlue^TM^ Plus2 Pre-stained Protein Standard (Invitrogen) was used as a ladder to visualize lower molecular weight protein bands. The proteins were visualized using a LI-COR Odyssey imager.

### Bacterial two-hybrid and β-galactosidase reporter gene assays

For the bacterial two-hybrid (BACTH) assay, a *cyaA* mutant host strain (SAM85) carrying *lacI*^q^ and T18 and T25 fusion proteins of *B. pertussis* adenylyl cyclase was used. Several clones from the transformation plate were picked to reduce heterogeneity (*61*) and inoculated in LB containing 100 µg ml^-1^ carbenicillin and 50 µg ml^-1^ kanamycin. Cultures were grown overnight at 30°C with shaking. The cultures were then back-diluted 1:1000 into LB supplemented with antibiotics and 500 µM IPTG. Cells were grown at 30°C until OD_600_ reached between 0.2-0.4, after which β-galactosidase activities (Miller units) were measured as previously described (*84*).

For the β-galactosidase reporter gene assay, strains carrying P*_omrB_*-*lacZ* with pSV34, pSV70, pJS13 or pEB52 (empty vector) were grown overnight in LB medium with 100 µg ml^-1^ carbenicillin. The next day, the cultures were diluted 1:100 in the same medium and grown at 37 °C. After 3 hours of growth, 0.5 mM IPTG was added to the cultures, and the cells were harvested after 2 hours of induction. For high-throughput measurements in a 96-well format, the protocol was modified as previously described (*84*).

### Multiple Sequence Alignment of YoaI

The protein sequences of YoaI orthologs from Gammaproteobacteria were obtained from the NCBI protein database using Entrez queries. To minimize redundancy, only one representative sequence per species was retained, and sequences exceeding 40 amino acids in length were excluded.

Multiple sequence alignments were performed using ClustalW (*85*, *86*), and results were visualized together with generating a sequence logo of conserved residues using msaPrettyPrint package from R (*87*), applying a 50% identity threshold to highlight conserved regions.

### Statistical Analysis

Unless otherwise stated, statistical comparisons of experimental results from independent experiments across conditions were performed using an unpaired two-tailed Student’s *t* test; a P value < 0.05 was considered statistically significant. Details of the sample sizes, number of replicates, and statistical test being used are described in each figure legend in the main text and Supplementary Materials.

## Supplementary Materials

Figs. S1 to S8

Tables S1 to S7

References (88) to (96)

## Acknowledgements

The authors would like to thank Drs. Mark Goulian, Bryce Nickels, Gisela Storz, and Maude Guillier for generously sharing strains and plasmids. We thank Johnathan Akdemir for his help with plasmid preparations during the cloning and construction of YoaI mutants. We thank Dr. M. Goulian, and the past and present members of the Yadavalli, Storz, and Shah Labs for their input and helpful discussions. **Funding:** S.S.Y. is supported by the National Institutes of Health - National Institute of General Medical Sciences (NIH-NIGMS) ESI-MIRA R35 GM147566 and institutional start-up funds from Rutgers. P.S. was supported by NIH-NIGMS grant R35 GM124976 and start-up funds from the Human Genetics Institute of New Jersey at Rutgers University. S.V. is supported by the Waksman Institute Busch Predoctoral fellowship (2022–24). The funders did not play any role in the study design, data collection and analysis, decision to publish, or preparation of the manuscript. **Author contributions:** S.V., P.S., and S.S.Y. conceived the experiments. All authors planned the experiments, designed the methodology, and performed data analysis. S.V., P.S., and S.S.Y. wrote the initial draft of the manuscript, and all authors reviewed the final manuscript. **Competing interests:** P.S. is a Director at an RNA-therapeutics startup. The other authors declare that they have no competing interests. **Data and materials availability:** Next-generation sequencing data generated in this study are deposited at Gene Expression Omnibus (GEO) with accession number GSE276379. The code used for data processing and analysis is available in a series of R Markdown documents hosted on GitHub (https://github.com/yadavalli-lab/Small-proteins-induced-under-low-magnesium-stress-in-E.coli). An archived version of the code for the current manuscript is deposited at Zenodo (DOI 10.5281/zenodo.15597036). All other data needed to evaluate the conclusions in the paper are present in the paper or the Supplementary Materials. The strains and plasmids generated in this study are available from the lead contact, Srujana S. Yadavalli, under a material transfer agreement with Rutgers University.

