## Supplementary Figures and Tables for "Analysis of stress-induced small proteins in *Escherichia coli* reveals that YoaI mediates cross-talk between distinct signaling systems"

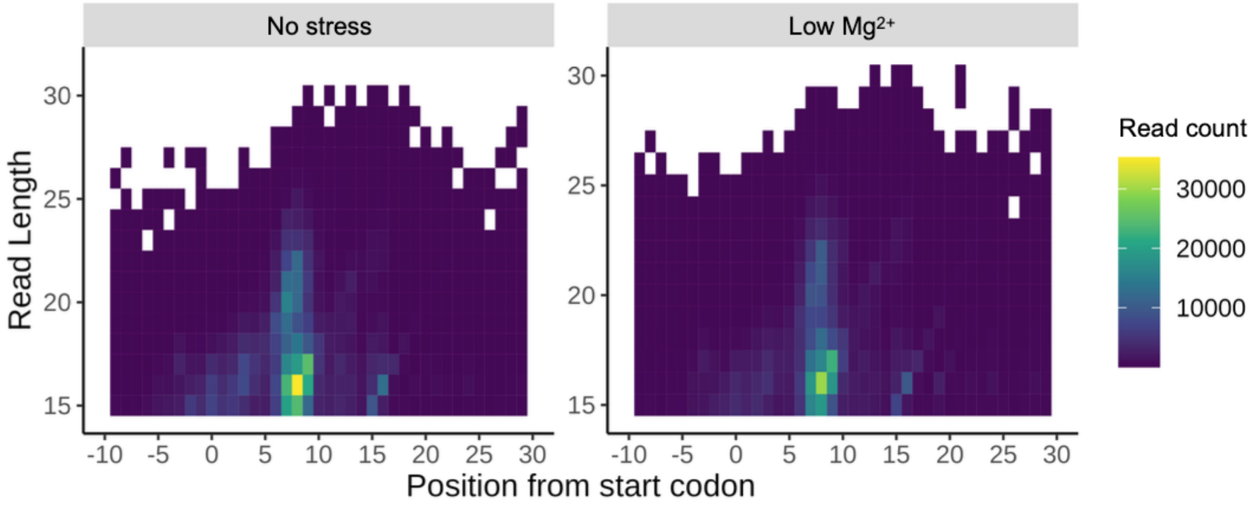


**Fig. S1. Ribogrid analysis representing Ribo-RET reads aligned to the genome.** The rows represent reads of varying lengths, and the columns indicate the position of the 3'-end of a footprint, n = 1.

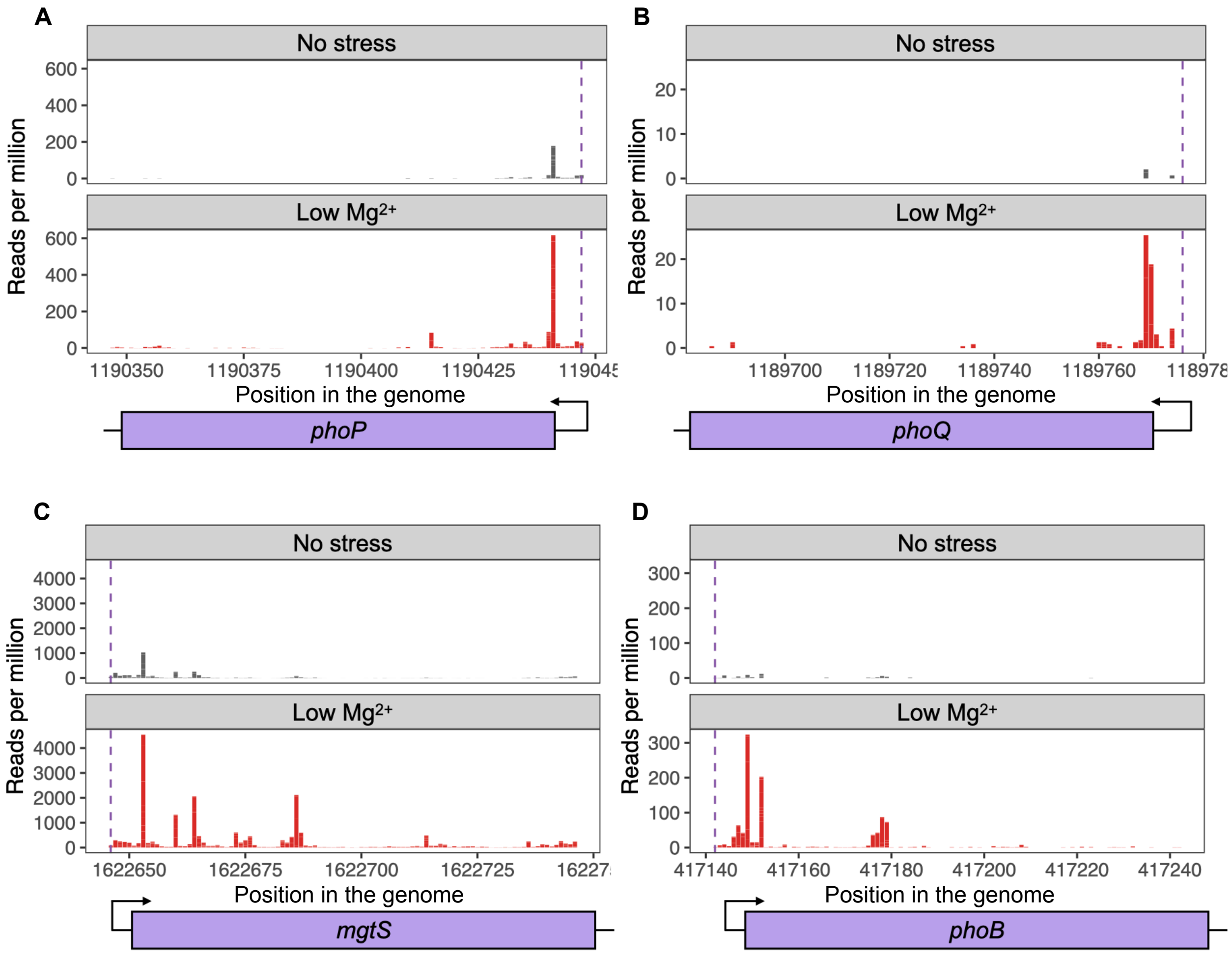


**Fig. S2. Validation of Ribo-RET in detecting ribosome occupancy on annotated translation start sites of proteins known to be induced under low-$Mg^{2+}$ stress.** Ribo-RET reads mapping to the translation start site of (**A**) PhoP, (**B**) PhoQ, (**C**) MgtS, and (**D**) PhoB under no-stress condition and low-$Mg^{2+}$ stress. n = 1. A purple dashed line indicates the translation start site.

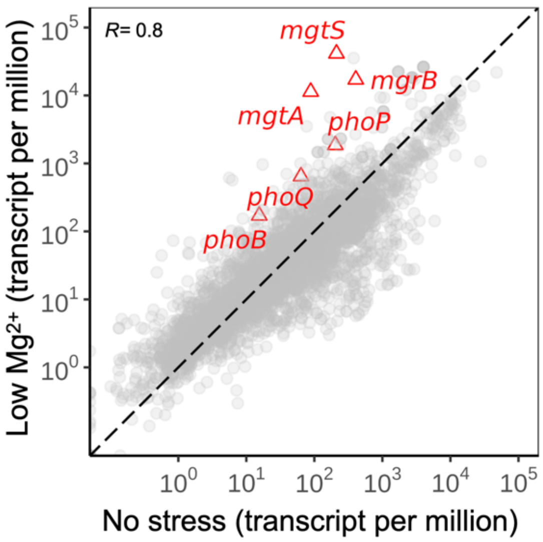


**Fig. S3. Validation of RNA-Seq in detecting transcripts known to be induced under low-$Mg^{2+}$ stress.** Scatterplot showing the correlation between the normalized RNA-Seq reads mapping to all the annotated genes under low-$Mg^{2+}$ stress and no stress. *mgtS, mgrB, phoP, phoQ*, *mgtA,* and *phoB,* known to be induced under $Mg^{2+}$ starvation, are highlighted in red triangles; the rest of the transcripts are shown as gray circles. Pearson's coefficient, r = 0.8. Data is derived from the average of n = 2 biological replicates.

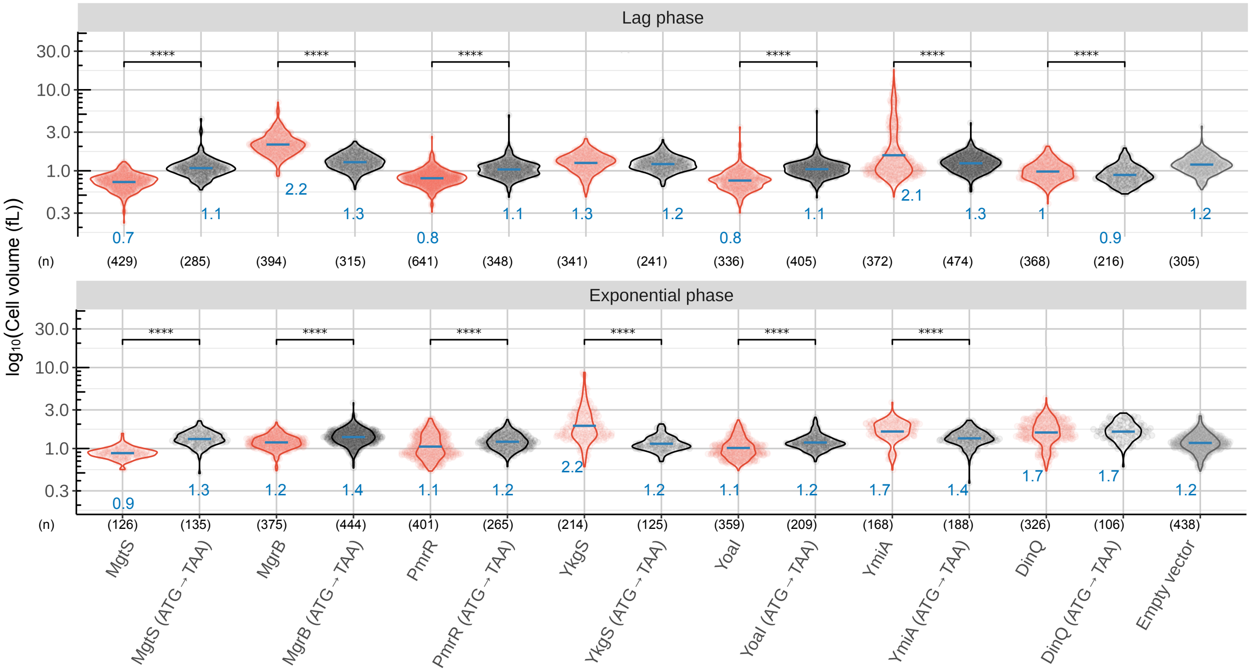


**Fig. S4. Quantification of cell size changes associated with overexpression of the small proteins MgtS, MgrB, PmrR, YkgS, YoaI, YmiA, and DinQ.** Cell volume quantification (representative micrograph images are shown in Fig. 6B) for the cells expressing small proteins during lag (top) and exponential (bottom) phases after induction is shown. Each circle corresponds to a single cell. Cells expressing WT small proteins are shown in red, and cells expressing the start codon mutants or empty vector (pEB52) are shown in gray, n ≥ 106 cells per strain from two biological replicates per condition. The numbers under the x-axis specify the number of cells measured in each group. Mean cell volumes are indicated with blue bars and numbers. P-values indicate the results of a t-test when the strains expressing a small protein were compared to their variant with the translation start codon mutated, ****P ≤ 0.001.

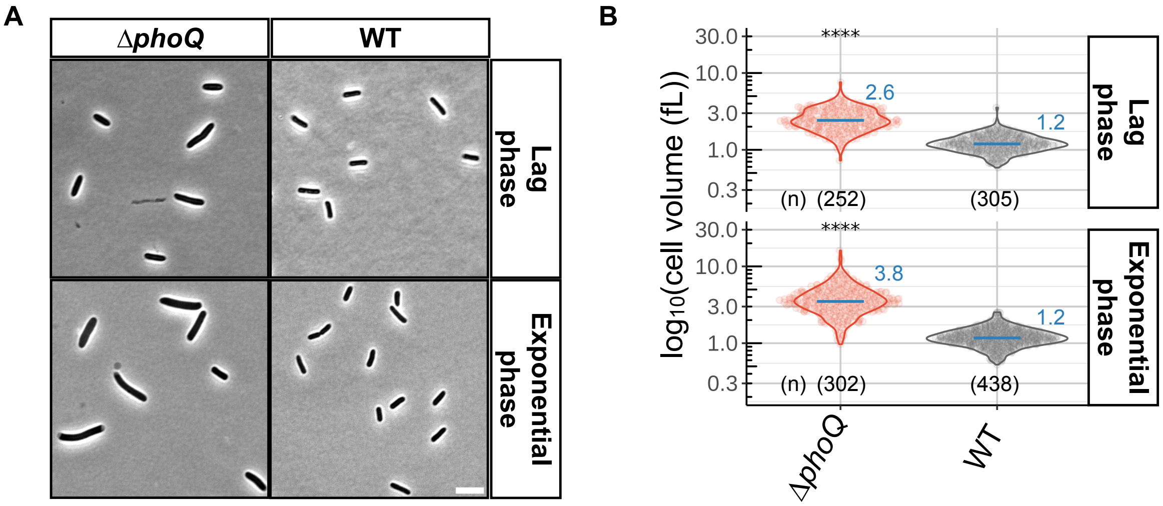


**Fig. S5. Cell size changes associated with Δ*phoQ* under $Mg^{2+}$ starvation.** (**A**) Representative phase contrast micrographs of Δ*phoQ* (TIM202) and WT (MG1655) *E. coli* expressing an empty vector (pEB52) during lag and exponential phases, n = 2 biological replicates per strain. Scale bar, 5 µm. Please note that the control image in panel S5A is identical to that used in Fig. 7B, and the experiments were conducted side by side with the same controls. (**B**) Quantification of cell volume for the cells depicted in (A). Each circle corresponds to a single cell. n ≥ 252 cells per strain from 3 biological replicates per condition. Mean cell volumes are shown with blue bars and text. P-values indicate the results of a t-test when Δ*phoQ* cells were compared to the wild-type cells, ****P ≤ 0.0001.

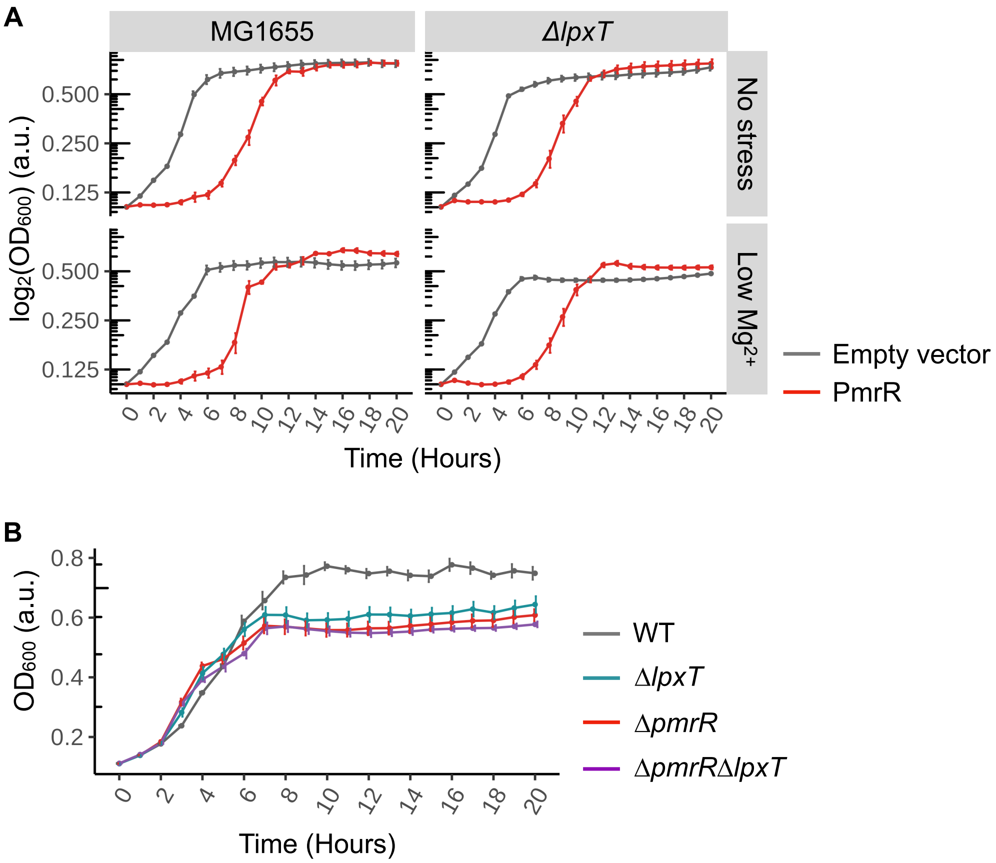


**Fig. S6. Role of LpxT in the growth phenotypes associated with PmrR.** (**A**) Growth comparison of *E. coli* K-12 MG1655 WT and Δ*lpxT* strain (SV47) carrying a plasmid encoding the small protein PmrR (pSV35) or an empty vector (pEB52). (**B**) The plot represents growth curves of WT *E. coli* MG1655 and mutants corresponding to gene deletions Δ*pmrR* (SV35), Δ*lpxT* (SV47), or Δ*pmrR* Δ*lpxT* (SV54). All data represent averages and standard errors of means for n = 4 biological replicates per strain.

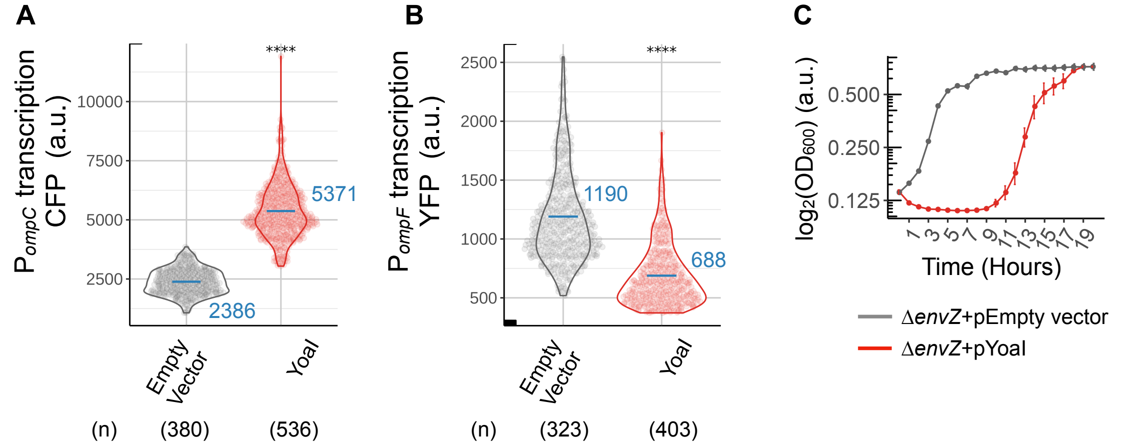


**Fig. S7. Effect of YoaI overexpression on the EnvZ-OmpR system.** (**A and B**) Expression of promoter reporters for genes encoding porins. CFP fluorescence driven by the *ompC* promoter (A), and YFP fluorescence driven by the *ompF* promoter (B) were measured during the exponential growth phase in WT (MDG147) cells overexpressing YoaI (pSV34) or an empty vector (pEB52). Each circle corresponds to a single cell. n ≥ 323 cells per strain from three independent replicates per condition. Mean fluorescence is shown in blue bars and text. P-values indicate the results of a t-test when cells overexpressing YoaI were compared to the empty vector control, ****P ≤ 0.0001. (**C**) Growth curves of Δ*envZ* (SV98) cells overexpressing YoaI (pSV34) or an empty vector (pEB52). Data represent averages and standard error of means for n = 4 biological replicates per strain.

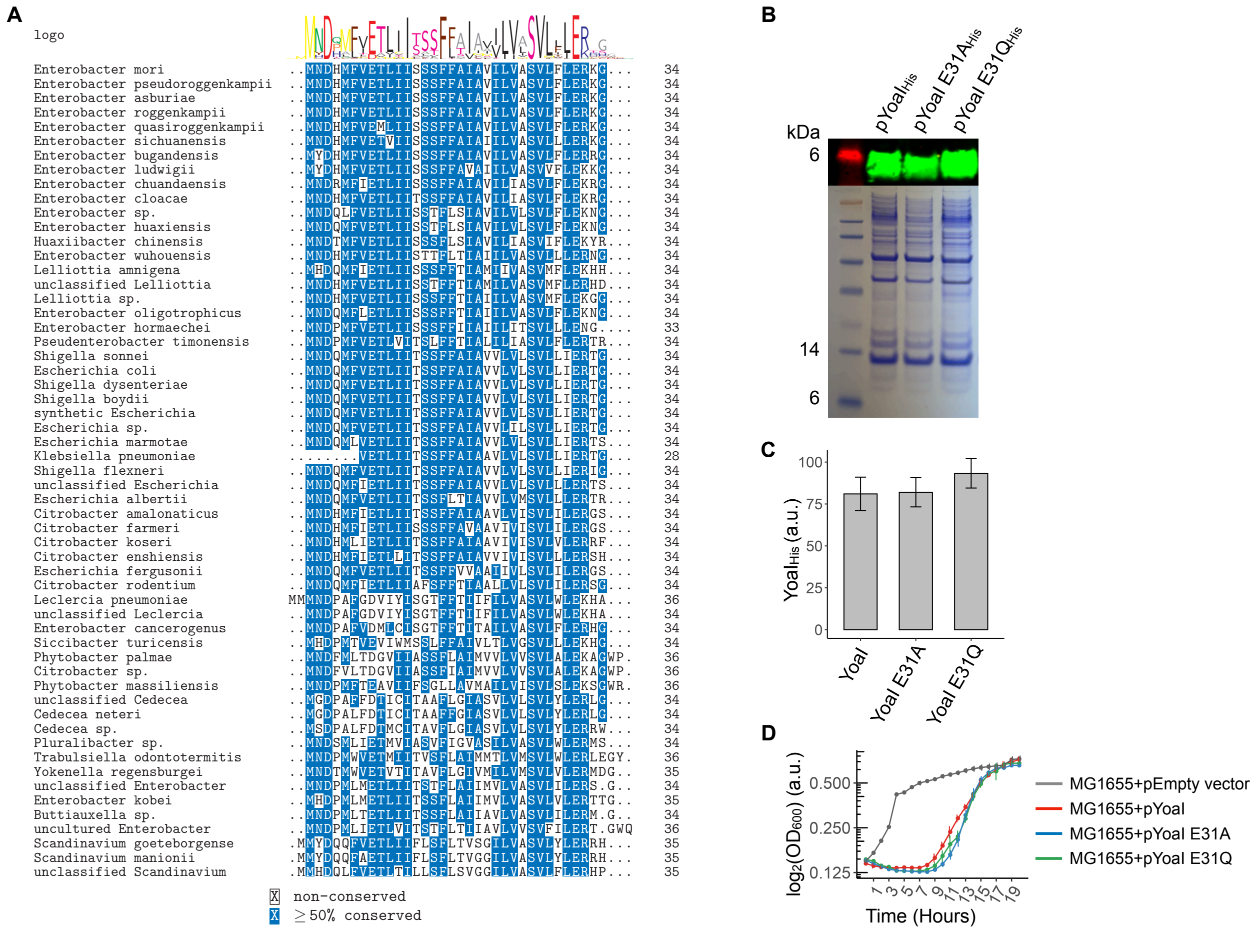


**Fig. S8. Mutagenesis and expression of YoaI mutants.** (**A**) Multiple sequence alignment and consensus amino acid sequence of YoaI from 57 representative Gammaproteobacteria. (**B**) Membrane fractions of *E. coli* K-12 MG1655 cells expressing YoaI-6XHis (pSV44), YoaI E31A-6XHis (pJS18), or YoaI E31Q-6XHis (pJS24) were analyzed by western blotting using an antibody that recognizes His and by Coomassie Brilliant Blue staining. The data represent results from n = 3 biological replicates per strain. (**C**) Quantification of YoaI-His abundance from the western blot in (B). The data represent mean ± standard error of the mean for n = 3 biological replicates per strain. P-values were calculated using a t-test comparing WT YoaI to its mutant derivatives, with no statistically significant differences observed ($P > 0.05$). (**D**) Growth curves of *E. coli* MG1655 cells expressing WT YoaI (pSV34), YoaI E31A (pSV70), YoaI E31Q (pJS13), or an empty vector (pEB52). Data represent averages and standard error of mean for n = 4 biological replicates per strain.

**Table S1. List of strains used in the study.**

| Strain | Genotype | Source or Reference |
|---|---|---|
| AML67 | MG1655 *ΔmgrB*::(FRT-kan-FRT) *ΔlacZ* $\lambda_{att}$::($P_{mgtA}$-*lacZ cat*) | (*30*) |
| *E. cloni* | F⁻ *mcrA Δ(mrr-hsdRMS-mcrBC) endA1 recA1 φ80lacZΔM15 ΔlacX74 araD139 Δ(ara-leu)7697 galU galK rpsL nupG tonA (attL araC-PBAD-trfA250 bla attR) λ−* | Lucigen Corporation |
| *E. coli K-12 MG1655* | F- λ- ilvG- rfb-50 rph-1 | *E. coli* Genetic Stock Center, CGSC# 7740 |
| GSO195 | MG1655 *ΔdinQ*::(FRT-kan-FRT) | (*49*) |
| GSO219 | MG1655 *ΔydgU*::(FRT-kan-FRT) | (*49*) |
| GSO225 | MG1655 *ΔymiA*::(FRT-kan-FRT) | (*49*) |
| GSO232 | MG1655 Δ*yoaI*::(FRT-kan-FRT) | (*49*) |
| GSO317 | MG1655 *yoaI*-SPA::kan | (*22*) |
| JM2110 | MG1508 *mhpR*-$P_{LtetO\text{-}1\text{-}omrB\text{-}104+8}$-*lacZ*$_{+28}$ *Δmini-λ-Tet* | Guillier lab |
| JM2113 | OK510 *argG*-[TT1-$P_{LtetO\text{-}1\text{-}omrB\text{-}104+8}$-mScarlet$_{+4}$-FRT-*nptII*-FRT-TT2]-*yhbX*, *Δmini-λ-Tet* | Guillier lab |
| JW2162 | BW25113 Δ*lpxT*::(FRT-kan-FRT) | (*88*) |
| JW3367 | BW25113 Δ*envZ*::(FRT-kan-FRT) | (*88*) |
| JW3460 | BW25113 Δ*phoB*::(FRT-kan-FRT) | *(84)* |
| MDG147 | MG1655 (seq) Φ(*ompF*$^+$-*yfp*$^+$)30 Φ(*ompC*$^+$-*cfp*$^+$)31 | (*65*) |
| MG1508 | MG1655 *mal::lacI*$^q$, *mini- λ-Tet*, *mhpR*-$P_{LtetO\text{-}1}$-*cat-sacB-lacZ* | (*89*) |
| OK510 | MG1432 *argG*-[TT1-$P_{LtetO\text{-}}$1(no -10)-*sacB-cat*-mScarlet(no ATG)- FRT-*nptII*-FRT-TT2]-*yhbX* | (*64*) |
| SAM85 | F+ *lacI*$^q$*ZΔM15* Tn*10*(Tet$^R$), *cya-99 araD139 galE15 galK16 rpsL1* (Str$^R$) *hsdR2 mcrA1 mcrB1*. Strain prepared by conjugation: BTH101 (F-) x XL1-Blue (F+). | (*29*) |
| SV25 | MG1655 *ΔyriAB*::(FRT-kan-FRT), coordinates deleted: 3,640,612 - 3,640,699 | This study |
| SV27 | MG1655 *ΔyadW*::(FRT-kan-FRT), coordinates deleted: 176,552 - 176,594 | This study |
| SV29 | MG1655 *ΔyddY*::(FRT-kan-FRT), coordinates deleted: 1,567,219 - 1,567,178 | This study |
| SV31 | MG1655 *ΔmgtT*::(FRT-kan-FRT), coordinates deleted: 1,622,742 - 1,622,816 | This study |

| | | |
|---|---|---|
| SV33 | MG1655 *ΔyobF*::(FRT-kan-FRT), coordinates deleted: 1,907,451 - 1,907,573 | This study |
| SV35 | MG1655 *ΔpmrR*::(FRT-kan-FRT), coordinates deleted: 4,332,116 - 4,332,178 | This study |
| SV37 | MG1655 *ΔyqhI*::(FRT-kan-FRT), coordinates deleted: 3,147,597 - 3,147,740 | This study |
| SV39 | MG1655 *ΔyadX*::(FRT-kan-FRT), coordinates deleted: 175,048 - 175,077 | This study |
| SV41 | MG1655 *ΔymiC*::(FRT-kan-FRT), coordinates deleted: 1,335,572 - 1,335,667 | This study |
| SV43 | MG1655 *ΔykgS*::(FRT-kan-FRT), coordinates deleted: 290,510 - 290,641 | This study |
| SV47 | MG1655 Δ*lpxT*::(FRT-kan-FRT) | This study |
| SV48 | MG1655 Δ*phoB*::(FRT-kan-FRT) | This study |
| SV50 | MG1655 *ΔpmrR* | This study |
| SV54 | MG1655 *ΔlpxT*::(FRT-kan-FRT) *ΔpmrR* | This study |
| SV56 | MG1655 *ΔdinQ*::(FRT-kan-FRT) | This study |
| SV57 | MG1655 *ΔymiA*::(FRT-kan-FRT) | This study |
| SV58 | MG1655 *ΔydgU*::(FRT-kan-FRT) | This study |
| SV59 | MG1655 *ΔyoaI*::(FRT-kan-FRT) | This study |
| SV60 | MG1655 *yoaI*-SPA-kan *ΔlacZ ΔphoQ* | This study |
| SV64 | MG1655 *ΔmgtS*::(FRT-kan-FRT), coordinates deleted: 1,622,646 - 1,622,735 | This study |
| SV91 | MG1655 Δ*yoaI* | This study |
| SV95 | MG1655 *argG*-[TT1-P$_{\text{LtetO-1-}omrB\text{-104+8}}$-mScarlet$_{+4}$-FRT-*nptII*-FRT-TT2]-*yhbX*, *Δmini-λ-Tet* Δ*yoaI* | This study |
| SV98 | MG1508 *mhpR*-P$_{\text{LtetO-1-}omrB\text{-104+8}}$-*lacZ*$_{+28}$ *Δmini-λ-Tet* Δ*envZ::kan* | This study |
| SV99 | MG1655 *argG*-[TT1-P$_{\text{LtetO-1-}omrB\text{-104+8}}$-mScarlet$_{+4}$-FRT-*nptII*-FRT-TT2]-*yhbX*, *Δmini-λ-Tet* | This study |
| SV100 | MG1655 *argG*-[TT1-P$_{\text{LtetO-1-}omrB\text{-104+8}}$-mScarlet$_{+4}$]-*yhbX*, *Δmini-λ-Tet* Δ*yoaI* | This study |
| SV101 | MG1655 *argG*-[TT1-P$_{\text{LtetO-1-}omrB\text{-104+8}}$-mScarlet$_{+4}$]-*yhbX*, *Δmini-λ-Tet* Δ*yoaI* Δ*envZ::kan* | This study |
| TIM92 | MG1655 λ$_{att}$::(P$_{mgrB}$-*yfp*) HK$_{\text{att}}$::(P$_{tetA\text{-}cfp}$) | (*90*) |
| TIM100 | MG1655 *ΔphoQ* λ$_{\text{att}}$::(P$_{mgrB\text{-}yfp}$) HK$_{\text{att}}$::(P$_{tetA\text{-}cfp}$) | (*90*) |

| | | |
|---|---|---|
| TIM202 | MG1655 Δ*lacZ*::FRT Δ*phoQ*::FRT | (*91*) |
| TOP10 | *F– mcrA Δ(mrr-hsdRMS-mcrBC) φ80lacZΔM15 ΔlacX74 recA1 araD139 Δ(ara-leu)7697 galU galK λ– rpsL(StrR) endA1 nupG* | Invitrogen |
| XL1-Blue | *recA1 endA1 gyrA96 thi-1 hsdR17 supE44 relA1 lac [F´ proAB lacIqZΔM15 Tn10 (Tetr)]* | (*92*) |

Φ and Ψ denote transcriptional and translational fusions, respectively

**Table S2. List of plasmids used in the study.**

| Plasmid | Relevant genotype | Source or reference |
|---|---|---|
| pAL27 | pKT25 $P_{lac}$-*cyaA*$_{T25}$-*phoQ*, $Kan^R$ | (*30*) |
| pAL33 | pUT18C $P_{lac}$-*cyaA*$_{T18}$-*mgrB*, $Amp^R$ | (*30*) |
| pAL38 | pEB52 $P_{trc}$-*gfpA206K-mgrB*, $Amp^R$ | (*30*) |
| pAL39 | pEB52 $P_{trc}$-*gfpA206K*, $Amp^R$ | (*30*) |
| pBAD24 | pMB1-derived plasmid, *araBAD,* $Amp^R$ | (*93*) |
| pCP20 | ori(pSC101) *rep101(ts) bla* λ$P_R$-FLP λcI(ts) cat. $Amp^R$, $Cam^R$ | (*73*) |
| pEB52 | pTrc99a with the NcoI site removed, $Amp^R$ | (*30*) |
| pJS1 | pEB52 $P_{trc}$-*yriA*, $Amp^R$ | This study |
| pJS2 | pEB52 $P_{trc}$-*yadW*, $Amp^R$ | This study |
| pJS3 | pEB52 $P_{trc}$-*yqhI*, $Amp^R$ | This study |
| pJS4 | pEB52 $P_{trc}$-*ykgS*, $Amp^R$ | This study |
| pJS5 | pEB52 $P_{trc}$-*yobF*, $Amp^R$ | This study |
| pJS6 | pEB52 $P_{trc}$-*yddY*, $Amp^R$ | This study |
| pJS7 | pEB52 $P_{trc}$-*mgtS*, $Amp^R$ | This study |
| pJS13 | pEB52 $P_{trc}$-*yoaI* (E31Q), $Amp^R$ | This study |
| pJS15 | pUT18C $P_{lac}$-*cyaA*$_{T18}$-*yoaI* (E31A), $Amp^R$ | This study |
| pJS16 | pUT18C $P_{lac}$-*cyaA*$_{T18}$-*yoaI* (E31Q), $Amp^R$ | This study |
| pJS18 | pEB52 $P_{trc}$-*yoaI* (E31A)-GGSG-6XHis, $Amp^R$ | This study |
| pJS24 | pEB52 $P_{trc}$-*yoaI* (E31Q)-GGSG-6XHis, $Amp^R$ | This study |
| pKD13 | ori(R6K) FRT-kan-FRT, $Amp^R$ | (*72*) |
| pKD46 | ori(pSC101) *rep101*(*ts*) $P_{araB}$-*gam-betexo araC*, $Amp^R$ | (*72*) |
| pKK14 | pKT25 $P_{lac}$-*cyaA*$_{T25}$-EnvZ, $Kan^R$ | (*29*) |
| pKT25 | *ori*$_{p15A}$ $P_{lac}$-*cyaA*$_{T25}$-MCS, $Kan^R$ | Euromedex |
| pMR120 | A derivative of pSMART VC (Lucigen) containing two copies of the *rrnB* transcription terminator, $P_{putative\ promoter}$-*yfpA206K*-$P_{tetA}$-*cfp*, $Cam^R$ | (*38*) |
| pPJ1 | pEB52 $P_{trc}$-*gfpA206K-mgtT*, $Amp^R$ | This study |
| pPJ2 | pEB52 $P_{trc}$-*gfpA206K-pmrR*, $Amp^R$ | This study |
| pPJ3 | pEB52 $P_{trc}$-*gfpA206K-ydgU*, $Amp^R$ | This study |
| pPJ4 | pEB52 $P_{trc}$-*yoaI-gfpA206K*, $Amp^R$ | This study |

| | | |
|---|---|---|
| pPJ5 | pEB52 $P_{trc}$-*gfpA206K-yqhI*, $Amp^R$ | This study |
| pPJ6 | pEB52 $P_{trc}$-*gfpA206K-yobF*, $Amp^R$ | This study |
| pPJ7 | pEB52 $P_{trc}$-*gfpA206K-yriA*, $Amp^R$ | This study |
| pPJ8 | pEB52 $P_{trc}$-*gfpA206K-yddY*, $Amp^R$ | This study |
| pPJ9 | pEB52 $P_{trc}$-*gfpA206K-yadW*, $Amp^R$ | This study |
| pPJ10 | pEB52 $P_{trc}$-*gfpA206K-yadX*, $Amp^R$ | This study |
| pPJ11 | pEB52 $P_{trc}$-*gfpA206K-yriB*, $Amp^R$ | This study |
| pPJ12 | pEB52 $P_{trc}$-*gfpA206K-ymiA*, $Amp^R$ | This study |
| pPJ13 | pEB52 $P_{trc}$-*gfpA206K-ymiC*, $Amp^R$ | This study |
| pPJ14 | pMR120 putative $P_{pmrR}$-*yfp*, $Cam^R$ | This study |
| pPJ16 | pMR120 putative $P_{asr-ydgU}$-*yfp*, $Cam^R$ | This study |
| pPJ17 | pMR120 putative $P_{ydgU}$-*yfp*, $Cam^R$ | This study |
| pPJ18 | pMR120 putative $P_{yoaI}$-*yfp*, $Cam^R$ | This study |
| pPJ19 | pMR120 putative $P_{yqhI}$-*yfp*, $Cam^R$ | This study |
| pPJ20 | pMR120 putative $P_{yriA-yriB}$-*yfp*, $Cam^R$ | This study |
| pPJ23 | pMR120 putative $P_{yadX-clcA-yadW}$-*yfp,* $Cam^R$ | This study |
| pSMART | pSMART VC *BamHI* (BAC) with single-copy replication origin *ori2 repE IncC parABC*, *oriV cat* $Cam^R$ | Lucigen Corporation |
| pSV14 | pEB52 $P_{trc}$-*yobF*-GGSG-6XHis, $Amp^R$ | This study |
| pSV16 | pMR120 putative $P_{ykgS}$-*yfp*, $Cam^R$ | This study |
| pSV17 | pMR120 putative $P_{proP-pmrR}$-*yfp*, $Cam^R$ | This study |
| pSV18 | pMR120 putative $P_{yadW}$-*yfp*, $Cam^R$ | This study |
| pSV19 | pMR120 putative $P_{ymiC}$-*yfp*, $Cam^R$ | This study |
| pSV20 | pMR120 putative $P_{ymiA-yciX-ymiC}$-*yfp*, $Cam^R$ | This study |
| pSV21 | pMR120 putative $P_{dinQ}$-*yfp*, $Cam^R$ | This study |
| pSV24 | pMR120 putative $P_{yobF}$-*yfp*, $Cam^R$ | This study |
| pSV25 | pEB52 $P_{trc}$-*gfpA206K-ykgS*, $Amp^R$ | This study |
| pSV26 | pEB52 $P_{trc}$-*pmrR*-GGSG-6XHis, $Amp^R$ | This study |
| pSV28 | pMR120 putative $P_{yddY}$-*yfp*, $Cam^R$ | This study |
| pSV29 | pMR120 putative $P_{mgtS-mgtT}$-*yfp*, $Cam^R$ | This study |
| pSV30 | pEB52 $P_{trc}$-*mgtT*, $Amp^R$ | This study |
| pSV31 | pEB52 $P_{trc}$-*ydgU*, $Amp^R$ | This study |
| pSV32 | pEB52 $P_{trc}$-*yadX*, $Amp^R$ | This study |

| | | |
|---|---|---|
| pSV33 | pEB52 $P_{trc}$-*ymiC*, $Amp^R$ | This study |
| pSV34 | pEB52 $P_{trc}$-*yoaI*, $Amp^R$ | This study |
| pSV35 | pEB52 $P_{trc}$-*pmrR*, $Amp^R$ | This study |
| pSV36 | pEB52 $P_{trc}$-*ymiA*, $Amp^R$ | This study |
| pSV37 | pEB52 $P_{trc}$-*yriB*, $Amp^R$ | This study |
| pSV38 | pEB52 $P_{trc}$-*pmrR*$_{ATG->TAA}$ where the start codon ATG is substituted with a stop TAA, $Amp^R$ | This study |
| pSV39 | pEB52 $P_{trc}$-*yoaI*$_{ATG->TAA}$ where the start codon ATG is substituted with a stop TAA, $Amp^R$ | This study |
| pSV40 | pEB52 $P_{trc}$-*mgtS*$_{ATG->TAA}$ where the start codon ATG is substituted with a stop TAA, $Amp^R$ | This study |
| pSV41 | pEB52 $P_{trc}$-*ykgS*$_{ATG->TAA}$ where the start codon ATG is substituted with a stop TAA, $Amp^R$ | This study |
| pSV42 | pEB52 $P_{trc}$-*ymiA*$_{ATG->TAA}$ where the start codon ATG is substituted with a stop TAA, $Amp^R$ | This study |
| pSV44 | pEB52 $P_{trc}$-*yoaI*-GGSG-6XHis, $Amp^R$ | This study |
| pSV45 | pEB52 $P_{trc}$-*yobF*$_{ATG->TAA}$ where the start codon ATG is substituted with a stop TAA, $Amp^R$ | This study |
| pSV47 | pUT18C $P_{lac}$-*cyaA*$_{T18}$-*yoaI*, $Amp^R$ | This study |
| pSV54 | pBAD24 $P_{araBAD}$-*dinQ,* $Amp^R$ | This study |
| pSV55 | pBAD24 $P_{araBAD}$-*dinQ*$_{ATG->TAA}$ where the start codon ATG is substituted with a stop TAA*,* $Amp^R$ | This study |
| pSV58 | pEB52 $P_{trc}$-*yriAB,* $Amp^R$ | This study |
| pSV59 | pEB52 $P_{trc}$-*yriB*$_{ATG->TAA}$ where the start codon ATG is substituted with a stop TAA, $Amp^R$ | This study |
| pSV60 | pEB52 $P_{trc}$-*mgrB,* $Amp^R$ | This study |
| pSV61 | pEB52 $P_{trc}$-*mgrB*$_{ATG->TAA}$ where the start codon ATG is substituted with a stop TAA, $Amp^R$ | This study |
| pSY68 | pKT25 $P_{lac}$-*cyaA*$_{T25}$-*phoR*, $Kan^R$ | (*29*) |
| pSV70 | pEB52 $P_{trc}$-*yoaI* (E31A). $Amp^R$ | This study |
| pTrc99a | *lacI*$^q$, $P_{trc}$-MCS, $Amp^R$ | (*94*) |
| pUT18C | pUC19 $P_{lac}$-*cyaA*$_{T18}$-MCS, $Amp^R$ | Euromedex |

$Amp^R$ = carries a carbenicillin/ampicillin-resistance marker; $Cam^R$ = carries a chloramphenicol-resistance marker; $Kan^R$ = carries a kanamycin-resistance marker

**Table S3. List of primers used in the study.**

| Primer | Sequence (5'-> 3') | Use |
|---|---|---|
| C-term-pmrR-F1 | AGCTTTCTTTATATCTGGTTTGCCACGTACGGCGGCAGCGGCCACCATCACCATCACCATTAAGCTTAATTAGCTGACCTACTAGAGTCG | Cloning of pSV26 |
| C-term-pmrR-R1 | GCTGACCACCAGCACGCTGAACACGGTAGTTAAACTTTCATAAACACGGTTTTTCATGCTTAATTTCTCCTCTTTAATTCTAGGTACCCG | Cloning of pSV26 |
| dinQ_del_screen_1 | aagatgggggcaaccaaaaaag | Validation of GSO195 |
| dinQ_del_screen_2 | ttcactctggcaatgcgcata | Validation of GSO195 |
| dinQ_F1 | ATATGAATTCCATCAATGCTAATACCATACTGAAACTATTGCA | Cloning of pSV21 |
| dinQ_R1 | ATATGGATCCCCGTTTTCTCCATGCGATGGAG | Cloning of pSV21 |
| dinQ_pBAD_R | CAGCGCAATTAACGCCCCTAGAACGATGATTGCTTTATCAATcatTGAATTCCTCCTGCTAGCCCAAA | Cloning of pSV54 |
| dinQ_pBAD_F | CTGGAACTGATCCGCTTTCTGCTTCAGCTTCTGAACTAAGGTACCCGGGGATCCTCTAG | Cloning of pSV54 |
| dinQ_stop_F | TAAATTGATAAAGCAATCATCGTTCTAGGGG | Cloning of pSV55 |
| dinQ_stop_R | TGAATTCCTCCTGCTAGCCCA | Cloning of pSV55 |
| envZ_F | TACTGAAGGCACTGGTCAGCC | Validation of JW3367 |
| envZ_R | AAGATTCTCGATCCGCGTAACACC | Validation of JW3367 |
| F_yoaI_BATCH | ACTGTCTAGAGatgAACGATCAAATGTTTGTCGAGACAC | Cloning of pSV47 |
| Fp_Inv_PCR_yobF | TAAGCTTAATTAGCTGACCTACTAGAGTCGAC | Cloning of pSV14 and pSV44 |
| JA1 | GCGTACGCAATCAAAATCCCCAGCCAATACAAcatTTAACACCATgcttaatttctcctctttaattctaggtacccg | Cloning of pSV58 |
| JA2 | AGACCGTAGGCCAGATAAGGTGTTTACGCTGATCAGGtaaGCTTAATTAGCTGACCTactagagtcgac | Cloning of pSV58 |
| JS4 | taaTTGTATTGGCTGGGGATTTTGATTGC | Cloning of pSV59 |
| lpxT_del_F | gacagccgtattgagctgatttcc | Validation of JW2162 |
| lpxT_del_R | aaagtgatacagaaagttaataagcgggg | Validation of JW2162 |

| | | |
|---|---|---|
| mgrB_forward | atgaaaaagtttcgatgggtcgttct | Cloning of pSV60 |
| mgrB_stop3 | TAAaaaaagtttcgatgggtcgttctg | Cloning of pSV61 |
| MgtS_del_F | AAAATTAAGGTAAGCGAGGAAACACACCACACCATAAACGGAGGCAAATAattccggggatccgtcgacctgcagttcgaagttcctatt | Construction of SV64 |
| MgtS_del_R | AACTGTAACAAGGGGCCGGTTAGGTGAGGGATTATCTCCGTTcatTAGTCtgtaggctggagctgcttcgaagttcctatact | Construction of SV64 |
| mgtS_del_screen_1 | cccgcgctttgttgatttaagtc | Validation of SV64 together with primer mgtT_del_screen2 |
| mgtS_forward_2_correct | TCTGGTTTTCTGGCCGCGTATTTCAGCCACAAATGGGATGACtaaTAAGCTTAATTAGCTGACCTactagagtcgac | Cloning of pJS7 |
| mgtS_reverse | AAATAAAATTATTCCCAGTACGGCCATAAAAACATTCATATTACCCAGcatgcttaatttctcctctttaattctaggtacccg | Cloning of pJS7 |
| mgtS_stop | TAACTGGGTAATTAAAATGTTTTTATGGCCGTACTGGGAATAATTTTATTTTCTGG | Cloning of pSV40 |
| mgtT_del_F | TATTTTCTGGTTTTCTGGCCGCGTATTTCAGCCACAAATGGGATGACTAAattccggggatccgtcgacc | Construction of SV31 |
| mgtT_del_R | CAAATCTATCCATGCAAGCATtcaCCGCCGGTTTACTGGCGGTTTTTTTTtgtaggctggagctgcttcg | Construction of SV31 |
| mgtT_del_screen1 | CACACCACACCATAAACGGAGG | Validation of SV31 |
| mgtT_del_screen2 | ACCGGAAGAAATCGCTGCATG | Validation of SV31 |
| mgtT_F | GAGAATTCTCAGCTGCGGTATTTACTGTCGG | Cloning of pSV29 |
| mgtT_R | GGGGATCCTAGTCATCCCATTTGTGGCTGAAATACG | Cloning of pPJ1 |
| mgtT_R1 | ATATGGATCCTATTTGCCTCCGTTTATGGTGTGGTG | Cloning of putative pSV29 |
| mgtT-forward | ATGAACGGAGATAATCCCTCACCTAAC | Cloning of pSV30 |
| N_term_pmrR_forward | ATGAAAAACCGTGTTTATGAAAGTTTAACTACCG | Cloning of pSV35 |
| pEB52-rev | GCTTAATTTCTCCTCTTTAATTCTAGGTACCCG | When targeting N-terminal-tagged small proteins, this primer, along with a small protein-specific forward primer, |

| | | |
|---|---|---|
| | | removes the tags by binding to the pEB52 backbone |
| phoB_del_F | cgaaaaagcatgggcgcgatta | Validation of JW0389 |
| phoB_del_R | agggcaggtaaccaaaaaatgcac | Validation of JW0389 |
| pmrR Primer 1 | CTGGTGGTCAGCAGCTTTCTTTATATCTGGTTTGCC ACGTACTGATAAGCTTAATTAGCTGACCTactagagt | Cloning of pPJ2 |
| pmrR Primer 2 | CACGCTGAACACGGTAGTTAAACTTTCATAAACACG GTTTTTCATtttgtatagttcatccatgccatgtg | Cloning of pPJ2 |
| pmrR_del_F | CTTAATCTCTGACGCGCATACTCTCCTCCAGGTTAA CGGAGGAGAGTGCAattccggggatccgtcgacc | Construction of SV35 |
| pmrR_del_R | GCCTGGGTACGGCTGAAGAAAGAtcaGTACGTGGCA AACCAGATATAAGtgtaggctggagctgcttcg | Construction of SV35 |
| pmrR_del_scree n1 | GCGAATTGATGAATAAGCTGAAACGG | Validation of SV35 |
| pmrR_del_scree n2 | CGTTTGTACGTATGGACAGCCG | Validation of SV35 |
| pmrR_stop | TAaAAAAACCGTGTTTATGAAAGTTTAACTACCG | Cloning of pSV38 |
| pmrR1_F | GAGAATTCTGGCTTCTACCTTGCCAGCG | Cloning of pPJ14 |
| pmrR1_R | GGGGATCCTGCACTCTCCTCCGTTAACCTG | Cloning of pPJ14 |
| pmrR2_F | GAGAATTCCCGACCACGCGTCACTATTACC | Cloning of pSV17 |
| pmrR2_R | GGGGATCCAGCTTTCCTCGCAGAGTTGG | Cloning of pSV17 |
| pSMART SV_19 | GACCATAACCGAAAGTAGTGAC | Sequencing primer for pMR120 based vectors |
| RP_Inv_yoaI | ATGGTGATGGTGATGGTGGCCGCTGCCGCCGCCG GTCCGTTCGATAAGAAG | Cloning of pSV44 |
| R_yoaI_BATCH | TACTGAATTCctaGCCGGTCCGTTCGATAAGAAG | Cloning of pSV47 |
| Rp_Inv_PCR_y obF | ATGGTGATGGTGATGGTGGCCGCTGCCGCCGATTG TTTTCTTCGCCCGCAGG | Cloning of pSV14 |
| SA26R | CATCCGCCAAAACAGCCAAG | Sequencing primer for pEB52 based vectors |
| yadW Primer 1 | TGCCGATGTCGTTTGGAGCAAAATATGAGTGATAAG CTTAATTAGCTGACCTactagagt | Cloning of pPJ9 |

| | | |
|---|---|---|
| yadW Primer 2 | ATTGGGCAAATTCTAACCCAATAATAATCGCCATtttgtatagttcatccatgccatgtg | Cloning of pPJ9 |
| yadW_del_F | GCCGCATCAGCCAGCGAGAATACTTGAACGAAATACCAGGGTATTAGATAattccggggatccgtcgacc | Construction of SV27 |
| yadW_del_R | TCGGTAAACTCCAGCGGCAGTGCTACGTCAtcaCTCATATTTTGCTCCAAACGtgtaggctggagctgcttcg | Construction of SV27 |
| yadW_del_screen1 | CGGGAAACCGCTATACTCGG | Validation of SV27 |
| yadW_del_screen2 | GGTGAAACCATACTGGAAGCCG | Validation of SV27 |
| yadW_forward | atgGCGATTATTATTGGGTTAGAATTTGCC | Cloning of pJS2 |
| yadW1_F | GAGAATTCTTCCATTTCGGTTGGTGCACCAA | Cloning of pSV18 |
| yadW1_R | GGGGATCCCCAATGATCACTTATTGGTCATACAAATAAGATGAC | Cloning of pSV18 |
| yadW2_F | GAGAATTCACCATGCGCACCATATCCATGG | Cloning of pPJ23 |
| yadW2_R | GGGGATCCGAAAATCCTTTGCAAAGCGTAATGTTTCAAAT | Cloning of pPJ23 |
| yadX Primer 1 | TCATCTTATTTGTATGACCAATAATAAGCTTAATTAGCTGACCTactagagt | Cloning of pPJ10 |
| yadX Primer 2 | CGGCATTTTACTCGCACGTTCCATtttgtatagttcatccatgccatgtg | Cloning of pPJ10 |
| yadX_del_F | AGTATAAAAGTTTTGTGCATTTGAAACATTACGCTTTGCAAAGGATTTTCattccggggatccgtcgacc | Construction of SV39 |
| yadX_del_R | AGAGGGAGTATCAGTTTTCATCCAATGATCACttaTTGGTCATACAAATAtgtaggctggagctgcttcg | Construction of SV39 |
| yadX_del_screen1 | CATGGAGGGTTCCTGATTCGTAG | Validation of SV39 |
| yadX_del_screen2 | GCCATAAACAAAATGGCTAACGGG | Validation of SV39 |
| yadX-forward | ATGGAACGTGCGAGTAAAATGC | Cloning of pSV32 |
| yoaI_A31Q | GCCGGTCCGctgGATAAGA | Cloning of pJS13 |
| yoaI_BATCH_E31A_F | GTCCGcgcGATAAGAAGAACG | Cloning of pJS15 |
| yoaI_BATCH_E31Q_F | GTCCGctgGATAAGAAGAACGGA | Cloning of pJS16 |
| yoaI_BATCH_E31_R | CGGCtagGAATTCATCGATATAActaagtaatatgg | Cloning of pJS15 and pJS16 |
| yoaI_E31A | GCCGGTCCGcgcGATAAGA | Cloning of pSV70 |

| | | |
|---|---|---|
| yoaI-forward | TAAGCTTAATTAGCTGACCTACTAGAGTCGAC | Cloning of pJS13 and pSV70 |
| yoaI_His_E31A | GCCGGTCCGcgcGATAAGA | Cloning of pJS18 |
| yoaI_His_E31Q_F2 | AACGGACAAAACCAGTACAACAGC | Cloning of pJS24 |
| yoaI_His_E31Q_R2 | CTTCTTATCcagCGGACCGG | Cloning of pJS24 |
| yoaI_His_R | GGCGGCAGCGGCCACCAT | Cloning of pJS18 |
| yddY Primer 1 | GCACGTTGCGTTTCATTTTGATAAGCTTAATTAGCTGACCTactagagt | Cloning of pPJ8 |
| yddY Primer 2 | GAGATCGACTAACTGCACCATtttgtatagttcatccatgccatgtg | Cloning of pPJ8 |
| yddY_del_F | GTCCTAAATCGCTTATTTCTTTTCAGTATATCTTCATATTTCAGGAGAATattccggggatccgtcgacc | Construction of SV29 |
| yddY_del_R | AATACATCTCCATAATTCACACCCTTATAAGGCTGGGAAATCAGACGGAAtgtaggctggagctgcttcg | Construction of SV29 |
| yddY_del_screen1 | CATACGATATTCGCTGTCACCG | Validation of SV29 |
| yddY_del_screen2 | CCGCGGCTTTAGCACGAAT | Validation of SV29 |
| yddY_F | GAGAATTCCTGTATATCGGTATCGCCTTCTATAAAGTGG | Cloning of pSV28 |
| yddY_forward | ATGGTGCAGTTAGTCGATCTCG | Cloning of pJS6 |
| yddY_R | GGGGATCCATTCTCCTGAAATATGAAGATATACTGAAAAG | Cloning of pSV28 |
| ydgU Primer 1 | ATTTTATGCGCACTGATTACCGCCCGTTTTTATCTTTCCTGATAAGCTTAATTAGCTGACCTactagagt | Cloning of pPJ3 |
| ydgU Primer 2 | AAGGATGATCAGAATGAACTCAAAGCGATAACGGCCCACCATtttgtatagttcatccatgccatgtg | Cloning of pPJ3 |
| ydgU_del_screen_1 | gcagcgaagaaacacgccaa | Validation of GSO219 |
| ydgU_del_screen_2 | caaaagcagacgtcaaactaattgca | Validation of GSO219 |
| ydgU-forward | ATGGTGGGCCGTTATCGCTT | Cloning of pSV31 |
| ydgU1_F | ACTCGAGAATTCGAAACCAACCACTCACGGAAGTCTG | Cloning of pPJ17 |
| ydgU1_R | CCGGGGATCCACTACCCTCCGCTAAAGGCG | Cloning of pPJ17 |
| ydgU2_F | ACTCGAGAATTCATACCCGTCCGGACTTATTGCC | Cloning of pPJ16 |

| | | |
|---|---|---|
| ydgU2_R | GGGGATCCTGTCATACCCTCAATTTGTTTTTTCATTTAACCC | Cloning of pPJ16 |
| ykgS_del_F2 | CAGAAGCAGAAAGACATTGGATCGAATTCTACAACCAGGTCGAGTCAGAaattccggggatccgtcgacctgcagttcgaagttcctatt | Construction of SV43 |
| ykgS_del_R2 | CAAAGTCATCGGGCATTATCTGAACATAAAACACTATCAATAAGTTGGAGtgtaggctggagctgcttcgaagttcctatact | Construction of SV43 |
| ykgS_del_screen1 | GGCAAACCCGTCCGTGTG | Validation of SV43 |
| ykgS_del_screen2 | AAGTCGCTGTCGTTCTCAAA | Validation of SV43 |
| ykgS_F | ATATCTCGAGAAGTGTTTTTTGTATAAATCGGACATTTTATCCTCG | Cloning of pSV25 |
| ykgS_F1 | ATATCTCGAGAAGTGTTTTTTGTATAAATCGGACATTTTATCCTCG | Cloning of pSV16 |
| ykgS_forward | ATGAGAATGATTGGCCTTCTTTATGATTTTAAGG | Cloning of pJS4 |
| ykgS_R | ATATGGATCCTTCTGACTCGACCTGGTTGTAGAATTCG | Cloning of pSV25 |
| ykgS_R1 | ATATGGATCCTTCTGACTCGACCTGGTTGTAGAATTCG | Cloning of pSV16 |
| ykgS_stop | TAAAGATAAATTGGCCTTCTTTATGATTTTAAGGATTATGCT | Cloning of pSV41 |
| ymiA Primer 1 | TGGCGGTTTTTCTTGGTTCTGCACTTTTCTGGGTGGTTGTCGCACTGCTGATTTGGAAAGTGTGGGGATAATAAGCTTAATTAGCTGACCTactagagt | Cloning of pPJ12 |
| ymiA Primer 2 | GCCAGGCTTTACGTTTTAATTCAGGATCGCGGCGGGGTTCCTGATTTCCAGAAGGCATTGCTAACCTCATtttgtatagttcatccatgccatgtg | Cloning of pPJ12 |
| ymiA_del_screen_1 | gattgaattattcactggagacgattcg | Validation of GSO225 |
| ymiA_del_screen_2 | tcatcttcagcaaacagttctataaaggc | Validation of GSO225 |
| ymiA_F1 | ATATGAATTCTCAGCCACAGTACAACCAAAATTGG | Cloning of pSV20 |
| ymiA_R1 | ATATGATCCATAATCGCCATCACTTATCAGCAAGAC | Cloning of pSV20 |
| ymiA_stop | TAAAGGTTAGCATAACCTTCTGGAAATCAGGAACCCC | Cloning of pSV42 |
| ymiA-forward | ATGAGGTTAGCAATGCCTTCTGGA | Cloning of pSV36 |

| | | |
|---|---|---|
| ymiC Primer 1 | CTGTCGATGCTCTTCTGGGCCGAACTCCTCTGGATCATTACTCACTGATAAGCTTAATTAGCTGACCTactagagt | Cloning of pPJ13 |
| ymiC Primer 2 | AGAAAACGCGCCCATCCAGGACCAATATTTCATATTTGTGTTGATCATtttgtatagttcatccatgccatgtg | Cloning of pPJ13 |
| ymiC_del_F | ACAGCTTGAGTTATCTCAACACAAAATAATAACCGTTAAGGGTGTAGCCTattccggggatccgtcgacc | Construction of SV41 |
| ymiC_del_R | GAAACCTTGTGGCAAAGCAAATGACAACCCCGCCGCAGCGGGGTCAAGGAtgtaggctggagctgcttcg | Construction of SV41 |
| ymiC_del_screen1 | GATGATCAGCCGAAACAATAATTATCATCATTC | Validation of SV41 |
| ymiC_del_screen2 | AAGCCCTGATTCTCTTCAGGGT | Validation of SV41 |
| ymiC_F1 | ATATGAATTCCAGGAGCAACTGGAGTCGTC | Cloning of pSV19 |
| ymiC_R1 | ATATGGATCCAGGCTACACCCTTAACGGTTATTATTTTG | Cloning of pSV19 |
| ymiC-forward | ATGATCAACACAAATATGAAATATTGGTCCTGGA | Cloning of pSV33 |
| yoaI Primer 1 | ATTGCTGTTGTACTGGTTTTGTCCGTTCTTCTTATCGAACGGACCGGCcgtaaaggagaagaacttttcactgg | Cloning pPJ4 |
| yoaI Primer 2 | GGCAAAAAACGATGACGTGATAATCAGTGTCTCGACAAACATTTGATCGTTcatgcttaatttctcctctttaattctaggtac | Cloning pPJ4 |
| yoaI_del_screen_1 | tggttgccgaccctctactc | Validation of GSO232 |
| yoaI_del_screen_2 | agcctgcttactttacgcttgc | Validation of GSO232 |
| yoaI_F | GAGAATTCGCGGCTGGCTGCCCATTAA | Cloning of pPJ18 |
| yoaI_R | GGGGATCCAGCATGCCCCCGGGAGATA | Cloning pPJ18 |
| yoaI_stop | taAAACGATCAATAATTTGTCGAGACACTGATTATCACGTCATCG | Cloning of pSV39 |
| yoaI-forward | TAAGCTTAATTAGCTGACCTACTAGAGTCGAC | Cloning of pSV34 |
| yoaI-rev | GCCGGTCCGTTCGATAAGAAGA | Cloning of pSV34 |
| yobF Primer 1 | CCTTATATTGGTGCCTCATGCAGTAATGTGTCAGTTTTATCTATGTTATGCCTGCGGGCGAAGAAAACAATCcgtaaaggagaagaacttttcactgg | Cloning of pPJ6 |
| yobF Primer 2 | CTCGGCAGAGAAGCGGTATTCAACGTCAACGTGTTTACTCAGGACTTCTTTACTGAAAATGCCACAcatgcttaatttctcctctttaattctaggtac | Cloning of pPJ6 |

| | | |
|---|---|---|
| yobF_del_F | AAAGAAGTCCTGAGTAAACACGTTGACGTTGAATACCGCTTCTCTGCCGAattccggggatccgtcgacc | Construction of SV33 |
| yobF_del_R | TTGAACCACTTAACCTGACCTTTAATCTTTGCCATTTGAAAAATTCCttatgtaggctggagctgcttcg | Construction of SV33 |
| yobF_del_screen1 | GTGTGCAAAAAAAGTGGAAGACGT | Validation of SV33 |
| yobF_del_screen2 | CCATTACCCTGGATAGCGGAG | Validation of SV33 |
| yobF_F1 | ATATGAATTCTAATTTATTAGGATGTTTACATCGGATTTGTGATTAAGCG | Cloning of pSV24 |
| yobF_R1 | ATATGGATCCAAACAGAACTGTACCTCGTTTAACCC | Cloning of pSV24 |
| yobF_reverse | GATTGTTTTCTTCGCCCGCAGG | Cloning of pJS5 |
| yobF_stop | taaTGTGGCATTTTCAGTAAAGAAGTCCTG | Cloning of pSV45 |
| yqhI Primer 1 | ATCTCGAAACGTGAGTTCCTCAACCTTGCGGCGAAGTGCGGTAGGCGGGATGACGGCATTAGCGTTGTTTGATAAGCTTAATTAGCTGACCTactagagt | Cloning of pPJ5 |
| yqhI Primer 2 | TTTCCCGTGAGCGTAATAGTCATAGTAATCCAGCAACTCTTGTGGGAAATCTTTGGCGGTTAAACGCGGCATttgtatagttcatccatgccatgtg | Cloning of pPJ5 |
| yqhI_del_F2 | AATACGTGCTTATGCTTTGCTTAAAAAAACACCAACTGAGGAGTGCAACGaattccggggatccgtcgacctgcagttcgaagttcctatt | Construction of SV37 |
| yqhI_del_R2 | GGTCGGTAAACTCTACCTGAGTCGCCAGCGCATAATTTGGCTTGAGCAAAtgtaggctggagctgcttcgaagttcctatact | Construction of SV37 |
| yqhI_del_screen1 | TGTTTCTAGTTTAGCGATTCGCCAG | Validation of SV37 |
| yqhI_del_screen2 | GGGCTTCACCAGATAACCCC | Validation of SV37 |
| yqhI_F | GAGAATTCTATTTACGTGAACCGCCAGAGCC | Cloning of pPJ19 |
| yqhI_forward | ATGCCGCGTTTAACCGCC | Cloning of pJS3 |
| yqhI_R | GGGGATCCCGTTGCACTCCTCAGTTGGTG | Cloning of pPJ19 |
| yriA Primer 1 | GCAGACCGTAGGCCAGATAAGGTGTTTACGCTGATCAGGTAATAAGCTTAATTAGCTGACCTactagagt | Cloning of pPJ7 |
| yriA Primer 2 | GTACGCAATCAAAATCCCCAGCCAATACAACATtttgtatagttcatccatgccatgtg | Cloning of pPJ7 |
| yriA_F | TCGAGAATTCAAATCAGGTCAGGCTTAAGTAGCGAC | Cloning of pPJ20 |
| yriA_forward | ATGTTGTATTGGCTGGGGATTTTGAT | Cloning of pJS1 |

| | | |
|---|---|---|
| yriA_R | GGGGATCCTTAACACCATCATATTTTCCATCATTAG TGTGATC | Cloning of pPJ20 |
| yriAB_del_F | ACACTATTACAACAGAAAATAACCAGATGATCACAC TAATGATGGAAAATattccggggatccgtcgacc | Construction of SV25 |
| yriAB_del_R | TGTTCCTGAACGCCCGCATATGCGGGCGTTTTGCT TTTTGGCGCGCCTTGgtgtaggctggagctgcttc | Construction of SV25 |
| yriAB_del_screen1 | GAACGCGTAAGGGATAACGC | Validation of SV25 |
| yriAB_del_screen2 | CTGATCAACACCGTTCACGTTG | Validation of SV25 |
| yriB Primer 1 | GATTTTGATTGCGTACGCAGACCGtagTAAGCTTAAT TAGCTGACCTactagagt | Cloning of pPJ11 |
| yriB primer 2 | CCCAGCCAATACAACATTTAACACcattttgtatagttcatcca tgccatgtg | Cloning of pPJ11 |
| yriB-forward | atgGTGTTAAATGTTGTATTGGCTGGG | Cloning of pSV37 |

**Table S4. List of all proteins with ≥3-fold increase in read counts from Ribo-RET and differentially expressed transcripts from RNA-Seq.**

| Ribo-RET | | | | | RNA-Seq Differential Expression | | | | | |
|---|---|---|---|---|---|---|---|---|---|---|
| gene | size | no_stress | low_mg | fold_change | gene | base_Mean | log2FoldChange | lfcSE | pvalue | padj |
| *phnC* | standard | 0.68 | 79.66 | 117.25 | *mgtS* | 14972.46 | 7.82 | 0.31 | 2.11E-140 | 6.60E-137 |
| *iraM* | standard | 3.4 | 247.31 | 72.8 | *ymgL* | 17.66 | 7.53 | 2.75 | 2.84E-06 | 2.56E-05 |
| *phoR* | standard | 0.68 | 44.65 | 65.71 | *mgtA* | 3789.38 | 7.13 | 0.39 | 9.56E-75 | 9.97E-72 |
| *ptrA* | standard | 1.36 | 72.66 | 53.47 | *ymgK* | 236.85 | 6.93 | 0.62 | 6.26E-29 | 5.02E-27 |
| *phoE* | standard | 5.44 | 242.93 | 44.69 | *iraM* | 205.15 | 6.71 | 0.63 | 4.09E-27 | 2.78E-25 |
| *eptA* | standard | 5.44 | 232.43 | 42.76 | *mgtL* | 11342.3 | 6.61 | 0.43 | 1.18E-54 | 5.27E-52 |
| *ydeT* | standard | 4.76 | 148.38 | 31.2 | *ydeT* | 28.94 | 6.33 | 1.43 | 6.39E-06 | 5.33E-05 |
| *arnC* | standard | 0.68 | 20.13 | 29.64 | *ydeS* | 9.66 | 6.26 | 2.76 | 1.15E-04 | 7.48E-04 |
| *yciX* | standard | 1.36 | 38.96 | 28.67 | *asr* | 332.57 | 5.91 | 0.48 | 3.80E-35 | 4.96E-33 |
| *waaH* | standard | 2.04 | 58.22 | 28.56 | *ybjG* | 1315.13 | 5.78 | 0.35 | 4.37E-63 | 3.42E-60 |
| *rstB* | standard | 0.68 | 18.82 | 27.7 | *mgrB* | 6552.83 | 5.68 | 0.31 | 4.00E-76 | 6.25E-73 |
| *phoQ* | standard | 2.04 | 54.71 | 26.84 | *phnD* | 25.86 | 5.34 | 1.17 | 1.71E-06 | 1.61E-05 |
| *ugpB* | standard | 10.87 | 288.45 | 26.53 | *yebO* | 1871.52 | 5.1 | 0.33 | 1.23E-55 | 6.42E-53 |
| *phoB* | standard | 29.21 | 711.72 | 24.36 | *yriB* | 18.19 | 4.72 | 1.24 | 3.12E-05 | 2.30E-04 |
| *appC* | standard | 6.79 | 162.39 | 23.9 | *phnC* | 17.35 | 4.63 | 1.25 | 4.47E-05 | 3.18E-04 |
| *otsB* | standard | 0.68 | 14.88 | 21.9 | *pstS* | 274.79 | 4.41 | 0.41 | 1.93E-28 | 1.48E-26 |
| *argD* | standard | 10.87 | 199.6 | 18.36 | *yfeO* | 91.2 | 4.31 | 0.56 | 1.21E-15 | 2.81E-14 |
| *cysD* | standard | 20.38 | 358.92 | 17.61 | *ydgU* | 24.42 | 4.31 | 0.99 | 2.00E-06 | 1.85E-05 |
| *cbl* | standard | 2.04 | 35.02 | 17.18 | *dgcZ* | 61.86 | 4.29 | 0.64 | 3.40E-12 | 6.33E-11 |
| *ynaL* | standard | 1.36 | 21.01 | 15.46 | *pgpC* | 397.23 | 4.24 | 0.37 | 2.38E-31 | 2.26E-29 |
| *asr* | standard | 51.64 | 796.2 | 15.42 | *ygdR* | 155.39 | 4.19 | 0.46 | 9.92E-21 | 3.34E-19 |
| *yriA* | small | 2.72 | 40.71 | 14.98 | *alaV* | 973.94 | 4.18 | 0.36 | 2.73E-32 | 2.85E-30 |
| *glsA* | standard | 0.68 | 10.07 | 14.82 | *alaU* | 973.94 | 4.18 | 0.36 | 2.73E-32 | 2.85E-30 |
| *yciW* | standard | 8.15 | 119.06 | 14.6 | *alaT* | 973.94 | 4.18 | 0.36 | 2.73E-32 | 2.85E-30 |
| *yohC* | standard | 8.15 | 118.18 | 14.5 | *metZ* | 673.96 | 4.12 | 0.43 | 6.41E-23 | 2.87E-21 |
| *yeeE* | standard | 4.08 | 58.65 | 14.39 | *metW* | 673.96 | 4.12 | 0.43 | 6.41E-23 | 2.87E-21 |
| *sbp* | standard | 6.79 | 96.3 | 14.17 | *metV* | 673.96 | 4.12 | 0.43 | 6.41E-23 | 2.87E-21 |
| *ydgU* | small | 2.04 | 27.14 | 13.31 | *glyW* | 9479.84 | 4.06 | 0.33 | 1.60E-35 | 2.27E-33 |
| *narU* | standard | 1.36 | 17.95 | 13.21 | *glyV* | 9479.84 | 4.06 | 0.33 | 1.60E-35 | 2.27E-33 |
| *arnB* | standard | 11.55 | 151.45 | 13.11 | *glyX* | 9479.84 | 4.06 | 0.33 | 1.60E-35 | 2.27E-33 |
| *argA* | standard | 16.31 | 212.29 | 13.02 | *glyY* | 9479.84 | 4.06 | 0.33 | 1.60E-35 | 2.27E-33 |
| *ais* | standard | 9.51 | 123 | 12.93 | *ais* | 16.7 | 4.03 | 1.15 | 6.60E-05 | 4.54E-04 |
| *ycaP* | standard | 1.36 | 17.51 | 12.88 | *arnB* | 38.15 | 3.91 | 0.76 | 2.56E-08 | 3.31E-07 |
| *phoA* | standard | 12.23 | 156.26 | 12.78 | *ugpB* | 80.2 | 3.83 | 0.57 | 1.18E-12 | 2.26E-11 |
| *ydeQ* | standard | 1.36 | 17.07 | 12.56 | *alaX* | 648.46 | 3.79 | 0.38 | 2.29E-24 | 1.21E-22 |
| *yncG* | standard | 2.04 | 25.39 | 12.46 | *alaW* | 648.46 | 3.79 | 0.38 | 2.29E-24 | 1.21E-22 |

| | | | | | | | | | | |
|---|---|---|---|---|---|---|---|---|---|---|
| *ilvB* | standard | 1.36 | 16.2 | 11.92 | *rttR* | 21.62 | 3.74 | 1.06 | 4.26E-05 | 3.06E-04 |
| *yoaI* | small | 13.59 | 161.95 | 11.92 | *metY* | 276.5 | 3.73 | 0.43 | 6.96E-19 | 2.05E-17 |
| *cysP* | standard | 63.87 | 758.56 | 11.88 | *leuW* | 4113.99 | 3.68 | 0.35 | 1.34E-27 | 9.30E-26 |
| *yfcG* | standard | 4.08 | 48.15 | 11.81 | *arnC* | 29.57 | 3.66 | 0.82 | 9.33E-07 | 9.24E-06 |
| *lysA* | standard | 2.04 | 23.64 | 11.6 | *rstA* | 323.01 | 3.65 | 0.38 | 8.71E-23 | 3.84E-21 |
| *artP* | standard | 43.48 | 491.11 | 11.29 | *msrQ* | 38.7 | 3.63 | 0.72 | 5.34E-08 | 6.58E-07 |
| *yhbW* | standard | 1.36 | 15.32 | 11.27 | *arnF* | 56.91 | 3.62 | 0.61 | 3.42E-10 | 5.22E-09 |
| *dacD* | standard | 0.68 | 7.44 | 10.95 | *phoA* | 78.32 | 3.62 | 0.55 | 3.42E-12 | 6.33E-11 |
| *ruvC* | standard | 0.68 | 7.44 | 10.95 | *ymcF* | 40.82 | 3.61 | 0.71 | 3.74E-08 | 4.73E-07 |
| *ydbL* | standard | 1.36 | 14.88 | 10.95 | *dacC* | 178.45 | 3.59 | 0.42 | 1.11E-18 | 3.21E-17 |
| *argG* | standard | 20.38 | 217.98 | 10.69 | *ileV* | 995.19 | 3.57 | 0.34 | 3.75E-26 | 2.35E-24 |
| *argC* | standard | 6.11 | 64.78 | 10.59 | *ileU* | 995.19 | 3.57 | 0.34 | 3.75E-26 | 2.35E-24 |
| *ybhP* | standard | 0.68 | 7 | 10.31 | *ileT* | 995.19 | 3.57 | 0.34 | 3.75E-26 | 2.35E-24 |
| *yobF* | small | 12.91 | 132.63 | 10.27 | *phoQ* | 258.31 | 3.57 | 0.38 | 1.64E-21 | 6.11E-20 |
| *pstS* | standard | 48.92 | 490.68 | 10.03 | *phoB* | 66.9 | 3.56 | 0.57 | 4.75E-11 | 7.87E-10 |
| *nadD* | standard | 1.36 | 13.57 | 9.99 | *eptA* | 15.82 | 3.5 | 1.11 | 1.72E-04 | 0.001078186 |
| *gadE* | standard | 2.72 | 26.7 | 9.82 | *arnA* | 35.6 | 3.49 | 0.74 | 2.45E-07 | 2.70E-06 |
| *ydeS* | standard | 2.72 | 26.7 | 9.82 | *ydhI* | 62.43 | 3.45 | 0.58 | 3.12E-10 | 4.78E-09 |
| *bcsB* | standard | 0.68 | 6.57 | 9.66 | *pdeR* | 34.61 | 3.44 | 0.74 | 3.72E-07 | 3.97E-06 |
| *pcm* | standard | 0.68 | 6.57 | 9.66 | *pstC* | 127.36 | 3.42 | 0.46 | 7.14E-15 | 1.56E-13 |
| *yadW* | small | 1.36 | 13.13 | 9.66 | *secG* | 2556.53 | 3.41 | 0.38 | 2.56E-20 | 8.08E-19 |
| *pagP* | standard | 8.15 | 75.29 | 9.23 | *phoP* | 729.83 | 3.38 | 0.33 | 5.44E-25 | 3.27E-23 |
| *pstA* | standard | 2.04 | 18.82 | 9.23 | *slyB* | 3744.69 | 3.34 | 0.3 | 7.35E-30 | 6.39E-28 |
| *waaQ* | standard | 2.04 | 18.82 | 9.23 | *hokD* | 178.33 | 3.31 | 0.42 | 3.78E-16 | 8.96E-15 |
| *aroE* | standard | 0.68 | 6.13 | 9.02 | *pagP* | 21.67 | 3.28 | 0.91 | 3.55E-05 | 2.57E-04 |
| *mgrB* | small | 250.71 | 2248.97 | 8.97 | *yliM* | 5741.91 | 3.21 | 0.32 | 5.03E-24 | 2.58E-22 |
| *dtpD* | standard | 2.72 | 24.07 | 8.86 | *yjcB* | 202.42 | 3.2 | 0.4 | 7.82E-17 | 1.93E-15 |
| *tatD* | standard | 8.15 | 71.78 | 8.8 | *relE* | 150.84 | 3.19 | 0.44 | 6.33E-14 | 1.28E-12 |
| *bcr* | standard | 2.04 | 17.51 | 8.59 | *rstB* | 99.93 | 3.16 | 0.49 | 8.30E-12 | 1.48E-10 |
| *pstC* | standard | 44.16 | 377.31 | 8.54 | *ynfB* | 711.59 | 3.09 | 0.34 | 1.32E-20 | 4.34E-19 |
| *blc* | standard | 2.72 | 23.2 | 8.54 | *ybfA* | 383.07 | 3.09 | 0.38 | 3.59E-17 | 9.13E-16 |
| *nudL* | standard | 2.72 | 23.2 | 8.54 | *deoR* | 205.33 | 3.08 | 0.4 | 6.56E-16 | 1.54E-14 |
| *insQ* | standard | 6.79 | 56.9 | 8.38 | *sraG* | 43.7 | 3.08 | 0.65 | 2.09E-07 | 2.33E-06 |
| *cysJ* | standard | 60.47 | 504.25 | 8.34 | *valT* | 8108.95 | 3.08 | 0.33 | 7.92E-22 | 3.02E-20 |
| *pdeR* | standard | 8.15 | 67.41 | 8.27 | *valZ* | 8108.95 | 3.08 | 0.33 | 7.92E-22 | 3.02E-20 |
| *iaaA* | standard | 30.57 | 251.69 | 8.23 | *valU* | 8108.95 | 3.08 | 0.33 | 7.92E-22 | 3.02E-20 |
| *yhdN* | standard | 4.08 | 32.83 | 8.05 | *valX* | 8108.95 | 3.08 | 0.33 | 7.92E-22 | 3.02E-20 |
| *chaC* | standard | 5.44 | 43.33 | 7.97 | *valY* | 8108.95 | 3.08 | 0.33 | 7.92E-22 | 3.02E-20 |
| *mgtT* | small | 1082.31 | 8622.95 | 7.97 | *tqsA* | 29.99 | 3.05 | 0.76 | 6.71E-06 | 5.58E-05 |
| *gadX* | standard | 4.08 | 31.95 | 7.84 | *lysT* | 11718.26 | 3.05 | 0.33 | 3.00E-21 | 1.04E-19 |
| *rssB* | standard | 4.08 | 31.95 | 7.84 | *lysW* | 11718.26 | 3.05 | 0.33 | 3.00E-21 | 1.04E-19 |
| *hisG* | standard | 13.59 | 105.93 | 7.8 | *lysY* | 11718.26 | 3.05 | 0.33 | 3.00E-21 | 1.04E-19 |

| | | | | | | | | | | |
|---|---|---|---|---|---|---|---|---|---|---|
| *cysW* | standard | 0.68 | 5.25 | 7.73 | *lysZ* | 11718.26 | 3.05 | 0.33 | 3.00E-21 | 1.04E-19 |
| *glaR* | standard | 0.68 | 5.25 | 7.73 | *lysQ* | 11718.26 | 3.05 | 0.33 | 3.00E-21 | 1.04E-19 |
| *ibsD* | small | 0.68 | 5.25 | 7.73 | *lysV* | 11718.26 | 3.05 | 0.33 | 3.00E-21 | 1.04E-19 |
| *ibsE* | small | 0.68 | 5.25 | 7.73 | *appC* | 46.53 | 3.02 | 0.63 | 1.57E-07 | 1.80E-06 |
| *ybdR* | standard | 0.68 | 5.25 | 7.73 | *fabR* | 293.94 | 3.02 | 0.37 | 4.17E-17 | 1.05E-15 |
| *hisD* | standard | 9.51 | 73.1 | 7.68 | *csrC* | 1142.49 | 3 | 0.32 | 6.80E-22 | 2.80E-20 |
| *tqsA* | standard | 8.15 | 61.28 | 7.52 | *phoR* | 33.35 | 2.98 | 0.72 | 3.93E-06 | 3.47E-05 |
| *purF* | standard | 15.63 | 117.31 | 7.51 | *ygcN* | 30.28 | 2.95 | 0.75 | 1.01E-05 | 8.10E-05 |
| *glcD* | standard | 8.15 | 60.84 | 7.46 | *yfiS* | 34.41 | 2.94 | 0.74 | 7.12E-06 | 5.88E-05 |
| *speG* | standard | 26.5 | 196.53 | 7.42 | *queE* | 213.58 | 2.94 | 0.39 | 5.47E-15 | 1.21E-13 |
| *ftsQ* | standard | 1.36 | 10.07 | 7.41 | *yodB* | 20 | 2.9 | 0.92 | 1.76E-04 | 0.001100074 |
| *ygbE* | standard | 3.4 | 24.95 | 7.34 | *tadA* | 238.08 | 2.89 | 0.38 | 6.15E-15 | 1.36E-13 |
| *yqhI* | small | 2.04 | 14.88 | 7.3 | *arnD* | 13.79 | 2.89 | 1.12 | 0.001001481 | 0.005046111 |
| *miaA* | standard | 13.59 | 96.73 | 7.12 | *ybgS* | 71.99 | 2.87 | 0.54 | 1.44E-08 | 1.91E-07 |
| *speC* | standard | 19.7 | 140.07 | 7.11 | *metU* | 243.67 | 2.87 | 0.42 | 6.17E-13 | 1.21E-11 |
| *appB* | standard | 0.68 | 4.81 | 7.09 | *metT* | 243.67 | 2.87 | 0.42 | 6.17E-13 | 1.21E-11 |
| *argH* | standard | 2.72 | 19.26 | 7.09 | *serV* | 8051.92 | 2.86 | 0.35 | 2.19E-17 | 5.70E-16 |
| *artJ* | standard | 13.59 | 96.3 | 7.09 | *cspG* | 125.16 | 2.85 | 0.44 | 1.28E-11 | 2.26E-10 |
| *cobC* | standard | 2.72 | 19.26 | 7.09 | *ugpA* | 23.5 | 2.84 | 0.85 | 9.04E-05 | 6.07E-04 |
| *rcdA* | standard | 2.72 | 19.26 | 7.09 | *gltW* | 2671.27 | 2.83 | 0.34 | 1.73E-17 | 4.54E-16 |
| *ydgD* | standard | 4.76 | 33.7 | 7.09 | *gltU* | 2671.27 | 2.83 | 0.34 | 1.73E-17 | 4.54E-16 |
| *hrpB* | standard | 4.08 | 28.89 | 7.09 | *gltT* | 2671.27 | 2.83 | 0.34 | 1.73E-17 | 4.54E-16 |
| *ydgI* | standard | 29.21 | 204.85 | 7.01 | *gltV* | 2671.27 | 2.83 | 0.34 | 1.73E-17 | 4.54E-16 |
| *yodB* | standard | 5.44 | 37.64 | 6.93 | *speG* | 402.93 | 2.82 | 0.35 | 2.21E-16 | 5.39E-15 |
| *phoH* | standard | 16.99 | 116.87 | 6.88 | *glnW* | 5581.76 | 2.79 | 0.34 | 2.38E-17 | 6.10E-16 |
| *ydhJ* | standard | 23.78 | 163.27 | 6.87 | *glnU* | 5581.76 | 2.79 | 0.34 | 2.38E-17 | 6.10E-16 |
| *yebW* | standard | 7.47 | 50.77 | 6.79 | *phnF* | 9.4 | 2.78 | 1.4 | 0.004516358 | 0.018545518 |
| *dinQ* | small | 1.36 | 9.19 | 6.76 | *ydhJ* | 35.29 | 2.76 | 0.7 | 8.20E-06 | 6.70E-05 |
| *yfbP* | standard | 0.68 | 4.38 | 6.44 | *appB* | 29.37 | 2.76 | 0.75 | 2.63E-05 | 1.97E-04 |
| *insD2* | standard | 0.68 | 4.38 | 6.44 | *pstA* | 84.17 | 2.76 | 0.5 | 3.89E-09 | 5.37E-08 |
| *lhr* | standard | 0.68 | 4.38 | 6.44 | *phoH* | 157.53 | 2.74 | 0.55 | 6.19E-08 | 7.50E-07 |
| *nhoA* | standard | 0.68 | 4.38 | 6.44 | *glsA* | 31.04 | 2.74 | 0.74 | 2.48E-05 | 1.86E-04 |
| *recX* | standard | 0.68 | 4.38 | 6.44 | *ycgX* | 11.14 | 2.72 | 1.26 | 0.002969559 | 0.012959207 |
| *ugpA* | standard | 0.68 | 4.38 | 6.44 | *clcA* | 33.59 | 2.7 | 0.71 | 1.68E-05 | 1.29E-04 |
| *yacH* | standard | 0.68 | 4.38 | 6.44 | *relB* | 212.04 | 2.68 | 0.39 | 9.27E-13 | 1.79E-11 |
| *ycaC* | standard | 2.04 | 13.13 | 6.44 | *mlaF* | 310.63 | 2.64 | 0.36 | 2.15E-14 | 4.54E-13 |
| *yidD* | standard | 0.68 | 4.38 | 6.44 | *rhsC* | 20.37 | 2.59 | 0.88 | 3.93E-04 | 0.002230904 |
| *patA* | standard | 3.4 | 21.01 | 6.18 | *pmrD* | 10.57 | 2.59 | 1.28 | 0.004234691 | 0.017596745 |
| *yncJ* | standard | 50.28 | 310.34 | 6.17 | *ymdF* | 73.54 | 2.57 | 0.52 | 9.59E-08 | 1.12E-06 |
| *rseA* | standard | 205.86 | 1264.99 | 6.14 | *mlrA* | 23.28 | 2.55 | 0.82 | 2.35E-04 | 0.001436063 |
| *opgG* | standard | 13.59 | 83.17 | 6.12 | *mqsA* | 81.15 | 2.54 | 0.5 | 5.50E-08 | 6.75E-07 |
| *ybiO* | standard | 1.36 | 8.32 | 6.12 | *zntA* | 87.62 | 2.54 | 0.48 | 1.74E-08 | 2.26E-07 |

| | | | | | | | | | |
|---|---|---|---|---|---|---|---|---|---|
| *yfeO* | standard | 1.36 | 8.32 | 6.12 | *mdfA* | 45.72 | 2.54 | 0.62 | 5.74E-06 | 4.90E-05 |
| *hemD* | standard | 2.04 | 12.26 | 6.01 | *mqsR* | 84.83 | 2.51 | 0.49 | 4.53E-08 | 5.69E-07 |
| *iscX* | standard | 12.23 | 73.54 | 6.01 | *thrV* | 83.75 | 2.47 | 0.53 | 4.55E-07 | 4.79E-06 |
| *mdlA* | standard | 2.04 | 12.26 | 6.01 | *slp* | 50.9 | 2.47 | 0.6 | 6.20E-06 | 5.20E-05 |
| *argR* | standard | 27.18 | 163.27 | 6.01 | *ycfJ* | 63.77 | 2.46 | 0.55 | 1.07E-06 | 1.04E-05 |
| *pmrR* | small | 32.61 | 193.91 | 5.95 | *thrU* | 145.51 | 2.43 | 0.42 | 6.65E-10 | 9.68E-09 |
| *yddY* | small | 22.42 | 130.88 | 5.84 | *ytjA* | 131.58 | 2.42 | 0.44 | 4.71E-09 | 6.41E-08 |
| *yeeD* | standard | 108.03 | 627.68 | 5.81 | *folA* | 228.35 | 2.4 | 0.38 | 5.26E-11 | 8.65E-10 |
| *apaH* | standard | 2.72 | 15.76 | 5.8 | *mlaE* | 239.82 | 2.4 | 0.37 | 1.49E-11 | 2.60E-10 |
| *folM* | standard | 3.4 | 19.7 | 5.8 | *yddY* | 25.91 | 2.4 | 0.79 | 3.10E-04 | 0.001817096 |
| *ibsA* | small | 0.68 | 3.94 | 5.8 | *ycaC* | 70.21 | 2.39 | 0.53 | 9.42E-07 | 9.30E-06 |
| *insD3* | standard | 0.68 | 3.94 | 5.8 | *appA* | 25.61 | 2.37 | 0.77 | 2.88E-04 | 0.001708407 |
| *insD4* | standard | 0.68 | 3.94 | 5.8 | *yadE* | 24.1 | 2.37 | 0.8 | 3.94E-04 | 0.002230904 |
| *insD5* | standard | 0.68 | 3.94 | 5.8 | *ykfI* | 55.47 | 2.37 | 0.56 | 3.18E-06 | 2.85E-05 |
| *insD6* | standard | 0.68 | 3.94 | 5.8 | *arfA* | 24.15 | 2.36 | 0.8 | 4.15E-04 | 0.002324926 |
| *mhpF* | standard | 0.68 | 3.94 | 5.8 | *ychS* | 106.85 | 2.34 | 0.47 | 9.09E-08 | 1.08E-06 |
| *xylB* | standard | 0.68 | 3.94 | 5.8 | *yohC* | 26.27 | 2.32 | 0.77 | 3.43E-04 | 0.00200324 |
| *ydcK* | standard | 1.36 | 7.88 | 5.8 | *yjbJ* | 153.23 | 2.32 | 0.42 | 4.38E-09 | 5.98E-08 |
| *btuE* | standard | 19.7 | 113.81 | 5.78 | *paoD* | 10.75 | 2.3 | 1.23 | 0.00655796 | 0.025024217 |
| *yjdC* | standard | 8.83 | 48.59 | 5.5 | *yiaG* | 86.74 | 2.3 | 0.5 | 6.84E-07 | 6.95E-06 |
| *glsB* | standard | 10.19 | 56.03 | 5.5 | *rmf* | 1448.26 | 2.29 | 0.32 | 2.75E-13 | 5.45E-12 |
| *recJ* | standard | 6.79 | 37.21 | 5.48 | *ycaP* | 13.38 | 2.27 | 1.07 | 0.004022703 | 0.01682759 |
| *dosC* | standard | 16.31 | 87.54 | 5.37 | *yjdN* | 14.7 | 2.25 | 1.02 | 0.003375377 | 0.01444809 |
| *yecE* | standard | 2.04 | 10.94 | 5.37 | *symR* | 13.43 | 2.24 | 1.08 | 0.004360634 | 0.01804818 |
| *cpxP* | standard | 86.29 | 461.35 | 5.35 | *hemL* | 436.05 | 2.23 | 0.35 | 1.79E-11 | 3.08E-10 |
| *ampH* | standard | 38.05 | 202.22 | 5.32 | *glk* | 152.05 | 2.22 | 0.42 | 1.62E-08 | 2.12E-07 |
| *ycgV* | standard | 13.59 | 71.78 | 5.28 | *arrS* | 71.94 | 2.22 | 0.54 | 5.76E-06 | 4.90E-05 |
| *ymgE* | standard | 19.02 | 100.24 | 5.27 | *ybjX* | 352.08 | 2.22 | 0.38 | 9.80E-10 | 1.41E-08 |
| *bepA* | standard | 20.38 | 105.49 | 5.18 | *yagP* | 15.62 | 2.21 | 0.99 | 0.003308268 | 0.014180234 |
| *ecpA* | standard | 1.36 | 7 | 5.15 | *yijD* | 302.74 | 2.19 | 0.35 | 8.95E-11 | 1.45E-09 |
| *rzoQ* | standard | 0.68 | 3.5 | 5.15 | *gadW* | 98.62 | 2.19 | 0.49 | 1.11E-06 | 1.08E-05 |
| *tfaP* | standard | 0.68 | 3.5 | 5.15 | *zraR* | 36.16 | 2.19 | 0.66 | 1.39E-04 | 8.84E-04 |
| *thiQ* | standard | 0.68 | 3.5 | 5.15 | *psiF* | 55.56 | 2.17 | 0.56 | 1.53E-05 | 1.19E-04 |
| *ybbA* | standard | 2.04 | 10.51 | 5.15 | *mgrR* | 14.06 | 2.17 | 1.04 | 0.004721938 | 0.019238208 |
| *ydbH* | standard | 0.68 | 3.5 | 5.15 | *ydgV* | 108.03 | 2.14 | 0.46 | 5.08E-07 | 5.30E-06 |
| *pgpC* | standard | 21.74 | 108.99 | 5.01 | *yohP* | 29.27 | 2.14 | 0.72 | 4.45E-04 | 0.002479686 |
| *yqgF* | standard | 2.72 | 13.57 | 4.99 | *ygiW* | 67.93 | 2.13 | 0.52 | 7.97E-06 | 6.53E-05 |
| *yeaY* | standard | 63.19 | 314.72 | 4.98 | *pgaD* | 9.14 | 2.13 | 1.38 | 0.012332021 | 0.042160671 |
| *waaO* | standard | 7.47 | 37.21 | 4.98 | *ytfK* | 199.14 | 2.12 | 0.38 | 4.18E-09 | 5.74E-08 |
| *ydcL* | standard | 16.99 | 84.04 | 4.95 | *sixA* | 116.56 | 2.12 | 0.44 | 3.09E-07 | 3.35E-06 |

| | | | | |
|---|---|---|---|---|
| *csdE* | standard | 6.11 | 30.2 | 4.94 |
| *yeeY* | standard | 2.04 | 10.07 | 4.94 |
| *yggC* | standard | 4.08 | 20.13 | 4.94 |
| *waaG* | standard | 2.04 | 10.07 | 4.94 |
| *rseB* | standard | 163.74 | 808.02 | 4.93 |
| *ykgS* | small | 11.55 | 56.9 | 4.93 |
| *yfhH* | standard | 16.31 | 80.1 | 4.91 |
| *yodE* | small | 40.77 | 200.03 | 4.91 |
| *artQ* | standard | 1.36 | 6.57 | 4.83 |
| *xynR* | standard | 1.36 | 6.57 | 4.83 |
| *yfjF* | standard | 5.44 | 26.26 | 4.83 |
| *waaP* | standard | 10.19 | 49.02 | 4.81 |
| *opgB* | standard | 25.14 | 119.5 | 4.75 |
| *rmf* | standard | 499.37 | 2361.46 | 4.73 |
| *yafS* | standard | 4.08 | 19.26 | 4.72 |
| *yegS* | standard | 8.15 | 38.52 | 4.72 |
| *clcB* | standard | 4.76 | 22.32 | 4.69 |
| *yhdV* | standard | 25.82 | 120.81 | 4.68 |
| *yriB* | small | 18.34 | 85.79 | 4.68 |
| *tcyJ* | standard | 86.97 | 399.19 | 4.59 |
| *gsiA* | standard | 7.47 | 34.14 | 4.57 |
| *amyA* | standard | 15.63 | 71.35 | 4.57 |
| *hscA* | standard | 41.44 | 187.78 | 4.53 |
| *cysI* | standard | 21.74 | 98.05 | 4.51 |
| *fepE* | standard | 0.68 | 3.06 | 4.51 |
| *folK* | standard | 0.68 | 3.06 | 4.51 |
| *hisA* | standard | 0.68 | 3.06 | 4.51 |
| *hisM* | standard | 0.68 | 3.06 | 4.51 |
| *hyaF* | standard | 0.68 | 3.06 | 4.51 |
| *insD1* | standard | 0.68 | 3.06 | 4.51 |
| *moaE* | standard | 0.68 | 3.06 | 4.51 |
| *pdxI* | standard | 3.4 | 15.32 | 4.51 |
| *sapF* | standard | 0.68 | 3.06 | 4.51 |
| *waaF* | standard | 1.36 | 6.13 | 4.51 |
| *yafL* | standard | 0.68 | 3.06 | 4.51 |
| *ycbV* | standard | 0.68 | 3.06 | 4.51 |
| *ydaG* | small | 1.36 | 6.13 | 4.51 |
| *yfdX* | standard | 1.36 | 6.13 | 4.51 |
| *aat* | standard | 61.83 | 278.82 | 4.51 |
| *livJ* | standard | 2.04 | 9.19 | 4.51 |
| *paaK* | standard | 2.04 | 9.19 | 4.51 |
| *trmD* | standard | 148.11 | 664.89 | 4.49 |

| | | | | | |
|---|---|---|---|---|---|
| *yafW* | 13.72 | 2.1 | 1.05 | 0.005813937 | 0.022796754 |
| *yqhI* | 17.34 | 2.1 | 0.92 | 0.003119101 | 0.013443068 |
| *ybaT* | 19.7 | 2.09 | 0.86 | 0.002281827 | 0.010453641 |
| *opgB* | 81.48 | 2.08 | 0.49 | 3.69E-06 | 3.28E-05 |
| *glnX* | 1680.13 | 2.07 | 0.34 | 1.46E-10 | 2.27E-09 |
| *glnV* | 1680.13 | 2.07 | 0.34 | 1.46E-10 | 2.27E-09 |
| *holE* | 345.7 | 2.07 | 0.35 | 5.26E-10 | 7.79E-09 |
| *yhcN* | 112.26 | 2.07 | 0.44 | 5.33E-07 | 5.52E-06 |
| *glcD* | 12.51 | 2.07 | 1.11 | 0.007735031 | 0.028847334 |
| *yeeW* | 19.64 | 2.06 | 0.86 | 0.002416881 | 0.010944167 |
| *yciY* | 33.11 | 2.06 | 0.68 | 3.66E-04 | 0.002099702 |
| *gadA* | 55 | 2.06 | 0.6 | 1.09E-04 | 7.20E-04 |
| *kefB* | 29.32 | 2.06 | 0.71 | 6.02E-04 | 0.003210015 |
| *sibC* | 62.55 | 2.06 | 0.53 | 1.98E-05 | 1.51E-04 |
| *corA* | 126.56 | 2.03 | 0.44 | 7.50E-07 | 7.55E-06 |
| *ynfC* | 26.23 | 2.02 | 0.74 | 0.001064516 | 0.005332848 |
| *yoaC* | 21.34 | 2.01 | 0.82 | 0.002245505 | 0.010302324 |
| *leuP* | 370.92 | 1.99 | 0.44 | 1.03E-06 | 1.01E-05 |
| *proP* | 542.82 | 1.97 | 0.33 | 3.48E-10 | 5.29E-09 |
| *ecpA* | 34.9 | 1.97 | 0.66 | 4.90E-04 | 0.002672328 |
| *arnE* | 10.7 | 1.97 | 1.18 | 0.012091365 | 0.04162143 |
| *hscA* | 342.73 | 1.96 | 0.4 | 1.70E-07 | 1.92E-06 |
| *rpoE* | 319.84 | 1.95 | 0.35 | 5.02E-09 | 6.80E-08 |
| *amiC* | 208.92 | 1.95 | 0.38 | 5.25E-08 | 6.52E-07 |
| *ugpE* | 16.7 | 1.94 | 0.94 | 0.005879849 | 0.023026342 |
| *ymiA* | 23.96 | 1.91 | 0.78 | 0.002399407 | 0.010880791 |
| *yahO* | 74.6 | 1.91 | 0.51 | 3.29E-05 | 2.40E-04 |
| *phoU* | 114.77 | 1.91 | 0.44 | 2.77E-06 | 2.50E-05 |
| *ykfG* | 11.5 | 1.9 | 1.13 | 0.012368798 | 0.042181183 |
| *pstB* | 120.34 | 1.9 | 0.43 | 2.03E-06 | 1.86E-05 |
| *basS* | 32.19 | 1.89 | 0.68 | 9.49E-04 | 0.004844557 |
| *slyA* | 260.77 | 1.89 | 0.36 | 3.52E-08 | 4.47E-07 |
| *yhiD* | 15.9 | 1.89 | 0.94 | 0.007083942 | 0.026705609 |
| *kefG* | 73.43 | 1.87 | 0.51 | 4.47E-05 | 3.18E-04 |
| *yobH* | 41.89 | 1.86 | 0.61 | 4.13E-04 | 0.002321485 |
| *selC* | 48.44 | 1.85 | 0.58 | 2.89E-04 | 0.001710781 |
| *fdx* | 172.13 | 1.85 | 0.44 | 5.09E-06 | 4.40E-05 |
| *kbp* | 41.95 | 1.84 | 0.61 | 4.87E-04 | 0.002664475 |
| *gadF* | 49.84 | 1.84 | 0.63 | 6.72E-04 | 0.003539518 |
| *bglA* | 277.97 | 1.84 | 0.38 | 2.40E-07 | 2.66E-06 |
| *yodD* | 16.48 | 1.83 | 0.92 | 0.007466941 | 0.027947439 |
| *hdeB* | 216.8 | 1.83 | 0.43 | 3.89E-06 | 3.44E-05 |

| | | | | | | | | | | |
|---|---|---|---|---|---|---|---|---|---|---|
| *mgtS* | small | 2234.61 | 9955.35 | 4.46 | *osmC* | 176.67 | 1.83 | 0.41 | 1.61E-06 | 1.52E-05 |
| *bfr* | standard | 13.59 | 60.4 | 4.45 | *yhbE* | 148.23 | 1.81 | 0.43 | 5.00E-06 | 4.33E-05 |
| *osmF* | standard | 4.76 | 21.01 | 4.42 | *mlaD* | 217.43 | 1.8 | 0.37 | 2.49E-07 | 2.73E-06 |
| *tomB* | standard | 27.86 | 122.12 | 4.38 | *cspB* | 69.69 | 1.79 | 0.51 | 9.40E-05 | 6.26E-04 |
| *dkgB* | standard | 11.55 | 50.34 | 4.36 | *serW* | 53.51 | 1.79 | 0.58 | 3.98E-04 | 0.002250482 |
| *bcsG* | standard | 2.72 | 11.82 | 4.35 | *serX* | 53.51 | 1.79 | 0.58 | 3.98E-04 | 0.002250482 |
| *ydeP* | standard | 2.72 | 11.82 | 4.35 | *ortT* | 31.77 | 1.79 | 0.74 | 0.002848726 | 0.012607727 |
| *ycgZ* | standard | 20.38 | 88.42 | 4.34 | *yfeK* | 18.38 | 1.78 | 0.86 | 0.006865127 | 0.025974585 |
| *glk* | standard | 43.48 | 188.22 | 4.33 | *yafX* | 24.7 | 1.78 | 0.76 | 0.00357779 | 0.015231165 |
| *cynR* | standard | 22.42 | 96.73 | 4.31 | *basR* | 41.62 | 1.76 | 0.61 | 7.62E-04 | 0.003938452 |
| *fxsA* | standard | 8.83 | 38.08 | 4.31 | *yabI* | 44.88 | 1.76 | 0.58 | 5.35E-04 | 0.002873773 |
| *yhfG* | standard | 12.91 | 55.15 | 4.27 | *gadE* | 27.23 | 1.76 | 0.76 | 0.004028227 | 0.0168282 |
| *amiB* | standard | 9.51 | 40.27 | 4.23 | *azuC* | 71.86 | 1.76 | 0.62 | 9.66E-04 | 0.004912092 |
| *ldtC* | standard | 16.99 | 71.35 | 4.2 | *mdaB* | 102.97 | 1.75 | 0.45 | 2.21E-05 | 1.67E-04 |
| *rstA* | standard | 64.54 | 270.51 | 4.19 | *ynfS* | 90.26 | 1.75 | 0.46 | 3.43E-05 | 2.50E-04 |
| *ahr* | standard | 9.51 | 39.83 | 4.19 | *ydhR* | 93 | 1.73 | 0.46 | 4.57E-05 | 3.23E-04 |
| *fbaB* | standard | 9.51 | 39.83 | 4.19 | *rcnA* | 13.84 | 1.73 | 0.99 | 0.01356292 | 0.045147211 |
| *bcsF* | standard | 1.36 | 5.69 | 4.19 | *ysgD* | 402.58 | 1.72 | 0.35 | 2.73E-07 | 2.98E-06 |
| *chbG* | standard | 5.44 | 22.76 | 4.19 | *shoB* | 19.74 | 1.72 | 0.83 | 0.007317217 | 0.027419846 |
| *fadE* | standard | 2.72 | 11.38 | 4.19 | *ygdI* | 40.7 | 1.71 | 0.6 | 9.86E-04 | 0.004994682 |
| *ycgB* | standard | 7.47 | 31.08 | 4.16 | *mlaB* | 155.62 | 1.71 | 0.4 | 4.03E-06 | 3.53E-05 |
| *gsiB* | standard | 3.4 | 14.01 | 4.12 | *rem* | 13.58 | 1.7 | 1 | 0.01516317 | 0.049319708 |
| *pxpA* | standard | 3.4 | 14.01 | 4.12 | *ybaP* | 80.94 | 1.7 | 0.47 | 8.16E-05 | 5.53E-04 |
| *wecB* | standard | 2.04 | 8.32 | 4.08 | *mltF* | 22.43 | 1.68 | 0.78 | 0.006371506 | 0.02459144 |
| *ydeM* | standard | 2.04 | 8.32 | 4.08 | *yciX* | 63.21 | 1.67 | 0.51 | 2.67E-04 | 0.001591423 |
| *nimR* | standard | 10.87 | 44.21 | 4.07 | *leuT* | 333.1 | 1.66 | 0.42 | 1.71E-05 | 1.31E-04 |
| *psiF* | standard | 8.83 | 35.89 | 4.06 | *leuV* | 333.1 | 1.66 | 0.42 | 1.71E-05 | 1.31E-04 |
| *ynjB* | standard | 9.51 | 38.08 | 4 | *leuQ* | 333.1 | 1.66 | 0.42 | 1.71E-05 | 1.31E-04 |
| *yiaG* | standard | 10.19 | 40.71 | 3.99 | *trpT* | 53.05 | 1.66 | 0.55 | 6.59E-04 | 0.0034826 |
| *uhpB* | standard | 12.91 | 51.21 | 3.97 | *hdeA* | 666.9 | 1.65 | 0.37 | 2.31E-06 | 2.11E-05 |
| *ecpR* | standard | 5.44 | 21.45 | 3.95 | *elaB* | 154.33 | 1.65 | 0.4 | 1.08E-05 | 8.61E-05 |
| *mntP* | standard | 23.1 | 91.04 | 3.94 | *letA* | 25.84 | 1.64 | 0.73 | 0.005293456 | 0.021180592 |
| *msyB* | standard | 78.13 | 307.71 | 3.94 | *mltD* | 413.19 | 1.64 | 0.34 | 4.54E-07 | 4.79E-06 |
| *dgcQ* | standard | 6.11 | 24.07 | 3.94 | *glmY* | 73.6 | 1.61 | 0.49 | 2.56E-04 | 0.001541773 |
| *ydbJ* | standard | 99.87 | 392.19 | 3.93 | *maeA* | 355.15 | 1.61 | 0.37 | 3.00E-06 | 2.70E-05 |
| *alx* | standard | 7.47 | 29.33 | 3.92 | *ykfF* | 34.95 | 1.6 | 0.64 | 0.002915101 | 0.012813055 |
| *yraQ* | standard | 38.05 | 148.82 | 3.91 | *tatD* | 80.68 | 1.6 | 0.48 | 2.05E-04 | 0.001258503 |
| *ldtA* | standard | 29.89 | 116.87 | 3.91 | *ynaI* | 32.23 | 1.6 | 0.67 | 0.003919051 | 0.016504319 |
| *rhoL* | small | 10.87 | 42.46 | 3.91 | *rseA* | 791.21 | 1.59 | 0.33 | 2.88E-07 | 3.13E-06 |
| *ygiW* | standard | 6.79 | 26.26 | 3.87 | *glgS* | 35.1 | 1.59 | 0.66 | 0.003814277 | 0.016106441 |
| *gudX* | standard | 0.68 | 2.63 | 3.87 | *gcvB* | 3584.49 | 1.59 | 0.32 | 1.68E-07 | 1.91E-06 |
| *ibsB* | small | 2.72 | 10.51 | 3.87 | *msyB* | 63.26 | 1.58 | 0.52 | 6.14E-04 | 0.003265833 |

| | | | | | | | | | | |
|---|---|---|---|---|---|---|---|---|---|---|
| *malX* | standard | 0.68 | 2.63 | 3.87 | *valW* | 46.96 | 1.58 | 0.59 | 0.001741963 | 0.008208737 |
| *mdtD* | standard | 0.68 | 2.63 | 3.87 | *aspU* | 650.02 | 1.58 | 0.34 | 1.23E-06 | 1.18E-05 |
| *moa D* | standard | 1.36 | 5.25 | 3.87 | *aspV* | 650.02 | 1.58 | 0.34 | 1.23E-06 | 1.18E-05 |
| *msrP* | standard | 1.36 | 5.25 | 3.87 | *aspT* | 650.02 | 1.58 | 0.34 | 1.23E-06 | 1.18E-05 |
| *phnN* | standard | 0.68 | 2.63 | 3.87 | *omp X* | 4546.19 | 1.57 | 0.32 | 2.02E-07 | 2.27E-06 |
| *pphA* | standard | 1.36 | 5.25 | 3.87 | *pmrR* | 156.83 | 1.57 | 0.39 | 1.97E-05 | 1.50E-04 |
| *recO* | standard | 1.36 | 5.25 | 3.87 | *tyrV* | 115.05 | 1.56 | 0.44 | 1.12E-04 | 7.33E-04 |
| *rseC* | standard | 0.68 | 2.63 | 3.87 | *tyrT* | 115.05 | 1.56 | 0.44 | 1.12E-04 | 7.33E-04 |
| *sxy* | standard | 1.36 | 5.25 | 3.87 | *mlaC* | 223.38 | 1.56 | 0.37 | 6.04E-06 | 5.08E-05 |
| *trmO* | standard | 2.72 | 10.51 | 3.87 | *emrD* | 40.84 | 1.55 | 0.6 | 0.0025991 | 0.011651268 |
| *ydhP* | standard | 18.34 | 70.91 | 3.87 | *ydhP* | 31.46 | 1.54 | 0.67 | 0.005090086 | 0.020524328 |
| *yfbN* | standard | 0.68 | 2.63 | 3.87 | *dinJ* | 131.95 | 1.54 | 0.41 | 5.98E-05 | 4.14E-04 |
| *yjcO* | standard | 0.68 | 2.63 | 3.87 | *iscR* | 308.39 | 1.54 | 0.51 | 7.04E-04 | 0.003669559 |
| *pyrR* | standard | 2.04 | 7.88 | 3.87 | *sra* | 525.89 | 1.53 | 0.34 | 1.75E-06 | 1.64E-05 |
| *ybjC* | standard | 8.15 | 31.52 | 3.87 | *yoeB* | 53.81 | 1.52 | 0.54 | 0.001361759 | 0.006609666 |
| *paaJ* | standard | 2.04 | 7.88 | 3.87 | *gltS* | 37.19 | 1.5 | 0.62 | 0.004079394 | 0.017019232 |
| *tmcA* | standard | 2.04 | 7.88 | 3.87 | *osmY* | 248.23 | 1.5 | 0.4 | 4.80E-05 | 3.38E-04 |
| *yceM* | standard | 2.04 | 7.88 | 3.87 | *ugpC* | 28.39 | 1.46 | 0.69 | 0.009072358 | 0.032932027 |
| *ydeI* | standard | 2.04 | 7.88 | 3.87 | *soxS* | 136.64 | 1.45 | 0.42 | 1.65E-04 | 0.001041569 |
| *yghB* | standard | 42.8 | 164.58 | 3.85 | *yejG* | 207.82 | 1.45 | 0.4 | 8.98E-05 | 6.04E-04 |
| *ymiA* | small | 138.6 | 530.51 | 3.83 | *ykiD* | 30.44 | 1.44 | 0.66 | 0.008017026 | 0.029651623 |
| *yibF* | standard | 7.47 | 28.45 | 3.81 | *cof* | 36.87 | 1.43 | 0.61 | 0.005648098 | 0.022314267 |
| *yphH* | standard | 11.55 | 43.77 | 3.79 | *rcnR* | 28.58 | 1.43 | 0.68 | 0.00981519 | 0.035219873 |
| *curA* | standard | 22.42 | 84.92 | 3.79 | *rpoS* | 518.95 | 1.41 | 0.33 | 5.85E-06 | 4.95E-05 |
| *npr* | standard | 15.63 | 59.09 | 3.78 | *ryfA* | 56.68 | 1.41 | 0.52 | 0.002227727 | 0.010235766 |
| *ytfF* | standard | 9.51 | 35.89 | 3.77 | *yqhA* | 154.26 | 1.39 | 0.39 | 1.37E-04 | 8.75E-04 |
| *cysE* | standard | 47.56 | 178.59 | 3.76 | *psiE* | 28.88 | 1.39 | 0.68 | 0.011626599 | 0.040374065 |
| *yedA* | standard | 7.47 | 28.01 | 3.75 | *glyU* | 39.24 | 1.38 | 0.6 | 0.006460553 | 0.024823494 |
| *osm Y* | standard | 40.09 | 150.14 | 3.75 | *erpA* | 165.91 | 1.35 | 0.47 | 0.001365789 | 0.006615408 |
| *gad W* | standard | 63.19 | 236.37 | 3.74 | *yeaQ* | 65.54 | 1.35 | 0.5 | 0.002133684 | 0.009848486 |
| *dacC* | standard | 55.71 | 208.35 | 3.74 | *dkgB* | 31.73 | 1.35 | 0.65 | 0.011599563 | 0.040332427 |
| *ytiA* | standard | 3.4 | 12.69 | 3.74 | *yceK* | 99.9 | 1.33 | 0.46 | 0.00134666 | 0.006563393 |
| *dnaC* | standard | 9.51 | 35.45 | 3.73 | *adhP* | 30.59 | 1.33 | 0.65 | 0.012832445 | 0.043221442 |
| *phoP* | standard | 233.04 | 864.48 | 3.71 | *cspF* | 36.22 | 1.32 | 0.61 | 0.009769932 | 0.035138064 |
| *hybD* | standard | 2.72 | 10.07 | 3.7 | *cydH* | 77.27 | 1.32 | 0.47 | 0.001728634 | 0.008158215 |
| *tatB* | standard | 2.72 | 10.07 | 3.7 | *yefM* | 123.88 | 1.32 | 0.41 | 5.11E-04 | 0.002764991 |
| *yebE* | standard | 12.91 | 47.27 | 3.66 | *yedR* | 30.32 | 1.31 | 0.65 | 0.014078261 | 0.046418208 |
| *nagA* | standard | 14.95 | 54.71 | 3.66 | *degP* | 308.32 | 1.3 | 0.36 | 9.23E-05 | 6.19E-04 |
| *hem F* | standard | 6.11 | 22.32 | 3.65 | *yqeF* | 37.96 | 1.29 | 0.61 | 0.011566698 | 0.040303115 |
| *dadX* | standard | 2.04 | 7.44 | 3.65 | *yebV* | 154.21 | 1.29 | 0.41 | 5.61E-04 | 0.003003626 |
| *ggt* | standard | 4.08 | 14.88 | 3.65 | *iscS* | 431.78 | 1.29 | 0.45 | 0.001488886 | 0.007167266 |
| *yoeB* | standard | 2.04 | 7.44 | 3.65 | *zapC* | 130.31 | 1.28 | 0.4 | 5.74E-04 | 0.003072022 |

| | | | | | | | | | | |
|---|---|---|---|---|---|---|---|---|---|---|
| *letA* | standard | 52.99 | 193.03 | 3.64 | *yncL* | 120.28 | 1.28 | 0.42 | 9.49E-04 | 0.004844557 |
| *ytjA* | standard | 47.56 | 172.46 | 3.63 | *ohsC* | 109.96 | 1.27 | 0.42 | 0.001015212 | 0.005098873 |
| *uvrY* | standard | 27.18 | 98.05 | 3.61 | *yrbN* | 225.13 | 1.27 | 0.37 | 2.38E-04 | 0.001449485 |
| *yhjD* | standard | 8.15 | 29.33 | 3.6 | *mdtK* | 62.12 | 1.27 | 0.5 | 0.00443095 | 0.018242686 |
| *tehA* | standard | 18.34 | 65.66 | 3.58 | *yghE* | 41.85 | 1.27 | 0.58 | 0.010335855 | 0.036693964 |
| *adhP* | standard | 5.44 | 19.26 | 3.54 | *rsxA* | 53.19 | 1.26 | 0.54 | 0.006530064 | 0.024978693 |
| *cmo B* | standard | 1.36 | 4.81 | 3.54 | *sfsA* | 54.6 | 1.25 | 0.52 | 0.006450973 | 0.024823494 |
| *pncA* | standard | 1.36 | 4.81 | 3.54 | *ytiB* | 51.03 | 1.25 | 0.54 | 0.007595897 | 0.028362246 |
| *uhpC* | standard | 1.36 | 4.81 | 3.54 | *tusA* | 75.69 | 1.25 | 0.47 | 0.003008532 | 0.013056443 |
| *insA4* | standard | 4.08 | 14.44 | 3.54 | *rplS* | 741.32 | 1.24 | 0.34 | 1.17E-04 | 7.62E-04 |
| *pmrD* | standard | 158.3 | 556.33 | 3.51 | *rnlB* | 45.16 | 1.23 | 0.56 | 0.010036429 | 0.035808421 |
| *slp* | standard | 19.02 | 66.53 | 3.5 | *blr* | 68.02 | 1.23 | 0.49 | 0.004412675 | 0.018201483 |
| *yfjD* | standard | 4.76 | 16.63 | 3.5 | *bfr* | 69.08 | 1.23 | 0.49 | 0.004582157 | 0.018766453 |
| *talA* | standard | 13.59 | 47.27 | 3.48 | *ibpA* | 46.01 | 1.2 | 0.55 | 0.011140268 | 0.038903904 |
| *yhbO* | standard | 6.79 | 23.64 | 3.48 | *tatE* | 68.34 | 1.19 | 0.48 | 0.005545779 | 0.022021246 |
| *yidZ* | standard | 29.89 | 103.74 | 3.47 | *iscX* | 499.84 | 1.18 | 0.4 | 0.001193108 | 0.005907016 |
| *yecN* | standard | 42.8 | 147.95 | 3.46 | *obgE* | 194.26 | 1.17 | 0.39 | 9.99E-04 | 0.005046111 |
| *mep M* | standard | 15.63 | 53.84 | 3.45 | *tolC* | 692.49 | 1.17 | 0.33 | 2.01E-04 | 0.001238049 |
| *ecnA* | small | 2.04 | 7 | 3.44 | *iscU* | 305.42 | 1.16 | 0.43 | 0.002823147 | 0.012529967 |
| *fecC* | standard | 2.04 | 7 | 3.44 | *frmR* | 87.67 | 1.15 | 0.45 | 0.004308386 | 0.01787923 |
| *phnD* | standard | 4.08 | 14.01 | 3.44 | *rseC* | 118.42 | 1.15 | 0.41 | 0.002334944 | 0.010619245 |
| *ycgY* | standard | 2.04 | 7 | 3.44 | *eco* | 76.98 | 1.15 | 0.47 | 0.006182743 | 0.024121949 |
| *ychO* | standard | 4.08 | 14.01 | 3.44 | *atpI* | 308.54 | 1.14 | 0.35 | 4.75E-04 | 0.002606775 |
| *ynjD* | standard | 2.04 | 7 | 3.44 | *rpoZ* | 113.46 | 1.14 | 0.44 | 0.004444168 | 0.018273064 |
| *thiL* | standard | 8.83 | 30.2 | 3.42 | *roxA* | 178.9 | 1.14 | 0.38 | 0.001264669 | 0.006212163 |
| *ybiR* | standard | 8.83 | 30.2 | 3.42 | *lpxT* | 53.03 | 1.14 | 0.53 | 0.013268514 | 0.04421425 |
| *ytjB* | standard | 69.3 | 236.8 | 3.42 | *hdeD* | 64.47 | 1.13 | 0.53 | 0.012893055 | 0.043351172 |
| *metI* | standard | 4.76 | 16.2 | 3.41 | *rcnB* | 64.97 | 1.13 | 0.49 | 0.008988767 | 0.032666496 |
| *ybjS* | standard | 93.76 | 318.66 | 3.4 | *yffB* | 84.37 | 1.12 | 0.46 | 0.006420724 | 0.024741928 |
| *dnaK* | standard | 14.95 | 50.77 | 3.4 | *cdsA* | 70.95 | 1.1 | 0.49 | 0.011431427 | 0.03987618 |
| *rcsC* | standard | 12.91 | 43.77 | 3.39 | *iscA* | 310.37 | 1.1 | 0.42 | 0.004388433 | 0.018139242 |
| *phnO* | standard | 10.87 | 36.77 | 3.38 | *osmB* | 100.71 | 1.09 | 0.43 | 0.005613039 | 0.022203793 |
| *pspD* | standard | 40.77 | 137.44 | 3.37 | *lysU* | 87.48 | 1.08 | 0.46 | 0.008192364 | 0.030264355 |
| *hokD* | standard | 38.73 | 130.44 | 3.37 | *ecnB* | 69.09 | 1.08 | 0.48 | 0.010833276 | 0.038044129 |
| *inaA* | standard | 15.63 | 52.53 | 3.36 | *pepB* | 120.64 | 1.08 | 0.42 | 0.004873856 | 0.019754267 |
| *pabB* | standard | 39.41 | 131.75 | 3.34 | *rseB* | 238.27 | 1.07 | 0.36 | 0.001374135 | 0.006645545 |
| *yfdC* | standard | 19.7 | 65.66 | 3.33 | *nadD* | 71.06 | 1.05 | 0.47 | 0.012553295 | 0.042741306 |
| *fumE* | standard | 25.82 | 85.79 | 3.32 | *ivy* | 123.06 | 1.05 | 0.41 | 0.005102016 | 0.020545956 |
| *aroC* | standard | 4.76 | 15.76 | 3.31 | *bax* | 212.78 | 1.04 | 0.36 | 0.002015903 | 0.009331005 |
| *wecF* | standard | 16.99 | 56.03 | 3.3 | *bssS* | 188.72 | 1.03 | 0.37 | 0.002690689 | 0.011976054 |
| *yadX* | small | 23.78 | 78.35 | 3.29 | *bolA* | 157.88 | 1.02 | 0.4 | 0.00569691 | 0.022448028 |
| *degP* | standard | 21.06 | 69.16 | 3.28 | *ydgK* | 78.89 | 1.01 | 0.45 | 0.012656898 | 0.042898198 |

| | | | | |
|---|---|---|---|---|
| *ydhI* | standard | 14.27 | 46.84 | 3.28 |
| *ymdF* | standard | 14.27 | 46.84 | 3.28 |
| *hisQ* | standard | 16.99 | 55.59 | 3.27 |
| *yodC* | standard | 161.7 | 524.82 | 3.25 |
| *abpA* | standard | 24.46 | 79.23 | 3.24 |
| *ybjG* | standard | 303.02 | 980.92 | 3.24 |
| *yeaG* | standard | 29.89 | 96.73 | 3.24 |
| *mrcB* | standard | 48.24 | 155.83 | 3.23 |
| *mrdB* | standard | 1.36 | 4.38 | 3.22 |
| *acpS* | standard | 0.68 | 2.19 | 3.22 |
| *arnF* | standard | 0.68 | 2.19 | 3.22 |
| *essQ* | standard | 0.68 | 2.19 | 3.22 |
| *hcr* | standard | 0.68 | 2.19 | 3.22 |
| *ilvH* | standard | 0.68 | 2.19 | 3.22 |
| *insB2* | standard | 0.68 | 2.19 | 3.22 |
| *insB3* | standard | 0.68 | 2.19 | 3.22 |
| *kduI* | standard | 0.68 | 2.19 | 3.22 |
| *lpxK* | standard | 2.04 | 6.57 | 3.22 |
| *narJ* | standard | 0.68 | 2.19 | 3.22 |
| *sufD* | standard | 0.68 | 2.19 | 3.22 |
| *treA* | standard | 0.68 | 2.19 | 3.22 |
| *yaaI* | standard | 1.36 | 4.38 | 3.22 |
| *ydhT* | standard | 1.36 | 4.38 | 3.22 |
| *ydhU* | standard | 2.04 | 6.57 | 3.22 |
| *ynaK* | standard | 0.68 | 2.19 | 3.22 |
| *ynbA* | standard | 0.68 | 2.19 | 3.22 |
| *ypdC* | standard | 0.68 | 2.19 | 3.22 |
| *yqeC* | standard | 1.36 | 4.38 | 3.22 |
| *yqiA* | standard | 16.99 | 54.28 | 3.2 |
| *ybgA* | standard | 9.51 | 30.2 | 3.18 |
| *ydgJ* | standard | 108.03 | 342.73 | 3.17 |
| *ynfC* | standard | 8.83 | 28.01 | 3.17 |
| *yhbQ* | standard | 74.74 | 236.8 | 3.17 |
| *metL* | standard | 10.87 | 34.14 | 3.14 |
| *eam A* | standard | 14.27 | 44.65 | 3.13 |
| *dusC* | standard | 9.51 | 29.76 | 3.13 |
| *relE* | standard | 4.76 | 14.88 | 3.13 |
| *ygdR* | standard | 307.1 | 952.03 | 3.1 |
| *yfeY* | standard | 57.07 | 176.84 | 3.1 |
| *dnaG* | standard | 17.66 | 54.71 | 3.1 |
| *muk E* | standard | 35.33 | 108.99 | 3.08 |
| *slt* | standard | 15.63 | 48.15 | 3.08 |

| | | | | | |
|---|---|---|---|---|---|
| *lpxB* | 94.1 | 1 | 0.44 | 0.011029168 | 0.038645314 |
| *deaD* | 606.52 | 0.99 | 0.33 | 0.001428271 | 0.006886069 |
| *frmA* | 117.64 | 0.96 | 0.41 | 0.009881474 | 0.03533615 |
| *ybiE* | 142.68 | 0.96 | 0.39 | 0.008000962 | 0.029627231 |
| *yrbL* | 258.5 | 0.94 | 0.35 | 0.004091926 | 0.017048785 |
| *pheM* | 167.34 | 0.93 | 0.38 | 0.008333725 | 0.030642886 |
| *ybhL* | 238.82 | 0.91 | 0.38 | 0.008835762 | 0.032260328 |
| *hspQ* | 188.37 | 0.87 | 0.38 | 0.012704292 | 0.042928435 |
| *ydiH* | 403.56 | 0.86 | 0.33 | 0.005399113 | 0.021548245 |
| *cpxP* | 205.41 | 0.86 | 0.37 | 0.011798686 | 0.040793466 |
| *thrW* | 199.73 | 0.83 | 0.36 | 0.013973818 | 0.046178214 |
| *nlpD* | 735.03 | 0.77 | 0.31 | 0.008684907 | 0.031820928 |
| *csrB* | 2954.91 | 0.68 | 0.29 | 0.014267278 | 0.046991908 |

| | | | | |
|---|---|---|---|---|
| *ybiJ* | standard | 6.11 | 18.82 | 3.08 |
| *moaA* | standard | 26.5 | 81.41 | 3.07 |
| *zraR* | standard | 23.1 | 70.91 | 3.07 |
| *insH5* | standard | 32.61 | 99.8 | 3.06 |
| *cvpA* | standard | 17.66 | 53.4 | 3.02 |
| *tcdA* | standard | 30.57 | 91.92 | 3.01 |
| *ycjX* | standard | 8.15 | 24.51 | 3.01 |
| *yigL* | standard | 4.08 | 12.26 | 3.01 |
| *ymiC* | small | 282.63 | 839.53 | 3.00 |

**Table S5. Verification of protein expression from the genomic locus for the 17 candidates identified in this study using Ribo-RET.**

| Small Protein | Small protein expression by genomic tagging first reported in | Media used |
|---|---|---|
| MgtS | (*21*) | LB |
| | (*34*) | Supplemented N-minimal medium with no added $Mg^{2+}$ |
| MgrB | (*21*) | LB |
| PmrR | (*36*) | N-minimal medium, pH 7.7 with 10 μM $Mg^{2+}$ |
| MgtT | (*41*) | LB |
| YmiC | (*95*) | LB |
| YmiA | (*21*) | LB |
| YoaI | [This Study] | Supplemented minimal A medium with 10 mM, 1 mM and no added $Mg^{2+}$ |
| YobF | (*21*) | LB |
| | (*22*) | LB + heat shock (45°C) |
| YddY | (*95*) | LB |
| YriB | (*41*) | LB |
| YadX | (*41*) | LB |
| YkgS | (*95*) | LB |
| YriA | (*41*) | LB |
| YdgU | (*21*) | LB |
| YqhI | (*95*) | LB |
| YadW | (*95*) | LB |
| DinQ | (*50*) | LB |

**Table S6. Analysis of transcriptional regulation of low $Mg^{2+}$ stress-induced small proteins.** Putative regulatory regions corresponding to the stress-induced small proteins and their dependence on PhoQ-PhoP signaling system.

| Small protein | Putative region(s) of interest | Length of region (bp) | Genomic location in *E. coli* MG1655 | PhoQP-dependence | Low $Mg^{2+}$ stress-specific expression reported in |
|---|---|---|---|---|---|
| MgrB | $P_{mgrB}$ | 500 | 1908767 - 1909266 | Yes | (*30*) |
| MgtS, MgtT | $P_{mgtS\text{-}mgtT}$ | 405 | 1622241 - 1622645 | Yes | (*34*), This study |
| ProP, PmrR | $P_{proP\text{-}pmrR}$ | 500 | 4330002 - 4330501 | No | This study |
| PmrR | $P_{pmrR}$ | 500 | 4331616 - 4332115 | No | This study |
| YmiA, YmiC | $P_{ymiA\text{-}yciX\text{-}ymiC}$ | 500 | 1334648 - 1335147 | Yes | This study |
| YmiC | $P_{ymiC}$ | 272 | 1335300 - 1335571 | No | This study |
| YoaI | $P_{yoaI}$ | 500 | 1874183 - 1874682 | No | This study |
| YobF | $P_{yobF}$ | 500 | 1907592 - 1908091 | No | This study |
| YddY | $P_{yddY}$ | 500 | 1567220 - 1567719 | Yes | This study |
| YriA | $P_{yriA\text{-}yriB}$ | 500 | 3640697 - 3641196 | Yes | This study |
| YriB | | | | | |

| | | | | | |
|---|---|---|---|---|---|
| YadX, YadW | $P_{yadX\text{-}clcA\text{-}yadW}$ | 500 | 174548 - 175047 | Yes | This study |
| YadW | $P_{yadW}$ | 500 | 176052 - 176551 | No | This study |
| YkgS | $P_{ykgS}$ | 230 | 290280 - 290509 | No | This study |
| YdgU | $P_{asr\text{-}ydgU}$ | 400 | 1670976 - 1671375 | Yes | This study |
| | $P_{ydgU}$ | 500 | 1671277 - 1671776 | No | This study |
| YqhI | $P_{yqhI}$ | 500 | 3147741 - 3148240 | No | This study |
| DinQ | $P_{dinQ}$ | 272 | 3647789-3648060 | No | This study |

**Table S7. Localization, Tag, and Membrane Topology Prediction of Small Proteins**

| Small Protein | Localization | Epitope tag | Reference | Prediction of transmembrane helix and orientation | | |
|---|---|---|---|---|---|---|
| | | | | **TMHMM** (*57*) | **TMPred** (*58*) | **Phobius** (*59*) |
| MgtS | Membrane | C-terminal/ FLAG and C-terminal/ GFP | (*21*, *96*) | Yes | Predicted helix: 7-25 (19aa); Inside to Outside: N-terminus facing cytoplasm | Cytoplasmic (1-6aa) / Transmembrane (7-25aa) / Non - cytoplasmic (26-31aa) |
| MgrB | Membrane | C-terminal/ FLAG and N-terminal/ GFP | (*21*, *30*) | Yes | Predicted helix: 6-24 (19aa); Inside to Outside: N-terminus facing cytoplasm | Positively charged or n-region (1-5aa) / Hydrophobic α-helical or h-region (6-17aa)/ Non cytoplasmic (22-47) |
| PmrR | Membrane | C-terminal/ 6XHis | This study | Yes | Predicted helix: 5-27 (23aa); Inside to Outside: N-terminus facing cytoplasm | Cytoplasmic (1-8aa) / transmembrane (9-26 aa) / non cytoplasmic (27-29aa) |
| | | N-terminal/ FLAG-6XHis (*S. enterica* PmrR) | (*36*) | | | |
| MgtT | Cytoplasm | N-terminal/ GFP | This study | No | NA | Non cytoplasmic |
| YmiC | Membrane | N-terminal/ GFP | This study | Yes | Predicted helix: 8-29 (22aa); Inside to Outside: N-terminus facing cytoplasm | Cytoplasmic (1-8aa) / Transmembrane (9-29aa) / Non - cytoplasmic (30-31aa) |
| YmiA | Membrane | N-terminal/ GFP | This study | Yes | Predicted helix: 22-42 (21aa); Inside to Outside: N-terminus facing cytoplasm | Cytoplasmic (1-20aa) / Transmembrane (21-42aa) / Non - cytoplasmic (43-46aa) |

| | | | | | | |
|---|---|---|---|---|---|---|
| YoaI | Membrane | C-terminal/ GFP | This study | Yes | Predicted helix: 10-30 (21aa); Outside to Inside: N-terminus facing periplasm | Non - cytoplasmic (1-5aa) / Transmembrane (6-30aa) / Cytoplasmic (31-34aa) |
| YobF | Membrane | C-terminal/ 6XHis | This study | Yes | Predicted helix: 25-41 (17aa); Outside to Inside: N-terminus facing periplasm | Non cytoplasmic |
| YddY | Cytoplasm | N-terminal/ GFP | This study | No | NA | Non cytoplasmic |
| YriB | Cytoplasm | N-terminal/ GFP | This study | No | NA | Non cytoplasmic |
| YadX | Cytoplasm | N-terminal/ GFP | This study | No | NA | Cytoplasmic |
| YkgS | Cytoplasm | N-terminal/ GFP | This study | No | NA | Cytoplasmic |
| YriA | Cytoplasm | N-terminal/ GFP | This study | No | NA | Non cytoplasmic |
| YdgU | Membrane | N-terminal/ GFP | This study | Yes | Predicted helix: 7-26 (20aa); Inside to Outside: N-terminus facing cytoplasm | Cytoplasmic (1-6aa) / Transmembrane (7-26aa) / Non - cytoplasmic (27aa) |
| YqhI | Cytoplasm | N-terminal/ GFP | This study | No | NA | Non cytoplasmic |
| YadW | Cytoplasm | C-terminal/ GFP | This study | No | NA | Non cytoplasmic |
| DinQ | Membrane | N-terminal FLAG | (*50*) | Yes | Predicted helix: 1-22 (22aa); Outside to Inside: N-terminal facing periplasm | Non - cytoplasmic (1-5aa) / Transmembrane (6-23aa) / Cytoplasmic (24-27aa) |
